# CD19 CAR Engineered T Regulatory Cells Promote Durable Immune Tolerance in Experimental Systemic Lupus Erythematosus

**DOI:** 10.64898/2026.10.05.756761

**Authors:** Travis Drow, Matteo Doglio, Annaiz Grimm, Victoria DeVault-Nelson, Maegan Hoover, Brock McKinney, Subhash K. Tripathi, Jennifer Mellen, Madison Milaszewski, Payam Zarin, Alaina Burgess, Christopher L. Moore, Pierluigi Carulli, Alessia Ugolini, Clara Bercher, Aesha Vakil, Noelle P. Dahl, Sivasankaran Munusamy Ponnan, Neelakshi Mungra, Sheeram Akilesh, Yu Chen, Parker Knebusch, Shaun W. Jackson, Chandra Patel, Tiffany Chen, Alberto Del Rio-Espinola, Chiara Bonini, Tom Wickham, Peter J. Cook, David J. Rawlings

## Abstract

While the immune reset induced by CD19 CAR T therapies has shown striking promise in systemic lupus erythematosus (SLE), the nature and duration of benefit mediated by B cell depletion remains unclear. Loss of tolerance in SLE is driven by extrafollicular and germinal center interactions between dendritic cells and self-reactive T and B cells. We hypothesized that CD19 CAR-engineered regulatory T cells (CAR19 EngTreg), capable of suppressing multiple autoimmune effector cell types, might promote long-term tolerance without producing high-levels of inflammatory mediators. To generate CAR19 EngTreg, we combined CD19 CAR delivery with homology directed repair at the *FOXP3* locus to drive constitutive, stable FOXP3 expression in CD4 T cells. CAR19 EngTreg expressed canonical Treg markers; exhibited minimal inflammatory cytokine production; and blocked human B cell differentiation into antibody secreting plasma cells. In humanized mouse models, transfer of CAR19 EngTreg led to a reduction in B cell numbers with low inflammatory cytokine production relative to CD19 CAR Teff, and reduced tissue inflammation. In a short-term, in vivo imiquimod-induced inflammatory model, murine CAR19 EngTreg reduced skin pathology, spleen size, and inflammatory cytokines. Finally, in two distinct mouse models of lupus, CAR19 EngTreg persisted for >4 months leading to reduced autoimmune T and B effector populations, sustained reduction in multiple autoantibodies and improved renal function and histopathology. Thus, CAR19 EngTreg may offer a promising and durable therapeutic avenue for B cell-mediated autoimmune disorders capable of long-term regulation of both autoreactive B and T cells.

**One Sentence Summary:** CD19 CAR-engineered regulatory T cells provide durable, low-inflammatory control of autoreactive B and T cell responses and improve disease outcomes across preclinical models of lupus.

## INTRODUCTION

Systemic lupus erythematosus (SLE) is a chronic autoimmune disease characterized by a breakdown of central and peripheral tolerance, leading to the production of autoantibodies (*1*) and immune complexes (*2, 3*), that deposit in tissues and drive end-organ injury (*4*). A key feature of SLE pathogenesis is the aberrant expansion of autoreactive B cells. B cells promote disease by functioning as both antibody secreting cells and as antigen-presenting cells (APCs) (*5, 6*) leading to expansion of self-reactive T cells including disease driving T-follicular helper (Tfh) and T peripheral helper (Tph) cells. Localized interactions of self-reactive T and B cells within germinal center (GC) and extrafollicular (EF) compartments, lead to generation of short- and long-lived autoantibody secreting, plasma cells (PCs) and self-reactive memory B cells. Ongoing autoantibody production and iterative interaction of autoreactive Tfh/Tph and memory B cell populations accelerate loss in immune tolerance and feed-forward cytokine dysregulation that reinforce disease activity (*7–9*).

The role of B cells in SLE has motivated multiple therapeutic strategies (*10*). Targeting B cell activating factor (BAFF), which promote B cell survival and activation, has demonstrated partial clinical benefit (*11, 12*). Through the course of differentiation, B cells express varying surface molecules that have emerged as candidate targets for B cell-depleting therapies. CD20, expressed from pre-B cell through memory B cell stages, has been the target of multiple monoclonal antibodies, including rituximab. However, despite depletion of circulating B cells, clinical trials using rituximab failed to demonstrate efficacy in SLE; (*13*) findings consistent with incomplete depletion as supported by biopsy evidence for residual B cells in secondary lymphoid tissue (*14*).

In contrast to CD20 which is minimally expressed on antibody secreting cells, CD19 is expressed from the pro-B cell stage through plasmablast (PB) and early PC development. Recently, autologous T cell therapy using CD19-directed chimeric antigen receptor (CAR19) T cells has shown striking efficacy in SLE (*15, 16*). CAR19 T cells traffic into lymphoid tissue leading to robust B cell depletion, correlating with a rapid decline in autoantibody titers and sustained clinical benefit (*17*). Notably, CAR T cell numbers decline in most subjects, leading to reconstitution of transitional and naïve B cells within a few months post-therapy (*16*). Surprisingly, despite B cell reconstitution, most patients in Phase I/II clinical trials have exhibited sustained reductions in autoantibodies and disease activity and sustained clinical benefit, referred to as immune reset (*18, 19*). The most likely mechanism for CAR T benefit, is rapid elimination of activated EF and GC cells, memory B cells and/or CD19^+^ PB and short-lived PCs (*20*). However, the precise nature and duration of tolerance remain unclear.

CAR19 T cell therapy is not predicted to impact disease-relevant genetic alterations in B cell function, such as dysregulated B cell-intrinsic signals that promote generation of a self-reactive B cell compartment and/or signals that promote expansion of autoreactive clones during EF and GC responses (*21–23*). Similarly, CAR19 T cell therapy has not been demonstrated to directly modulate Tfh or regulatory T cell (Treg) numbers or activity. Importantly, multi-parameter analyses of LN fine needle aspirates from SLE patients demonstrate both progressive loss of B cell tolerance (across GC, ABC and PCs) and Tfh/Tph expansion (*24*); implying that, over time, newly generated self-reactive B cells may undergo Tfh or Tph-driven activation driving disease relapse. Additionally, CAR19 T cell therapy (*25, 26*) is associated with potential safety concerns. Among patients reported across early clinical studies, a significant proportion exhibited SAEs including: Cytokine Release Syndrome (CRS, 55.9%); Immune Effector Cell-Associated Neurotoxicity Syndrome (2.8%), grade 3 or 4 hematological toxicity (40%); hypogammaglobulinemia (25.5%); and Immune Effector Cell-Associated Hemophagocytic Lymphohistiocytosis-like Syndrome or Macrophage Activating Syndrome (1.4%) (*19*). Collectively, these issues highlight the importance of identifying alternative approaches capable of safely mediating sustained immune tolerance in SLE.

Tregs are essential for the maintaining tolerance and preventing autoimmunity (*27*). In multiple autoimmune diseases, including SLE, Treg number and function are diminished, contributing to immune dysregulation (*28*). Restoring or enhancing Treg activity has emerged as a promising strategy for re-establishing tolerance and preventing disease progression (*29–31*). While adoptive transfer of polyclonal Tregs has shown only limited efficacy in early-stage clinical studies, engineering antigen-specific Tregs enables targeted trafficking to disease-relevant sites and potent, localized bystander suppression (*32, 33*). Previously, we described generation of engineered Tregs (EngTreg) from CD4+ T cells using homology-directed repair (HDR)-mediated insertion of a strong promoter element, MND, into the endogenous *FOXP3* locus (*34*). This platform was additionally enhanced via delivery of TCR or CAR constructs to confer antigen specificity (*35–37*) and introduction of a chemically inducible IL-2 receptor signaling complex (CISC) (*38, 39*). Paring the CISC to FOXP3 expression allows for selective IL-2 signaling to HDR edited cells, supporting in vitro manufacturing and in vivo engraftment, persistence, and therapeutic function.

To exploit the possible therapeutic benefit of EngTreg in SLE, here we developed a CD19-directed CAR EngTreg (CAR19 EngTreg) platform to directly modulate both the pathogenic B cell compartment and co-localized Teff and APC populations in SLE. Unlike conventional CAR19 Teff, CAR19 EngTreg are predicted to concurrently suppress both autoreactive B cell and Tfh (or Tph) activation in lymphoid and tissue-resident environments. By coupling B cell targeting with the suppressive program of FOXP3+ Tregs, this approach aims to directly target pathogenic B–T cell crosstalk to establish sustained tolerance. We describe the generation of human and mouse surrogate CAR19 EngTreg, define their phenotypic and functional properties, and demonstrate their capacity to durably modulate B and T cell responses in vitro and in vivo. Our findings establish CAR19 EngTreg as a mechanistically distinct immunotherapy for SLE that contrasts with the dominant cytotoxic paradigm of CAR19 Teff cells.

## RESULTS

### Generation of human CAR19 EngTreg

To initially generate a CAR19 EngTreg cell product, we utilized a previously validated, lentiviral (LV) plus *FOXP3* HDR editing strategy (*37*) (fig. S1A). Primary CD4+ T cells were stimulated with CD3/CD28 beads and transduced with a LV vector utilizing the MND (myeloproliferative sarcoma 103 virus enhancer, negative control region deleted, dl587rev primer-binding site substituted) promoter to drive expression of an anti-human CD19 CAR (fig. S1B). Two days after LV transduction, cells were gene edited using a *FOXP3* targeted RNP and AAV6 vector containing an HDR template designed to introduce the MND promoter and synthetic rapamycin-regulated IL-2 receptor (IL-2 CISC) in frame with endogenous FOXP3 (fig. S1C). Three days post-editing, CAR surface expression was >85% (fig. S1B, lower panel) and FOXP3 HDR rates reached ∼20-35% across donors (fig. S1C, lower panel). Control cells were HDR-edited without receiving CAR19 LV (polyclonal EngTreg) or were neither edited nor LV transduced (Mock). Following removal of IL-2 and a 7-day expansion of HDR edited cells in rapamycin media to select for IL-2 CISC expression, products were re-stimulated with CD3/CD28 beads and cultured until CISC^+^ cells reached >80%. CAR19 and polyclonal EngTreg exhibited equivalent enrichment, expansion and surface Treg immunophenotype (Fig. 1A-D). CAR19 EngTreg exhibited CAR surface expression (Fig. 1B) with the majority of staining dual-positive for P2A (tracking CISC) and CAR19; however, a minority population of CAR19^+^, P2A^-^ cells were present indicating a low level of CAR19 Teff contamination.

**Fig. 1.**
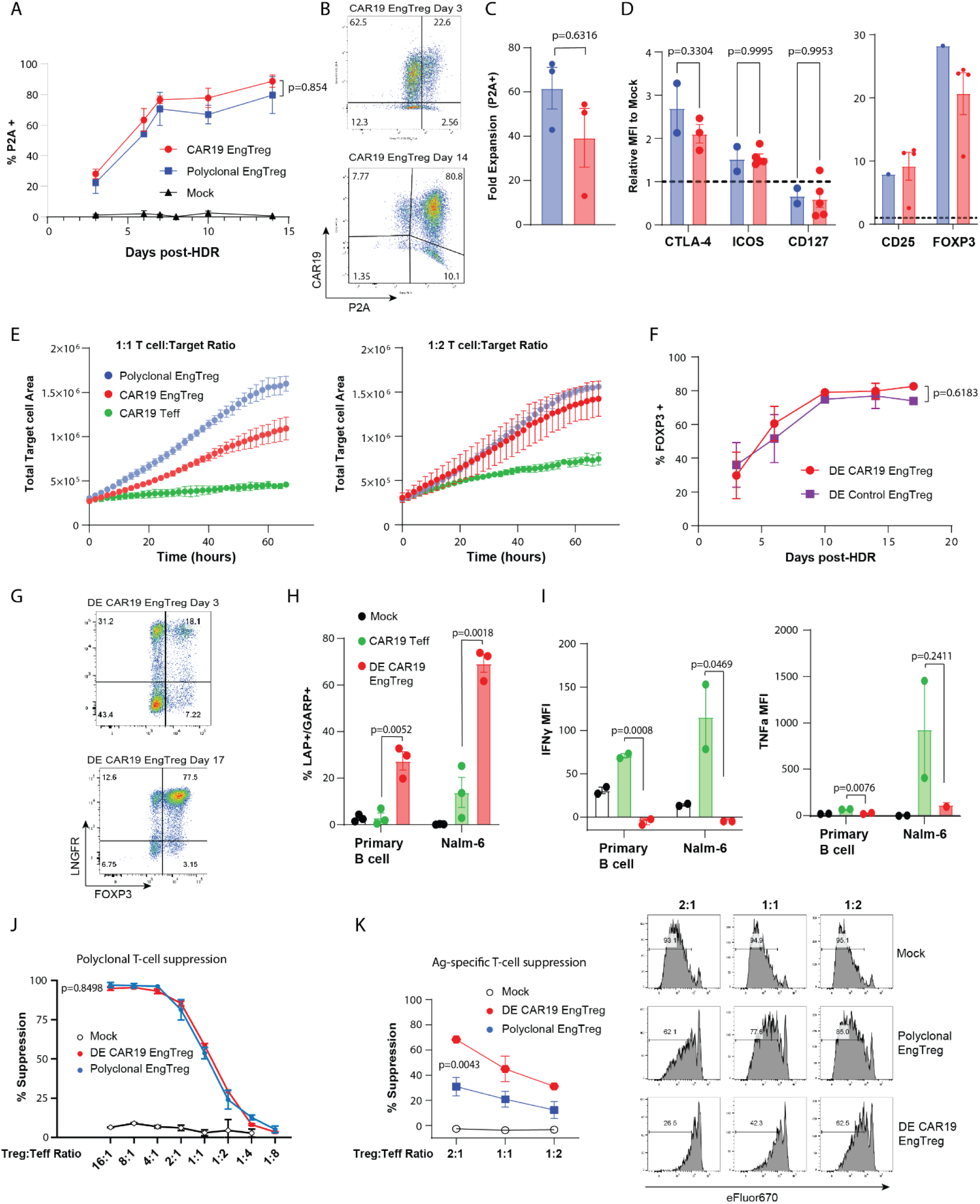
Production of CAR19 EngTreg using alternative strategies. (**A**) Enrichment of LV transduced, CISC-expressing CAR19 EngTreg during rapamycin selection. (**B**) Representative flow for LV CAR19 EngTreg at Day 3 post-editing (upper panel) and post-enrichment (lower panel). (**C**) Fold expansion (Day 3 to endpoint) of P2A+ polyclonal and LV CAR19 EngTreg. (**D**) Immunophenotyping of polyclonal (n=2 donors) vs LV CAR19 EngTreg (n=3 donors). Graphed data shows relative fold change in MFI of indicated Treg markers in EngTreg vs. mock edited cells (dotted line). (**E**) Imaging-based cytotoxicity analysis performed by culturing indicated T cell products with CD19^+^-K562-GFP target cells at 1:1 (left) and 1:2 (right) ratios. Total GFP+ cell area was quantitated continuously for 3 days. (**F**) Enrichment of FOXP3+ expression in DE CAR19, or DE control EngTreg. (**G**) Representative flow for DE CAR19 EngTreg at Day 3 post-editing (upper panel) and post-enrichment (lower panel). (**H**) TGF-b production (%LAP+/GARP+ cells) in DE CAR19 EngTreg, CAR19 Teff, or control, mock-edited T cells co-cultured with CD19^+^ primary B cells. (**I**) Cytokine production in DE CAR19 EngTreg, CAR19 Teff, or mock edited T cells in response to primary B cells or Nalm-6 B cells; graphs show MFI normalized to unstimulated cells. (**J**) Polyclonal suppression assay. Graph shows the % suppression of target Teff proliferation after 4 days in culture with CD3/CD28 activation beads and the indicated ratio of polyclonal or DE CAR19 EngTreg, or mock-edited control suppressor cells. (**K**) CD19 antigen-specific T cell suppression assay. (Left panel) Quantitation of % suppression across all Suppressor:Target ratios. (Right panel) Representative flow images assessing proliferation based upon eFlour670 dye dilution in Teff at D4 following co-culture with the indicated T cell products. All data is representative of n=3 experiments from 3 independent donors with the exceptions of E (single donor), I (two donors), and J (single donor). Graphed data represents mean +/- SEM. P values calculated by unpaired T test.

To assess CAR19 antigen-specific signaling, polyclonal EngTreg, CAR19 EngTreg, or CAR19 Teff cell products (generated by CAR19 LV transduction alone) were cultured with K562 target cells engineered either with or without CD19 surface expression. T cell activation markers were upregulated on both CAR19 EngTreg and Teff products (relative to polyclonal EngTreg) in the presence of CD19+ K562 targets (fig. S1D). Pro-inflammatory cytokine expression (IFN-gamma, IL-6, and TNF-alpha) was lower in CAR19 EngTreg relative to a CAR19 Teff (fig. S1E). To assess target cell cytotoxicity, we performed imaging-based Incucyte assays (Fig. 1E) and observed an intermediate level of cell killing by CAR19 EngTreg relative to CAR19 Teff.

To eliminate the potential confounding impact of contaminating Teff cells present in the LV+HDR CAR19 EngTreg products, we next developed a dual-HDR based engineering strategy. Importantly, this approach is analogous to the strategy employed to generate islet-antigen specific EngTreg in an ongoing T1D clinical trial (NCT06919354, (*36*)) and anticipated to be used for CAR19 EngTreg production in clinical applications. HDR editing at both *TRAC* and *FOXP3* (rather than *FOXP3* alone) was utilized to generate dual-edited CAR19 EngTreg (DE CAR19 EngTreg). We utilized HDR donor constructs including: a) a template targeting *FOXP3* encoding the MND promoter driving cis-linked expression of the split CISC component, IL-2R-g, (*34, 39*) and CAR19 in frame with endogenous FOXP3, thereby linking CAR19 and FOXP3 expression; and b) a template targeting *TRAC* encoding the MND promoter driving cis-linked expression the split CISC component, IL-2Rb, and an LNGFR tag designed to preserve endogenous TRAC expression (fig. S2A). DE CAR19 EngTreg expressed FOXP3 and CAR19, retained endogenous TCR expression (allowing for expansion with anti-CD3/CD28 stimulation) and could be enriched in rapamycin via the CISC signaling complex. Similar to control DE EngTreg expressing a defined TCR (DE Control EngTreg), average initial DE efficiency was 21.15% and reached 79.8% purity by endpoint (Fig. 1F-G, fig. S2C). No differences in Treg immunophenotype were observed between DE CAR19 EngTreg and polyclonal EngTreg control (fig. S2D).

DE CAR19 EngTreg activity in response to antigen engagement was assessed using primary B cells (CD19^mid^) and Nalm-6 (CD19^high^) B cells. In comparison to CAR19 Teff, DE CAR19 EngTreg expressed higher levels of TGF-b (Fig. 1H) and lower levels of pro-inflammatory cytokines (Fig. 1I, fig. S2E) following coculture with target cells. For all cytokines measured, higher responses were observed with Nalm6 target cells, consistent with higher antigen density. As with LV+HDR CAR19 EngTreg, DE CAR19 EngTreg displayed an intermediate level of cytotoxicity toward CD19+ target cells (fig. S2F). To assess DE CAR19 EngTreg suppressive activity, we tested suppression of Teff targets in response to both bead-based (Fig. 1J) and antigen-specific EngTreg stimulation (Fig. 1K, fig. S2G). In the bead-based assay, CAR19 EngTreg exhibited suppressive activity equivalent to polyclonal EngTreg, while CAR-stimulation using primary B cells resulted in significantly stronger suppression by DE CAR19 EngTreg. Taken together, these alternative strategies demonstrate efficient generation of CAR19 EngTreg exhibiting antigen-specific activation and Treg phenotype with a reduced pro-inflammatory and cytotoxic profile relative to CAR19 Teff.

### CAR19 EngTreg directly suppress human PC differentiation and antibody secretion

We next asked whether CAR19 EngTreg could directly suppress primary human B cell activation and/or differentiation in vitro. We examined the impact of CAR19 EngTreg vs polyclonal EngTreg on autologous B cells using an established model of human B cell differentiation capable of generating long-lived, IgG secreting PCs (*40*). In this culture model (Fig. 2A), primary B cells are isolated and sequentially cultured in alternative cytokine conditions to promote activation and expansion (Phase 1), Plasmablast differentiation (Phase 2), and PC generation (Phase 3). Progression through these phases leads to decreasing levels of CD19 expression, increased expression of PC markers including CD38 and CD138, and associated IgG class switch and secretion. This model permitted an analysis of EngTreg activity across stages of B cell differentiation using alternative times of intervention including when CD19 expression remains high, and later, as CD19 is downregulated. For these experiments, CAR19 or polyclonal EngTreg were pre-activated with αCD3/CD28 beads and co-cultured with activated CD19^hi^ B cells at Day 2 (‘Early Intervention’) or differentiating CD19^mid/lo^ B cells at Day 7 (‘Late Intervention’). EngTreg were added at T:B cell ratios of 1:1 or 1:5 (Fig. 2A).

**Fig. 2.**
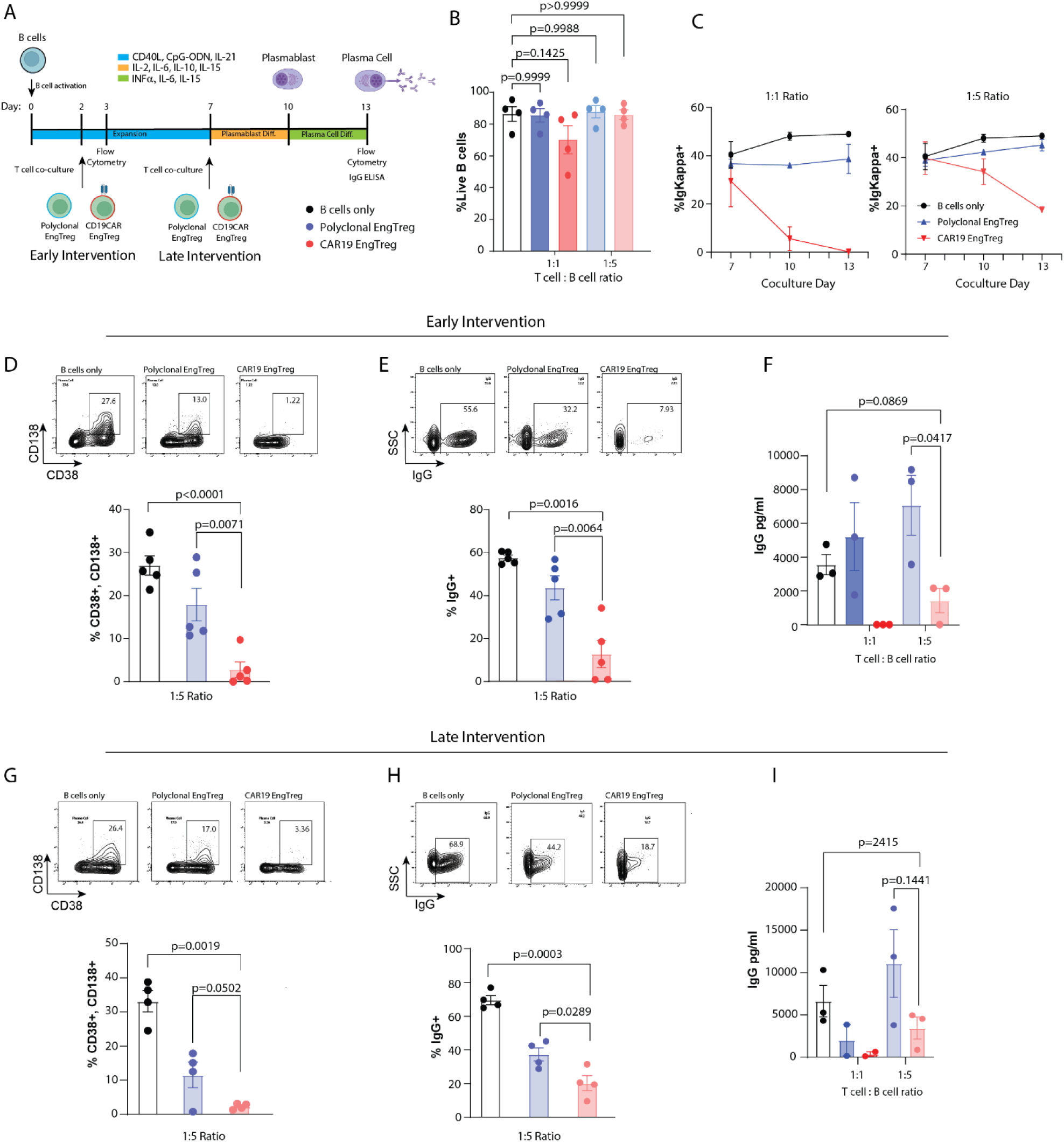
*In vitro* assessment of CAR19 EngTreg impact on primary human PC generation. (**A**) Schematic of in vitro PC culture system and experimental design. Primary B cells were cultured in indicated, sequential cytokine conditions to promote activation and PC differentiation. Activated T cell products were added at indicated Early and Late Intervention timepoints. (**B**) B cell viability at D3, 24h following Early Intervention with indicated T cell products. (**C**) Intracellular Ig-Kappa light chain flow cytometry from indicated culture conditions at days 7, 10, and 13. (**D**) (*Upper panel*) Representative flow cytometry showing PC generation based on CD38^+^CD138^+^ expression at D13 following Early Intervention. (*Lower panel*) Combined data for 1:5 ratio co-culture groups. (**E**) (*Upper panel*) Representative flow cytometry showing intracellular IgG expression on D13. (*Lower panel*) Combined data from 1:5 ratio co-culture groups. (**F**) Total human IgG in media supernatant on D13 for Early Intervention based on ELISA assay. (**G**) (*Upper panel*) PC generation on D13 for Late Intervention studies. (*Lower panel*) Combined from 1:5 ratio co-culture groups. (**H**) (*Upper panel*) Intracellular IgG expression on D13 following Late Intervention. (*Lower panel*) Combined from 1:5 ratio co-culture groups. (**I**) Total human IgG concentration on D13 in Late Intervention studies. Graphed data represents mean +/- SEM. P values calculated by unpaired T test. N>4 independent experiments using 4 independent PBMC donors.

In the Early Intervention timeline, CAR19 EngTreg, exhibited increased CD69 expression within 24h of co-culture with activated B cells; findings consistent with response to CD19 antigen engagement (fig. S3A-B). B cells co-cultured with CAR19 EngTreg exhibited a striking loss of surface CD19 expression despite retention of surface IgM (fig. S3C). Transfer of CD19 from B cells to CAR19 EngTreg was consistent with previously reported trogocytosis of target antigen by CAR Treg (fig. S3D) (*41*). On day 3, B cells co-cultured with CAR19 EngTreg at 1:1 ratio showed a trend toward reduced viability not seen in polyclonal or 1:5 ratio co-cultures (Fig. 2B, fig. S4A). B cell viability declined at later timepoints across all culture conditions, as previously reported (*40, 42*). As a surrogate for B cell viability over time, we monitored intracellular Ig-kappa light chain expression, anticipated to be maintained irrespective of differentiation status (Fig. 2C). While the proportion of Ig-kappa+ cells was stable through D13 in B cell only or polyclonal EngTreg co-cultures, Ig-kappa+ cells decreased in co-cultures with CAR19 EngTreg at 1:1 ratio; consistent with cytotoxic activity of CAR19. In contrast, at the lower 1:5 T:B cell ratio, loss of Ig-kappa+ cells was attenuated. Strikingly, co-culture with CAR19 EngTreg led to a marked reduction in both the proportion of CD38^+^CD138^+^ PCs and in intracellular IgG expression; findings distinct from co-cultures with polyclonal EngTreg or B cell only controls (Fig. 2D-E, fig. S4B). IgG secretion was also decreased in CAR19 EngTreg co-cultures (Fig. 2F). Late Intervention studies demonstrated similar impacts of CAR19 EngTreg on PC proportion and IgG expression. However, co-culture with polyclonal EngTreg also mediated partial inhibition of differentiation in this setting (Fig. 2G-I).

Collectively, these data imply that CAR19 EngTreg can directly suppress B-cell differentiation in-vitro, although this effect may be linked to cytotoxicity, especially at higher T cell:B cell ratios. To investigate suppression independent of B cell killing, we generated Perforin (PRF-1) knockout CAR19 EngTreg by modifying our LV+HDR production strategy, delivering a mixed RNP containing FOXP3 and PRF-1 sgRNAs (fig. S5A). The resulting cells showed high levels of FOXP3 and CAR19 expression (fig. S5B) and efficient PRF-1 gene knockout (KO) based on genomic DNA indel analysis (fig. S5C). Perforin-KO CAR19 EngTreg exhibited a Treg immunophenotype, activation-dependent TGF-b production, and antigen-specific T cell suppression (fig. S5D-F). As anticipated, Perforin-KO CAR19 EngTreg exhibited reduced cytotoxicity toward CD19+ target cells in-vitro (fig. S5G). Consistent with these observations, in primary B cell co-culture assays, perforin-KO CAR19 EngTreg reversed loss of Ig-kappa+ cells at later timepoints (fig. S6A). Importantly, perforin-KO CAR19 EngTreg retained the capacity to limit CD38^+^/CD138^+^ PC differentiation in the context of Early or Late Intervention, supporting the conclusion that CAR19 EngTreg can directly modulate B cell differentiation via mechanism(s) independent of cytotoxicity (fig. S6B-C).

### CAR19 EngTreg target B cells but exhibit reduced inflammatory cytokine production in vivo

To explore B cell targeting by CAR19 EngTreg in vivo, we utilized a humanized NBSGW mouse model (Fig. 3A) that exhibits high levels of human hematopoietic cell engraftment including human myeloid and B cells in bone marrow (BM) and spleen (*43*) following adoptive transfer of CD34+ hematopoietic stem and progenitor cells (HSPC). To establish humanized mouse cohorts, NBSGW mice were transplanted with 2 million HSPCs after low-dose busulfan conditioning (Fig. 3A). At week 14 post-transplant, peripheral blood (PB) displayed an average of 21.5% human cell chimerism including a predominant population of CD19+ B cells (fig. S7A-C). To assess in vivo activity of autologous CAR19 products, DE CAR19 EngTreg or CAR19 Teff were adoptively transferred into humanized recipients and cohorts were sacrificed at D7 post transfer to assess T cell engraftment, B cell depletion and cytokine production. CAR19 Teff recipients had significantly greater weight loss in comparison to EngTreg recipients (Fig. 3B). In both T cell transfer groups, endpoint flow analysis revealed expansion of CD3+ T cells (Fig. 3C, top row), dominated by proliferating Ki67+ CD4 T cells (Fig. 3D-E) and a reduction in CD19+ B cells (Fig. 3C, bottom row). Serum collected at D4 post-T cell transfer showed similar levels of perforin, granulysin and sFas, while Granzyme A and sFasL level were higher in CAR19 Teff vs EngTreg recipients (fig. S7D) indicating that B cell depletion in DE CAR19 EngTreg recipients was, in part, due to direct cytotoxic activity. Strikingly, endpoint serum analysis revealed significantly higher levels of IFN-gamma, CXCL10, and IL-2 in mice receiving CAR19 Teff (Fig. 3F). Thus, consistent with our in vitro findings, CAR19 EngTreg cells can expand and target human CD19+ B cells in vivo while exhibiting a reduced inflammatory cytokine profile relative to CAR19 Teff.

**Fig. 3.**
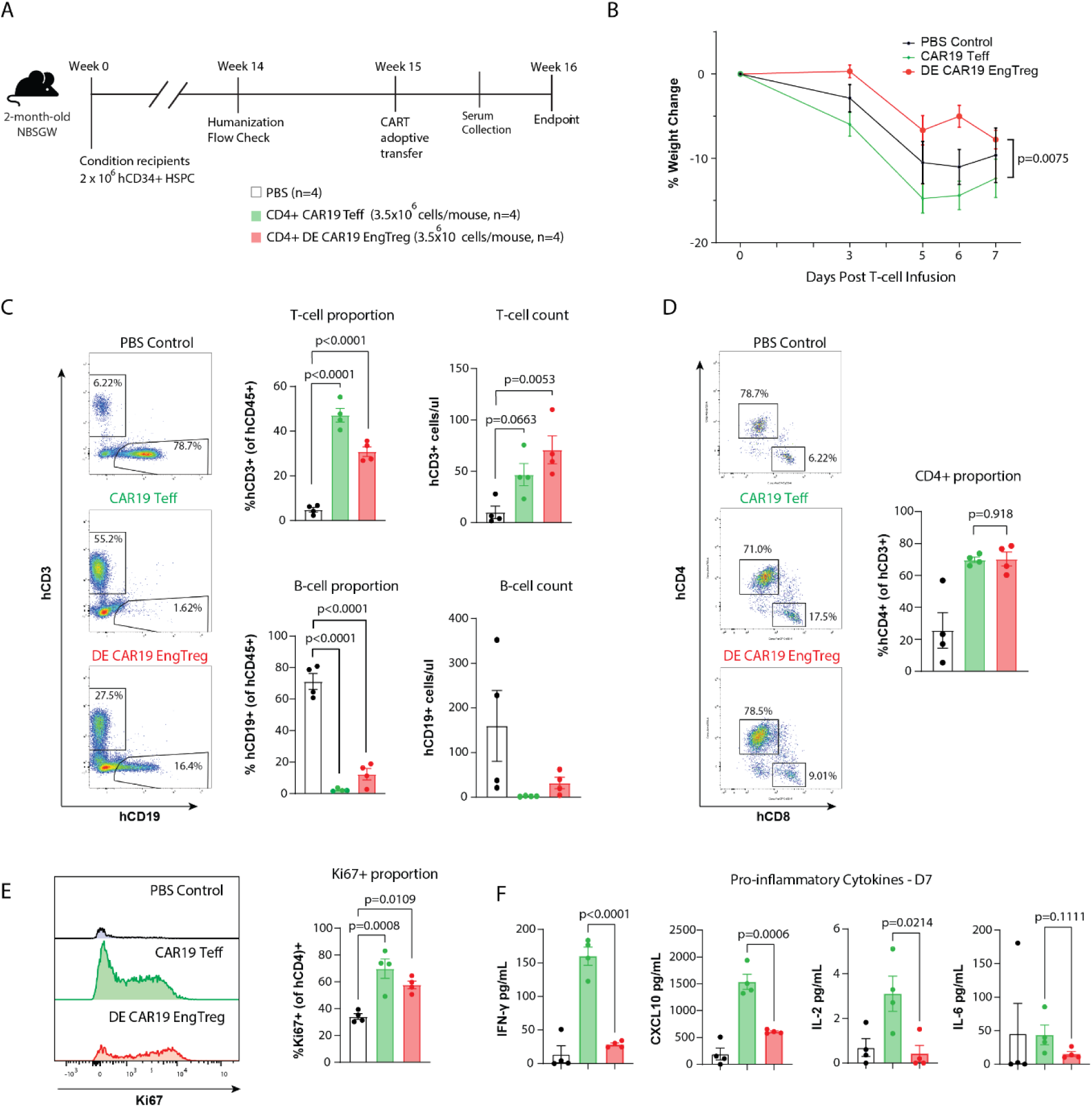
Adoptive transfer of CAR19 EngTreg mediates B cell depletion with reduced inflammatory cytokine production in a humanized mouse model. (A) Timeline and schematic of autologous T cell adoptive study in humanized NBSGW mouse. (B) % weight change from day of T cell transfer for indicated groups. Graphed data is mean +/- SEM. P-value calculated by 2-way ANOVA. (C) (*Left panels*) Representative flow cytometry showing B and T cells in spleen at endpoint gated on human CD45^+^ cells and indicated markers. (*Right panels*) Proportion and cell counts for CD3^+^ T (*top*) and CD19^+^ B cells (*bottom*). (D) (*Left panels*) Representative flow cytometry showing proportion of splenic CD4^+^ and CD8^+^ T cells (within CD3^+^ gate). (*Right panel*) Combined data showing proportion of CD4^+^ T cells. (**E**) (*Left panel*) Representative histograms and (*right panel*) combined data showing proportion of Ki67^+^CD4^+^ T cells. (**F**) cytometric Bead Array (CBA) assessment of serum showing inflammatory cytokines at endpoint. Data shown from single experiment using autologous HSPCs and T cell products. For C-F, data shown as mean +/- SEM, p-values calculated by unpaired T-test.

### Human CAR19 EngTreg reduce inflammatory lesions in a pristane mouse model

To explore the function of CAR19 EngTreg in the context of systemic inflammation, we generated humanized NSG mice using human cord-blood derived HSPCs and treated cohorts with pristane prior to adoptive allogeneic T cell transfer (*44, 45*). We utilized 8-week-old adult NSG mice transplanted with 80,000 CD34+ human cord-blood derived HSPCs after conditioning with total body irradiation (Fig. 4A). At time of pristane dosing (week 12 post HSPC transfer), all mice exhibited human immune system reconstitution including circulating hCD45+ cells, predominantly composed of B cells (data not shown). Three weeks after pristane administration, animals were divided into treatment groups and dosed with 3.5 million allogeneic CAR19 EngTreg or CAR19 Teff, or 10mg/kg of inebilizumab, an anti-CD19 monoclonal antibody (mAb) (*46*). CAR19 EngTreg products were generated by LV transduction + HDR editing. Control groups received pristane and PBS (disease control), or no pristane (humanization-only control). In addition to weekly PB monitoring, cohorts were sacrificed at weeks 4 or 8 post-treatment to assess organ inflammation. As expected, pristane dosing led to reduced levels of hCD45+ cells in PB relative to non-pristane controls (Fig. 4B). Circulating CD19+ B cells were reduced in mice receiving either T cell product or inebilizumab, with greatest impact observed in response to mAb treatment (Fig. 4C). Human cytokine levels were assessed on D0 and D7 post T cell transfer. CAR19 Teff treated mice exhibited the highest increase in proinflammatory cytokines (including IFN-gamma and TNF-alpha) and cytotoxic granules (including perforin, granulysin, Granzyme A, and sFasL) relative to recipients of CAR19 EngTreg, mAb, or pristane alone (Fig. 4D).

**Fig. 4.**
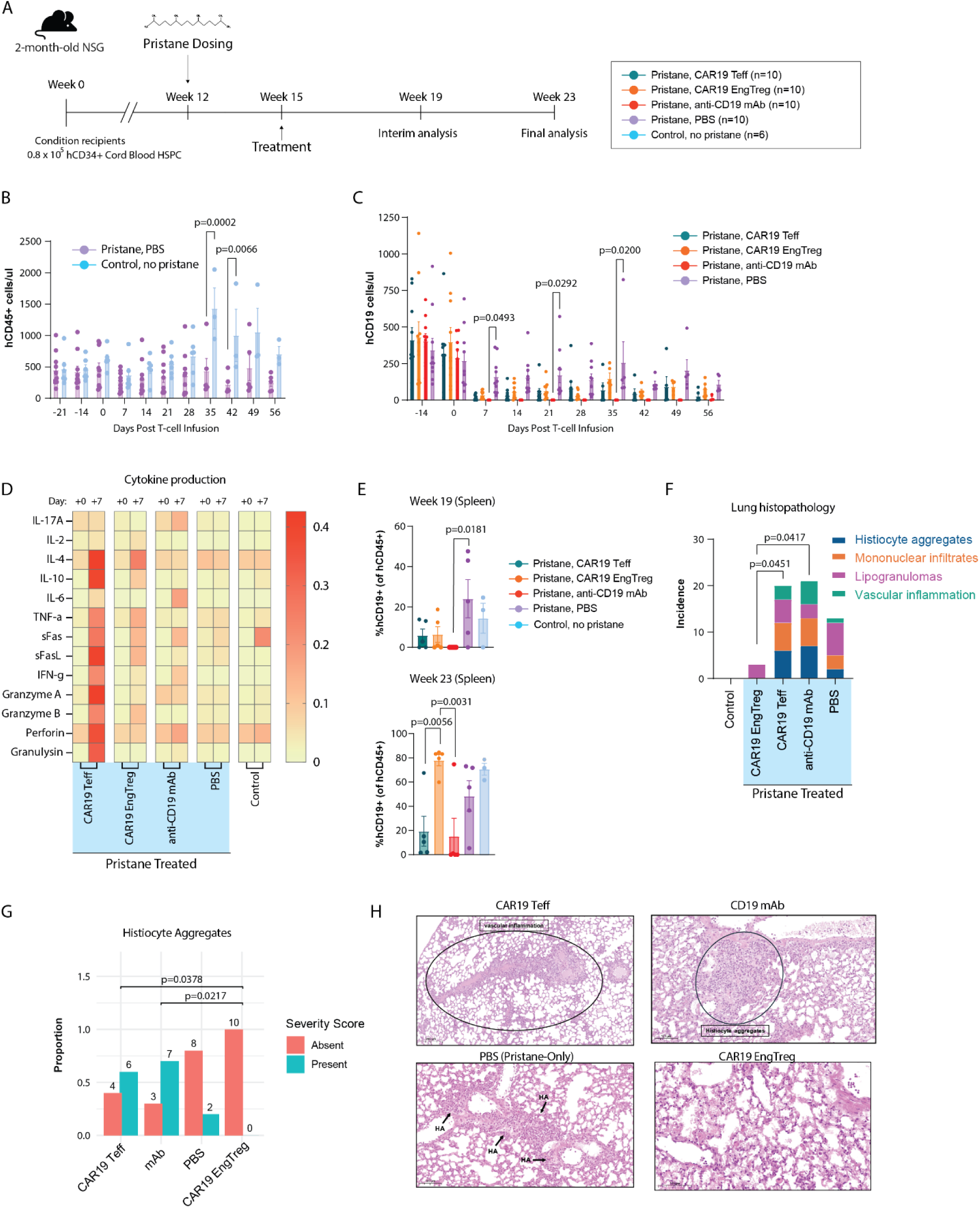
CAR19 EngTreg protect from inflammatory lesions in a pristane-induced inflammatory model in humanized mice. (**A**) Timeline and schematic of humanized mouse pristane treatment study. (**B-D**) Absolute counts of: (**B)** human CD45+ cells, (**C**) CD19^+^ B cells, and in the PB as assessed by flow cytometry for indicated groups. Results are expressed as mean +/- SD. Two-way ANOVA with Tukey correction for multiple comparisons. (**D**) Relative levels of human inflammatory cytokines in the PB prior to and at D7 after treatment assessed by CBA assay. (**E**) Flow based CD19+ B cell proportion in spleen 4-weeks (upper) and 8 weeks (lower) post-T cell transfer (humanization weeks 19 and 23, respectively). (**F**) Incidence of mice bearing indicated inflammatory lesions within each group based upon histopathological evaluation in lungs. (**G**) Proportion of mice in each treatment group with or without histiocyte aggregates. (**H**) Representative lung H&E-stained histological images from indicated treatment groups.

Mice from each group were sacrificed at weeks 19 and 23, with sustained loss of splenic CD19+ B cells observed in mAb treated mice (Fig. 4E). Splenic B cells had recovered by week 23 in CAR19 EngTreg treated mice, suggesting a more transient depletion in comparison with CAR19 Teff or mAb. We assessed lung and liver histology at both endpoints for evidence of inflammatory tissue damage. Histiocyte aggregates, mononuclear infiltrates, lipogranulomas, and vascular inflammation were increased in lung tissue of pristane treated mice, and only CAR19 EngTreg treated animals showed a significant reduction in these features (Fig. 4F-H). In liver, histocyte aggregates and lipogranulomas were observed only in pristane treated animals (fig. S8A). While histocyte aggregates occurred at similar or greater frequency in CAR19 Teff and mAb treated mice relative to PBS controls, these and all other scored markers of inflammation were absent in CAR19 EngTreg recipients (fig. S8B-C). Overall, adoptive transfer of CAR19 EngTreg in this humanized pristane model led to both reduced levels of inflammatory mediators in serum and a reduction in inflammation-associated tissue damage relative to CAR19 Teff.

### Murine surrogate CAR19 EngTreg limit TLR7 driven inflammatory responses in vivo

To explore the function of CAR19 EngTreg cells in immune-competent settings, we developed surrogate mouse HDR-based products for CAR19 EngTreg and Teff using a previously validated mouse CD19 CAR construct (*47*) (fig. S9A). To generate CAR19 mouse (m)EngTreg, a LNGFR.P2A.CD19CAR cassette was knocked into the first coding exon of *Foxp3* to generate cells co-expressing CD19CAR and FOXP3. To generate CAR19 mTeff, an LNGFR.P2A.CAR19 cassette was knocked into the Rosa26 safe harbor locus. Two days after editing, enrichment for tLNGFR surface expression yielded cell products with >90% CAR expression (fig. S9B-C).

Gain-of-function mutations in Toll-like receptor 7 (TLR7) are implicated in variants of human lupus and are sufficient to mediate lupus-like disease in mice (*48, 49*). Epicutaneous application of the TLR7 agonist, imiquimod (IMQ), mimics aspects of cutaneous lupus erythematosus (CLE) pathology, including increased cytokines and autoantibody production (*50, 51*). A short-term IMQ model study was selected to compare the early safety and therapeutic impact of CAR19 mEngTreg vs. mTeff. As a means of improving engraftment in immune-competent hosts, recipient mice were preconditioned with cyclophosphamide (Cy), a strategy to mimic clinical CAR-T conditioning and previously employed in murine CD19 CAR studies (*47*). To further promote mEngTreg engraftment and partially mimic IL-2 CISC activity in the human DE EngTreg product, a cohort of mEngTreg-treated mice and disease only mice (as cytokine-support controls) received low-dose (LD) murine IL-2 (fig. S10A). Both CAR19 mEngTreg and CAR19 mTeff were detectable in blood at D5 post transfer (fig. S10B). CAR19 mEngTreg engraftment remained stable at 3-5% of the CD4 compartment from D5 in PB to D12 in spleen (fig. S10B-C) and was not impacted by IL-2 dosing. Notably, a significantly higher proportion of engrafted CAR19 mTeff were detected in spleen at study endpoint, likely due to higher proliferative capacity (fig. S10C). Only CAR19 mTeff recipients exhibited a significant depletion of splenic B cells (fig. S10D). In contrast, only CAR19 mEngTreg treated animals showed improvement in CLE-like skin thickening (fig. S10E), reduced splenomegaly (fig. S10F) and a reduction in inflammatory monocytes (fig. S10G). To further characterize the inflammatory milieu at sacrifice, splenocytes were stimulated with PMA/Ionomycin prior to CBA analysis of media supernatant. CAR19 mTeff recipients, in comparison to CAR19 mEngTreg treated mice, exhibited higher levels of inflammatory cytokines and chemokines including IL-6, IFN-gamma, and CCL5 (fig. S10H). Together, these data indicate that CAR19 mEngTreg, but not CAR19 mTeff, can mitigate CLE-like tissue inflammation without amplifying inflammatory responses that can result in CRS-like cytokine production, monocyte activation and splenomegaly.

### CAR19 mEngTreg persist and limit renal pathology in a murine SLE model

Next, we evaluated CAR19 mEngTreg activity in the B6.NZMSle1/Sle2/Sle3 (Sle123) lupus model (Fig. 5A), which recapitulates key genetic and immunopathologic features of human SLE (*52, 53*). Seven-to eight-month-old Sle123 mice were treated with 3 million B6-derived CAR19 mEngTreg following cyclophosphamide preconditioning. CAR19 mEngTreg-treated mice and a subset of disease control mice also received LD murine IL-2. Age-matched wild type B6 mice were included as a healthy control group. Transferred CAR19 mEngTreg were detectable in blood for at least 2 weeks post-transfer and correlated with a transient reduction in circulating CD19⁺ B cells (Fig. 5B) and reduction of serum autoantibodies at week 4 (Fig. 5C). Recipient animals were sacrificed at 11-12-mo of age, a timepoint where Sle123 mice exhibit glomerulonephritis and multiple autoimmune disease findings (*54*). Strikingly, CAR19 mEngTreg persisted at low levels in the spleen for >4 months as identified by expression of the congenic marker CD45.1 and co-expression of surface LNGFR, or CAR, and intracellular FOXP3 (Fig. 5D). Serum neutrophil gelatinase-associated lipocalin (NGAL), a biomarker of kidney injury, was significantly reduced in CAR19 mEngTreg recipients compared to Sle123 disease controls (Fig. 5E) and correlated with improved renal histopathology, characterized by reduced glomerular diameter and combinatorial histopathology scores (Fig. 5F-G).

**Fig. 5.**
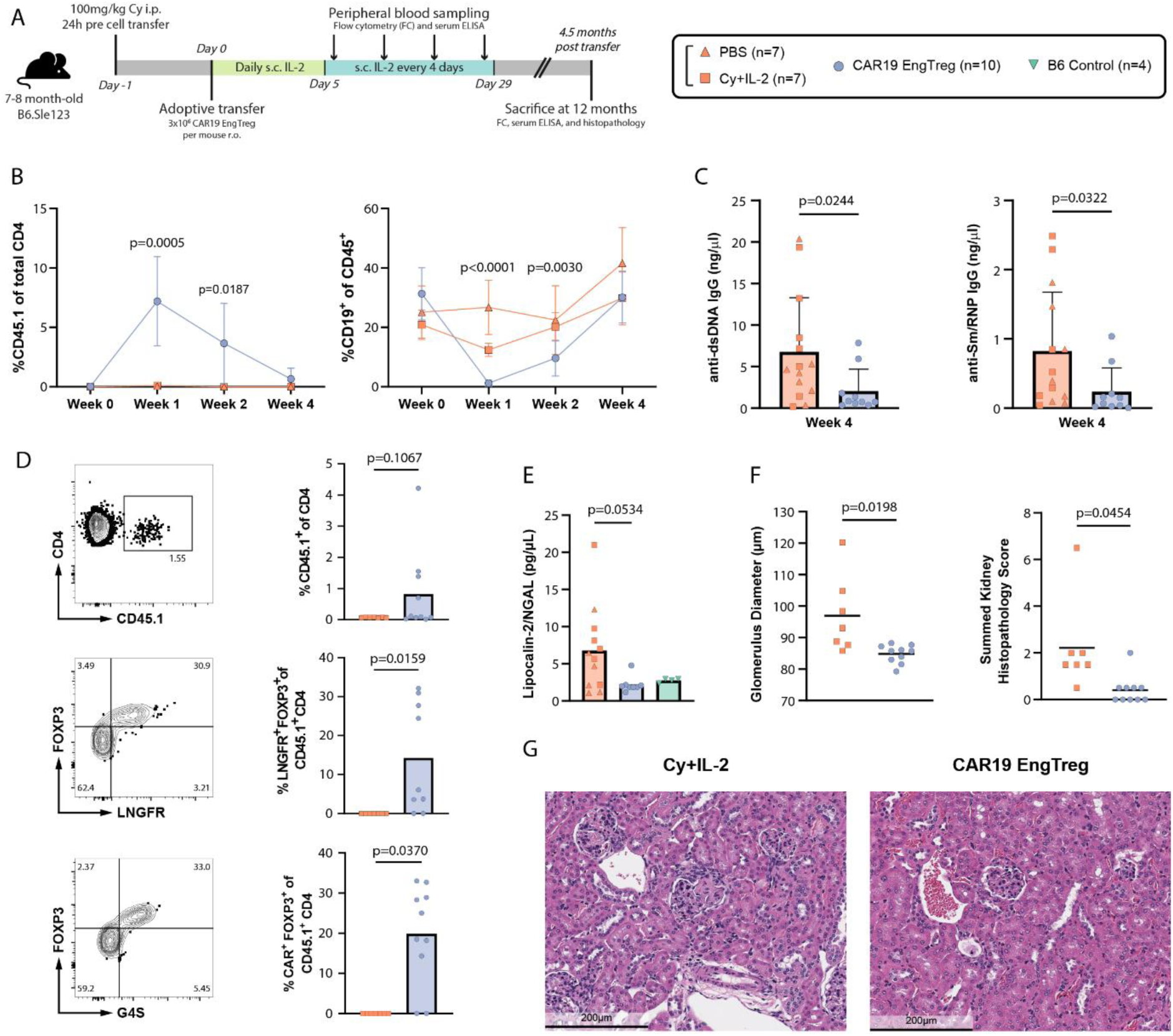
CAR19 mEngTreg engraft long-term and improve renal function in the SLE123 lupus model. (**A**) Experimental schematic and timeline for SLE123 mouse model; i.p., intraperitoneal; r.o., retro-orbital; s.c., subcutaneous delivery. (**B**) (*Left panel*) Frequency of CAR19 mEngTreg (CD45.1⁺ cells within CD4⁺ T cells) in PB over time (0-4 weeks) post-adoptive transfer. (*Right panel*) Frequency of CD19⁺ B cells within CD45⁺ leukocytes over time. Statistical significance, determined by two-way ANOVA, reflects comparisons between the CAR19 EngTreg and Cy+IL-2 groups. (**C**) dsDNA and Sm/RNP autoantibodies in serum of mice at 4 weeks post-adoptive transfer. (**D**) (*Left panel*) Representative flow cytometry and (*right panel*) combined data showing proportion of CAR19 mEngTreg in the spleen at endpoint (4.5 mo. post-adoptive transfer) identified by expression of CD45.1 and co-expression of FOXP3 and LNGFR or CD19 CAR (assessed by G4S linker staining). (**E**) Serum NGAL in mice at endpoint. (F) Glomerular diameter (left) and cumulative renal histopathology (right) in mice at endpoint. (G) Representative images of scored kidney sections (200µm field). Error bars represent mean ± s.d. Statistical significance for panels C-F determined by Welch’s t-test.

### CAR19 mEngTreg modulate both pathogenic B and T cells and reduce autoantibodies

Serum anti-dsDNA and anti-Sm/RNP autoantibodies were reduced in CAR19 mEngTreg recipients at multiple interim bleed timepoints through week 8 post-treatment (Fig. 6A, fig. S11A) and remained reduced at study endpoint (Fig. 6B). Next, endpoint serum samples were analyzed by autoantigen microarray (*55*). Strikingly, CAR19 mEngTreg treatment reduced reactivity across a broad spectrum of self-antigens in comparison to PBS or Cy/IL-2 treated Sle123 mice (Fig. 6C, fig. S11B). Consistent with decreased autoantibodies, high-parameter flow cytometry revealed a reduction in splenic germinal center (GC) B cells (Fig. 6D). Age-associated B cells (ABCs), key extrafollicular drivers of autoantibody production, were also reduced (Fig. 6E). H&E staining of spleen sections from mEngTreg-treated mice showed smaller and fewer white pulp foci compared to disease controls, indicative of reduced splenic lymphoid follicles, consistent with the reduction in GC B cells (Fig. 6F-H).

**Fig. 6.**
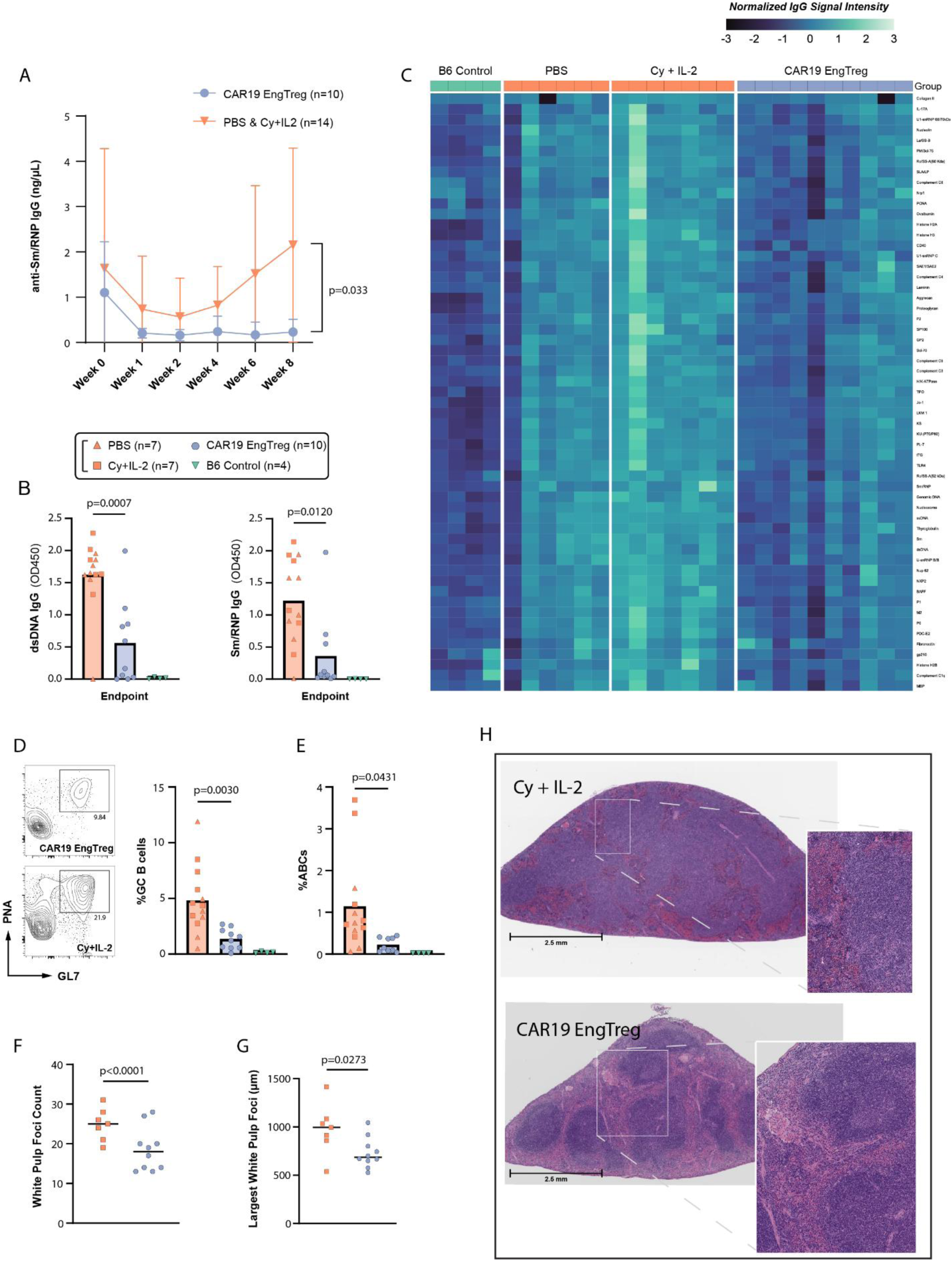
CAR19 mEngTreg reduce pathogenic B cell subsets and broad-spectrum autoantibodies in the SLE123 lupus model. (**A**) Serum anti-Sm/RNP IgG was measured by ELISA at baseline prior to pre-conditioning with cyclophosphamide (week 0) and at the indicated timepoints post adoptive cell transfer. Significance determined by repeated measures two-way ANOVA. (**B**) dsDNA (left) and Sm/RNP (right) autoantibodies in mice at study endpoint determined by ELISA. (**C**) Heatmap showing log2-normalized IgG signal intensities measured by autoantigen microarray using endpoint serum obtained from indicated groups. Each column shows data from an individual animal and color at top indicates treatment groups with target antigens indicated on right margin. (**D**) Representative flow cytometry showing GC B cells for CAR19 EngTreg (top) and Cy+IL-2 (bottom) treatment groups with proportion of splenic GC B cells at study endpoint (left panel). (**E**) Proportion of ABC B cells (defined as CD11b+, CD11c+, CXCR5^lo^, CD21^lo^) in spleen at study endpoint for indicated groups. (**F**) Number of white pulp foci and (**G**) size of largest white pulp foci (µm) in spleens at study endpoint for indicated groups. (**H**) Representative H&E-stained sections of spleen in Cy+IL-2 control (top) and CAR19 EngTreg (bottom) recipient mice. Error bars represent mean ± s.d. Statistical significance for panels A, D, and E determined by one-way ANOVA. Statistical significance for panels F and G determined by Mann-Whitney U test.

In addition to the impact on B cell mediated pathology, CAR19 mEngTreg treatment altered multiple T cell subsets implicated in human and murine lupus pathogenesis. Notably, CAR19 mEngTreg treatment reduced the proportion of memory CD44⁺CD4⁺ T cells and increased the proportion of naive CD62L^+^CD44^-^ CD4^+^ T cells in the spleen, shifting the balance from memory toward naïve T cell subsets (Fig. 7A-B). In the CD8 compartment, CAR19 mEngTreg treated animals exhibited an increased proportion of CD62L^+^CD44^-^ naïve CD8 T cells (Fig. 7C) and a decreased proportion of KLRG1^+^ CD8 T cells (Fig. 7D), a terminally differentiated, short-lived effector CD8 subset, consistent with reduced chronic antigen stimulation. Most notably, consistent with the observed reduction in GC B cells and diminished splenic lymphoid follicle formation, CAR19 mEngTreg treated mice exhibited a decrease in T follicular helper (Tfh) cells, the primary CD4⁺ T cell subset modulating GC responses (Fig. 7E-F). The proportion of T peripheral helper (Tph) cells, extrafollicular CD4⁺ T cells that provide B cell help outside of GCs was similarly decreased (Fig. 7G). Finally, CAR19 mEngTreg recipients exhibited a restoration in the Tfh:Tfr ratio equivalent to healthy B6 controls (Fig. 7H-J). Collectively, these results demonstrate that CAR19 mEngTreg persist long-term in vivo in lupus prone Sle123 recipient mice and reshape both pathogenic B and T cell compartments, providing durable immune tolerance.

**Fig. 7.**
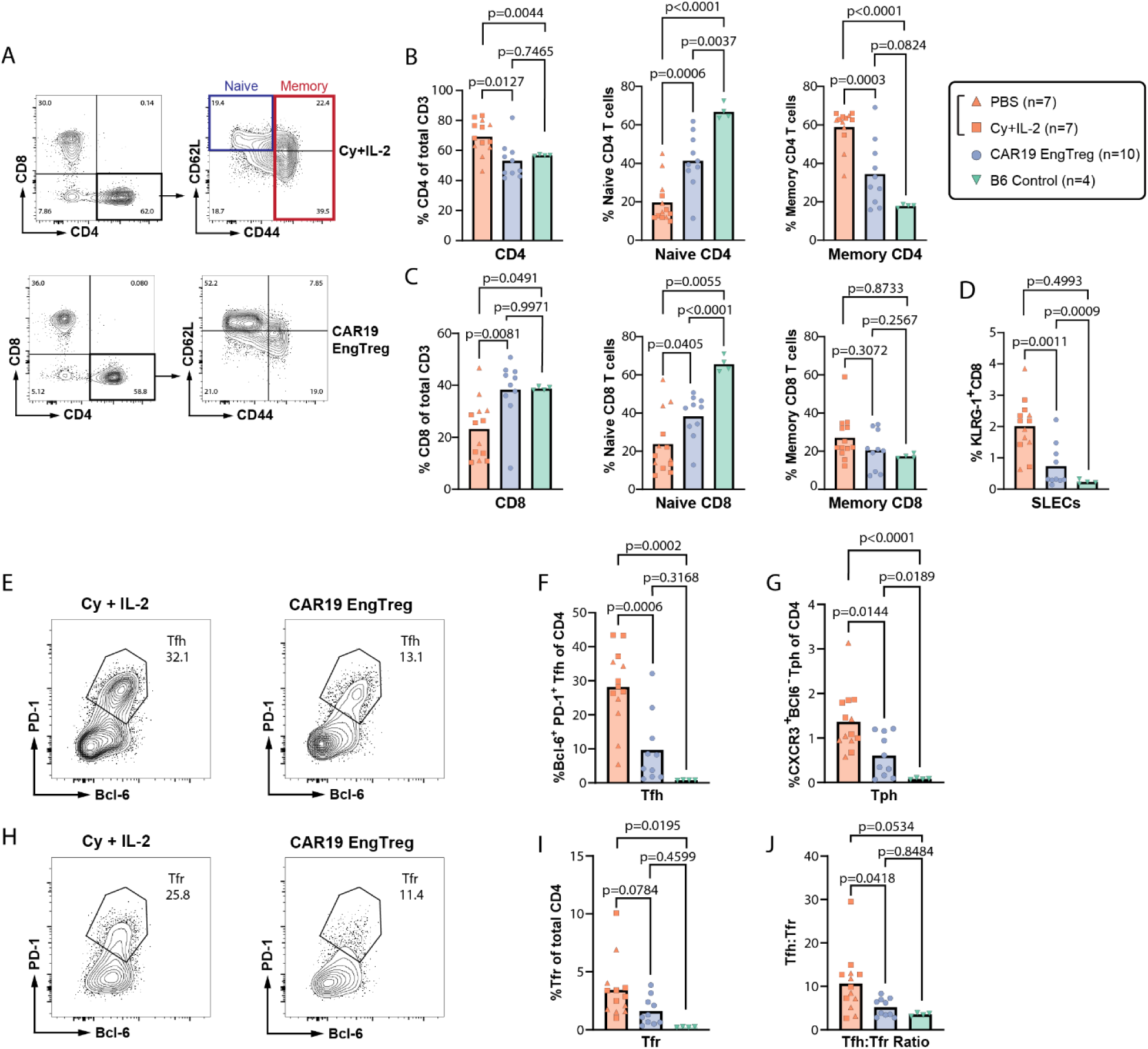
CAR19 mEngTreg reduce pathogenic T cell subsets in the SLE123 lupus model. (**A**) Representative flow cytometry plots showing gating of splenic CD4⁺ T cells (left) and subsequent analysis of CD62L and CD44 expression to define naïve and memory subsets (right) in CAR19 mEngTreg (top) and Cy+IL-2 only (bottom) treated animals. (**B**) Bar graphs show proportion of total (left), naïve (center), and memory (right) CD4⁺ T cells in indicated treatment groups and B6 control mice at endpoint. (**C**) Bar graphs show relative abundance of total (left), naïve (center) and memory CD8^+^ T cells at endpoint. (**D**) Proportion of KLRG-1^+^ short-lived effector (SLEC) CD8+ T cells at endpoint. (**E**) Representative flow cytometry showing T follicular helper (Tfh) cells in Cy+IL-2 only (left) and CAR19 EngTreg (right) treated mice. Bar graphs showing proportion of (**F**) Tfh and (**G**) Tph cells at endpoint. (**H**) Representative flow cytometry showing T follicular regulatory cell (Tfr) cells in Cy+IL-2 (left) and CAR19 mEngTreg (right) treated animals. (**I**) Bar graphs of proportion of Tfr at endpoint. (**J**) Ratio of splenic Tfh:Tfr at endpoint. Error bars represent mean ± s.d. Statistical significance for all comparisons determined by one-way ANOVA.

Next, we examined CAR19 mEngTreg activity in the more severe,Sle1b.yaa, lupus model (*56*) that exhibits increased genetic drive mediated by *Tlr7* gene duplication leading to amplification of TLR7-dependent immune cell activation (fig. S12A). Transferred CAR19 mEngTreg were present in PB to study endpoint where they comprised ∼25% of splenic FOXP3⁺ cells (fig. S12B-C). Circulating CD19⁺ B cells declined during the first 2 weeks in both CAR19 mEngTreg recipients and conditioning-only controls, followed by recovery of B cells by week 4 post-transfer (fig. S12D). Similar to our findings in SLE123 mice, CAR19 mEngTreg recipients exhibited reduced anti-dsDNA and anti-Sm/RNP autoantibodies at Week 4 (fig. S12E). At endpoint, spleen weight and CD19⁺B220⁺ B cells (in spleen and PB) were equivalent in mEngTreg recipients and controls (fig. S12F-G). Notably, B220⁺ B cells from CAR19 mEngTreg-treated mice displayed reduced surface CD19 expression in both PB and spleen, consistent with ongoing CAR19-mediated trogocytosis (fig. S12H). At endpoint, CAR19 mEngTreg-recipients exhibited reduced anti-dsDNA and a trend toward reduced anti-Sm/RNP (fig. S12I). Autoantibody microarray analysis demonstrated reduced reactivity across a broad spectrum of self-antigens (fig. S13). CAR19 mEngTreg treatment was associated with a trend toward reduced GC B cell frequencies (fig. S12K), significantly reduced LN CXCR5⁺PD-1⁺ Tfh, (fig. S12J) a trend for reduced CD11c⁺T-bet⁺ ABCs (a TLR7-associated population characteristic of the Sle1b.yaa model) (fig. S12L) and significantly reduced LN memory B cells (fig. S12M). EngTreg treatment was associated with a nonsignificant trend toward reduced composite renal histopathology activity relative to Cy+IL-2 controls, suggestive of attenuation of active inflammatory kidney injury (fig. S12N). These combined data provide complementary evidence that CAR19 mEngTreg persist long term in vivo in a severe murine lupus model leading to partial remodeling of autoreactive and antigen-experienced B and T cell responses.

## DISCUSSION

In this study, we provide evidence that CAR19 EngTreg represent a promising potential cell therapy for SLE, capable of stably suppressing B, T and other cellular interactions central to SLE pathogenesis. Of note, the functional activities of CAR19 EngTreg are largely distinct from the B cell ablative paradigm of CAR19 Teff, now emerging as the first effective cell therapy for SLE. Building on previous work, we utilized dual-locus HDR at *FOXP3* and *TRAC* to generate human EngTreg expressing CAR19 for B cell engagement and IL-2 CISC for persistence in IL-2-limited environments (*38, 39*). In vitro, CAR19 EngTreg retained Treg identity and produced minimal inflammatory cytokines, exhibiting reduced cytotoxicity relative to CAR19 Teff. In co-cultures with activated primary B cells and plasmablasts, CAR19 EngTreg inhibited the formation of PCs and abrogated antibody secretion. Adoptive transfer of CAR19 EngTreg into humanized mice led to B cell depletion with minimal secretion of inflammatory cytokines. Further, in a humanized pristane-induced inflammatory model, CAR19 EngTreg reduced histopathologic lesions, while CAR19 Teff mediated an increase in inflammatory cytokines and pathology. In parallel with human cell studies, we evaluated mouse surrogate CAR19 EngTreg in immune competent autoimmune models. In an imiquimod (IMQ)-induced inflammatory model, mCAR19 EngTreg restrained TLR7-driven inflammation and cutaneous lupus–like pathology while CAR19 Teff exacerbated cytokine production and monocyte recruitment. Finally, in both the B6.Sle123 and Sle1b.Yaa lupus models, mCAR19 EngTreg induced durable immunologic and clinical benefit. EngTreg persisted long-term in lymphoid tissues both strains and recipient mice exhibited sustained reductions in autoantibodies. In B6.Sle123 recipients, these findings correlated with long-term reduction in lymphoid follicles, GC and ABC B cells and CD4^+^Tfh and Tph, and effector memory, CD4^+^ and CD8^+^, T cells. Finally, mCAR19 EngTreg treatment reduced glomerular injury and improved composite renal pathology scores. Together, these findings support the conclusion that CAR19 EngTreg can enforce long-term immune tolerance by suppressing both autoreactive B and T cell circuits and modulating tissue-level inflammation.

A key innovation enabling this work lies in our optimized Treg engineering strategy. We assessed alternative approaches to generate human CAR19 EngTreg and characterized cell products in vitro. Most notably, by modification of a clinically approved method for dual-locus HDR at *FOXP3* and *TRAC*, we generated CAR19 EngTreg that integrate stable, constitutive FOXP3 expression in conjunction with cis-linked CAR19 and a rapamycin-inducible IL-2 signaling complex (CISC). This dual-HDR editing (DE) strategy, facilitated by integrating distinct components of CISC within each targeted locus, permitted efficient in vitro enrichment of DE EngTreg during manufacturing. Our DE approach contrasts with prior CAR19-Treg studies (*45, 57, 58*) using engineered nTreg products that rely on LV transduction of nTreg, where FOXP3 stability, random viral integration, and heterogeneous CAR expression may limit Treg stability and in vivo function. By directly targeting *FOXP3* in CD4+ T cells, our DE approach overcomes key challenges including reliance on rare Treg starting material, stringent FACS sorting, and variable gene transfer efficiencies that can limit manufacturing consistency.

CAR19 EngTreg share several key functional attributes with CAR19 Teff: both specifically recognize CD19, undergo antigen-triggered activation, and mediate B cell depletion in vitro and in vivo (*31, 59*). However, whereas CAR19 Teff mount robust cytotoxic programs marked by antigen-dependent cytotoxicity and high-level inflammatory cytokine production, CAR19 EngTreg maintain a regulatory phenotype and exhibit minimal production of inflammatory cytokines. In comparison with CAR19 Teff, CAR19 EngTreg exhibit reduced cytotoxic potential at lower, potentially more physiologic, effector to target cell ratios. Our in vitro data support the conclusion that CAR19 EngTreg are predominantly suppressive in nature, exhibiting potent regulatory activity in both polyclonal and antigen-specific suppression assays. Together, these results indicate that combining CD19-targeting with stable FOXP3 expression enables potent antigen-specific suppression with limited cytotoxicity relative to conventional CAR19 Teff.

As part of these studies, we directly assessed the capacity of CAR19 EngTreg to modulate primary B cell responses through non-cytotoxic mechanisms. In co-cultures with activated B cells and differentiating plasmablasts, CAR19 EngTreg exhibited suppressive activity toward both populations, inhibiting PC differentiation and antibody secretion. To assess the contribution of cytotoxicity, we generated perforin-KO CAR19 EngTreg. In both antigen-specific T cell suppression assays and in co-cultures with primary B cells, perforin-deficient CAR19 EngTreg retained suppressive activity, suggesting that direct killing is not the primary mechanism limiting in vitro PC generation and antibody secretion. Interestingly, consistent with direct B cell engagement, CAR19 EngTreg mediated CD19 trogocytosis of primary B cells; observations consistent with previous reports of robust trogocytosis of OX40L by OX40L-CAR nTreg (*41, 60*). CAR19 EngTreg expressed surface bound TGF-b and its expression increased following antigen engagement, implicating this pathway in suppression of B cell differentiation as reported in previous work (*58*). Our in vivo modeling further supports the concept that CAR19 EngTreg activate a suppressive program following CAR engagement that restrains inflammatory responses. In two humanized models including a pristane-induced inflammation model, while both CAR19 EngTreg and CAR19 Teff reduced B cell numbers, CAR19 EngTreg recipients exhibited lower levels of inflammatory cytokines and reduced inflammatory histopathology supporting the conclusion that human CAR19 EngTreg exert their impact predominantly via antigen-directed immunosuppression.

To achieve a more rigorous assessment of therapeutic potential, we generated surrogate mouse B cell–targeted EngTreg and tracked suppressive activity at physiologic effector-to-target ratios in vivo in the presence of endogenous immune cell networks and ongoing inflammation. In an IMQ-induced, TLR7-driven cutaneous inflammation model (*51*), mCAR19 EngTreg attenuated skin inflammation and reduced TLR7-driven cytokines despite only a limited impact on B cell numbers. Most notably, we examined the impact of mCAR19 EngTreg on autoreactive B and T cells in two distinct murine SLE models mirroring distinct mechanisms for loss of tolerance, GC dysregulation, autoantibody production, and renal disease characteristic of human SLE (*4, 53, 54, 56*). Strikingly, in the B6.Sle123 lupus model, CAR19 mEngTreg recipients exhibited durable reductions in both GC and ABC B cells and Tfh and Tph cells, T helper populations that iteratively drive aberrant B-cell responses. CAR19 mEngTreg recipients also exhibited a reduction in memory CD4^+^ and SLEC CD8^+^ T cells. Importantly, these impacts led to improved disease outcomes, including reduced glomerular injury, lower NGAL levels, and a sustained reduction across a broad spectrum of autoantibodies assessed using autoantigen arrays. In the Sle1b.Yaa model characterized by more aggressive, TLR-7 driven SLE-like pathology, CAR19 mEngTreg recipients exhibited a sustained reduction in autoantibodies and significantly reduced LN Tfh and MBCs with a nonsignificant trend for reduced splenic GC, ABC, and MBC populations. Collectively, these findings demonstrate that mCAR19 EngTreg exert coordinated suppression and restructuring of autoreactive immune circuits, thereby supporting immune tolerance.

Collectively, our findings suggest that CAR19 EngTreg provide therapeutic impacts that extend beyond the transient B cell depletion achieved by CD19-directed effector therapies. The sustained reduction in both GC and ABC B cells and Tfh/Tph cells in the Sle123 model suggest that CAR19 EngTreg-driven regulatory mechanisms may help to prevent the re-emergence autoreactive effector populations. By simultaneously suppressing multiple cellular programs that drive B-cell activation and systemic inflammation, CAR19 EngTreg modulate not only B cells and autoantibody production, but also the underlying signals from Tfh and Tph that promote the emergence and reactivation of autoreactive B cell clones. Consistent with this concept, direct targeting of Tfh cells—using agents such as abatacept or PD-L1 CAR-NK cells—has been shown to reduce Tfh activity and, consequently, limit B-cell expansion, plasmablast differentiation, and antibody production (*61, 62*). Similarly, CXCR5-CAR Tregs can disrupt Tfh–B cell interactions and suppress Tfh-induced B cell activation in vitro (*63*). Building on these approaches, CD19-directed EngTreg may partially mimic follicular regulatory T cells (Tfr) by providing suppression of Tfh-GC B cell interactions. Supporting this concept, a recent study demonstrated that Tregs retrovirally transduced with CD19 CAR preferentially localized near B-cell follicles, contributed to the Tfr pool, and suppressed Tfh and GC B-cell responses without measurable B-cell depletion in a murine cGVHD model (*64*).Consistent with this concept, CAR19 EngTreg therapy restored the Tfh/Tfr ratio in SLE123 recipients to levels present in healthy controls. Further, a key feature of both nTreg (*65, 66*) and EngTreg (*67*) is the capacity to promote tissue remodeling and repair. Mouse EngTreg express high-levels of ST2 (the receptor for IL-33; essential for programming tissue repair) and promote tissue healing in inflammatory models including acute kidney injury (*68*). Consistent with these attributes, EngTreg treatment reduced the incidence and severity of inflammatory lesions in both the humanized pristane-induced inflammatory model and the IMQ-induced cutaneous inflammation mouse model. In the Sle123 model, CAR19 mEngTreg improved combinatorial renal injury scores, consistent with reduced active renal inflammation and attenuation of progressive tissue injury.

Despite demonstrated efficacy of CAR19 EngTreg across in vitro and in lupus-relevant models, several limitations remain to be addressed in support of clinical translation. First, CAR19 EngTreg manufacturing is complex, requiring dual-locus gene editing to deliver both a CAR19 and a synthetic IL-2 CISC. Transition to clinical manufacturing will require detailed measurements of on- and off-target nuclease activity, efficiency and product purity. Second, while our data demonstrate differential activity of CAR19 EngTreg vs. CAR19 Teff in short-term and humanized settings, future head-to-head comparisons of CAR19 EngTreg with B cell ablative technologies (including CAR19 Teff, BiTEs, and mAbs) in lupus models will be important for clearly defining relative therapeutic impact and the risk for genetic-driven relapse. In previous studies, CAR19 Teff failed to maintain tolerance in mice expressing a human SLE TLR7 gain-of function variant (*69*) and, in our studies, CAR19 EngTreg exhibited less benefit in the TLR-7 driven, SLE1b.Yaa model in comparison to the multigenic SLE123 lupus model, highlighting the challenge of restoring tolerance in the presence of strong genetic drivers. Third, while our data support the conclusion that CAR19 EngTreg mediate therapeutic benefit via antigen-directed immunosuppression, the suppressive mechanisms are likely multifactorial and remain to be determined in more detail. Fourth, CISC mediated, cell intrinsic, IL2 signaling represents a key feature of DE CAR19 EngTreg allowing for rapamycin-based enrichment during manufacturing and improved in vivo engraftment and function in the absence of conditioning (*38, 39*). Here, because surrogate mEngTreg lacked engineered IL-2 support, low dose IL-2 was used to partially mimic CISC-driven in vivo benefits. While we observed no benefit from IL-2 and Cy alone in the control Sle123 cohort, the capacity for cell intrinsic, EngTreg support suggest that our modeling may underestimate the activity of human DE CAR19 EngTreg in future clinical settings. Finally, as with any immunosuppressive modality, evaluation of how CAR19 EngTreg impact protective immunity will be important. Notably, in prior studies, polyclonal EngTreg exhibited no impact on primary or secondary humoral responses or EBV viral challenge (*70*). Additional work, however, will be required to assess the potential impact of CAR19 EngTreg in vaccine responsiveness and antiviral immunity. Together, these limitations highlight important avenues for future investigation.

In summary, these studies suggest that CD19-directed EngTreg may be capable of efficiently mediating long-term immune tolerance and limiting end-organ injury by constraining both GC and EF responses and remodeling inflammatory microenvironments. Together, our findings provide a compelling foundation for advancing CAR19 EngTreg toward clinical development in SLE and other autoimmune disorders.

## MATERIALS AND METHODS

### Study Design

The objective of this study was to test whether CD19-directed engineered regulatory T cells (CAR19 EngTreg), generated by HDR at *FOXP3* coupled with delivery of a CD19 CAR, could provide durable suppression of pathogenic B and T cell responses in systemic lupus erythematosus (SLE). We first generated and characterized human CAR19 EngTreg products and assessed their phenotype, activation, cytokine production, cytotoxicity, and ability to suppress B cell differentiation and antibody production in vitro using primary human cells and established co-culture systems. Where possible, in vitro data were derived from at least three biological replicates using independent donors, with technical replicates performed within individual experiments. We next evaluated CAR19 EngTreg in vivo using multiple complementary models. In humanized mice, we assessed B cell targeting, T cell expansion, and inflammatory cytokine production following adoptive transfer of engineered human T cells. In a pristane-induced humanized lupus model, we assessed the ability of CAR19 EngTreg to reduce local and systemic inflammation relative to CAR19 Teff and anti-CD19 monoclonal antibody treatment. In parallel, murine surrogate CAR19 EngTreg were generated and tested in immune-competent models, including a short-term imiquimod-induced inflammatory model and the B6.Sle123 lupus model, where endpoint analyses included immunophenotyping, autoantibody production, and renal pathology. Endpoints and sample sizes for in vivo studies were predetermined based on prior experience with these models and published literature. Investigators were not blinded to treatment allocation unless otherwise noted. Animals were assigned to experimental groups following confirmation of model establishment or engraftment, and data were collected at predefined timepoints, including interim bleed collections and terminal endpoints. Inclusion criteria for in vitro and in vivo experiments were established based on successful cell product generation or model establishment, respectively. Outliers were not excluded unless attributable to technical failure. Sample sizes, number of biological and technical replicates, and statistical methods are provided in the figure legends.

### Primary human T cells

Human PBMCs were procured from Fred Hutchinson Cooperative Center for Excellence in Hematology Cell Processing Core Facility or purchased from STEMCELL Technologies. CD4+ T cells were isolated from PBMCs using the EasySep Human CD4+ T Cell Enrichment Kit (STEMCELL Technologies). Isolated CD4+ T cells were either cryopreserved or cultured immediately for downstream editing experiments. Methods used for T cell culturing, editing, and LV transduction have been described previously (*35*) and are summarized in Supplementary Materials and Methods.

### Flow Cytometry

Tables S1 and S2 list all antibodies and viability dyes used for flow cytometry, including fluorophore, clone, and vendor information. For surface staining, cells were incubated with viability dye and fluorophore-conjugated antibodies for 30 min at 4°C in the dark. For intracellular staining, cells were first stained for surface markers, followed by fixation and permeabilization using either BD Cytofix/Cytoperm or the BioLegend True-Nuclear Transcription Factor Buffer Set according to the manufacturer’s instructions. Cells were then stained with antibodies against intracellular targets for 30 min at 4°C in the dark. Samples were acquired on either FACSymphony A5 or LSRFortessa flow cytometers (BD Biosciences) and analyzed using FlowJo Version 10.

### Incucyte Assay

For Incucyte killing assays, we used GFP-expressing target (CD19.K562.GFP) cells and incubated with relevant T cells at 1:1 and 1:2 E/T ratio at 37°C for up to 3 days. Four images were recorded per well every 2 h and were analyzed with the Incucyte image analysis software.

### Cytokine profile assessment of CAR19 EngTreg

Cryopreserved CAR19 EngTreg, mock-transduced cells, polyclonal EngTreg, and TCR EngTreg were thawed and rested for 3 days in media supplemented with IL-2 (5 ng/mL; PeproTech). Cells were then stimulated with either anti-CD3/anti-CD28 Dynabeads (Thermo Fisher Scientific) at a 1:1 bead-to-cell ratio, primary human B cells, or NALM6 cells for 16 hours at 37°C. For intracellular cytokine detection, GolgiPlug and GolgiStop (BD Biosciences) were added during the final 6 hours of stimulation according to the manufacturer’s instructions. Cells stimulated with anti-CD3/anti-CD28 beads, primary B cells, or NALM6 cells were stained with antibodies against LAP (BioLegend) and GARP (BD Biosciences). Cells treated with GolgiPlug and GolgiStop were additionally stained intracellularly with antibodies against IFN-γ, TNF-α, and IL-2 (BioLegend) and analyzed by flow cytometry as described above. Supernatants collected from cocultures with primary B cell stimulation were stored at −80°C and subsequently analyzed using the LEGENDplex Human CD8/NK 13-plex panel (BioLegend) according to the manufacturer’s instructions.

### Bead-based and antigen specific T cell suppression assays

Bead-based (antigen-agnostic) suppression assays were performed as previously described (*34*). Percentage suppression calculated as [(%) Teff dividing without Treg – (%) Teff dividing with Treg) / (%) Teff dividing without Treg] x 100. In a separate experiment, an antigen-specific T effector suppression assay was performed. Suppressor T cell products (CAR19 EngTreg, polyclonal EngTreg, or mock-transduced polyclonal Teff cells) were thawed and rested in low-dose IL-2 for 2 days prior to labeling with CFSE and coculture with CellTrace Violet (CTV)-labeled primary B cells for 18 hours. In parallel, responder T cells consisting of autologous CD4+ Teff cells were thawed, rested, and stimulated with anti-CD3/anti-CD28 Dynabeads for 18 hours at 37°C. Following stimulation, responder cells were washed, depleted of Dynabeads, labeled with eFluor670, and added to the suppressor T cell and primary B cell cocultures. After 4 days of coculture, responder T cell proliferation was assessed by eFluor670 dilution using flow cytometry. Percent suppression was calculated using the following formula: suppression (%) = ([a − b]/a) × 100, where a represents the percentage of responder T cell proliferation in the absence of CAR19 EngTreg and b represents the percentage of responder T cell proliferation in the presence of CAR19 EngTreg.

### B cell suppression assay

Primary human B cells were isolated from PBMCs and activated in vitro to induce differentiation toward antibody-secreting cells using a previously described method of differentiating plasma cells in vitro (*42*). Detailed culture conditions, cytokine compositions, and co-culture timelines are provided in the Supplementary Materials and Methods. Briefly, CAR19-EngTreg and control cell populations (including PFN1-KO CAR19 EngTreg, polyclonal EngTreg, and CAR19 Teff) were activated prior to co-culture and introduced at defined T:B ratios during either early (day 2) or late (day 7) stages of B cell differentiation. Co-cultures were maintained through a staged cytokine-driven differentiation system promoting plasmablast and plasma cell development. B cell responses were assessed longitudinally by flow cytometry, and total IgG production was quantified from culture supernatants at the terminal timepoint by ELISA.

### ELISA

Human IgM and total IgG levels in culture supernatants and serum collected from humanized mice were quantified using the Human IgM Uncoated ELISA Kit and Human IgG Uncoated ELISA Kit (Thermo Fisher Scientific) according to the manufacturer’s instructions.

For congenic mouse studies, anti-dsDNA and anti-Sm/RNP autoantibodies were measured by indirect ELISA. Briefly, 96-well NUNC plates were first coated with 0.005% poly-L-lysine (Sigma-Aldrich) prior to coating with calf thymus dsDNA (100 μg/mL; Sigma-Aldrich) for anti-dsDNA ELISAs or coated only with Sm/RNP antigen (5 μg/mL; Arotec Diagnostics) for anti-Sm/RNP ELISAs. Plates were incubated with mouse serum samples, followed by detection with HRP-conjugated anti-mouse IgG antibody (Southern Biotech). Serum NGAL levels were quantified using the Mouse Lipocalin-2/NGAL Quantikine ELISA Kit (R&D systems) according to the manufacturer’s instructions. Signals were developed using OptEIA (BD) TMB substrate reagents and optical density quantified using a Spectramax i3x microplate reader.

### Humanized Mouse models

#### NBSGW model

8-week-old NBSGW mice were conditioned using Busulfan (12.5mg/kg, i.p.), followed by delivery of human CD34+ HPSC 24h later (0.8-2 million cells, r.o.). Human immune reconstitution was assessed 14 weeks later by PB flow cytometry. At week 15, mice received CAR19 EngTreg or CAR19 Teff cells (3.5 × 10^6 cells/mouse, i.v.). Body weight was monitored following T cell transfer. PB was collected 4 days post-transfer, and mice were sacrificed 7 days post-transfer for endpoint flow cytometry and serum analysis.

#### NSG-Pristane model

Eight-week-old NSG mice were sub-lethally irradiated and received 0.80*10^5^ human CD34+ cord blood hematopoietic stem cells i.v. (Lonza Bioscience). Human immune reconstitution was monitored weekly in PB every week by flow cytometry. After 13 weeks, mice received 500 μL pristane i.p. (Sigma-Aldrich) except for those designated as healthy controls. After 3 weeks, pristane-injected mice were randomized into treatment groups based on sex and hCD45+ cell count in the periphery. CAR19 EngTreg and CAR19-Teff treated animals received 3.5 x10^6^ engineered cells i.v. Mice in the mAb group received 10 mg/kg of inebilizumab (Uplizna, Horizon Therapeutics), initially i.v. and subsequently i.p. every 2 weeks (*71*). Finally, a group of mice received 200 ml of PBS i.v. (Corning). PB was monitored weekly by flow cytometry. Mice were sacrificed at 4- and 8-week endpoints for flow cytometric and histopathologic analysis of blood and tissues. All animal studies were approved by the Italian Health Ministry Institutional Animal Care and Use Committee (IACUC).

### Cytokine assay

Human cytokines were quantified using the LEGENDplex Human CD8/NK 13-plex panel (BioLegend) according to the manufacturer’s instructions. Samples were acquired by flow cytometry and analyzed using the LEGENDplex Data Analysis Software Suite (BioLegend).

### Pathological Evaluation

At the sacrifice, organs were harvested, fixed with neutral buffered formalin and subsequently embedded in paraffin. Hematoxylin and eosin stained 5 µm paraffin sections were examined for histopathological analysis by board-certified pathologists who were blinded to the treatment groups. Detailed criteria for evaluation are described in the Supplementary Methods and Materials.

### Primary mouse T cell isolation, culture, editing, and enrichment

CD4^+^ T cells were isolated from CD45.1 C57BL/6 splenocytes by negative selection using the EasySep mouse CD4^+^ T Cell Enrichment Kit (STEMCELL) and activated using murine anti-CD3/CD28–coated beads (Gibco Thermo Fisher Scientific) for 40-48 hours in RPMI-1640 containing 20% fetal bovine serum (Omega Scientific Inc.), 10mM HEPES, 2mM Glutamax, 55µM β-mercaptoethanol, and mouse IL-2 (50 ng/ml; PeproTech). After activation, cells were separated from beads, further cultured for approx. 16 hours in media, then washed twice in phosphate-buffered saline (PBS) and resuspended in MaxCyte Buffer at 1.25×10^8^ cells/ml. RNP was prepared by mixing 40 pmol of Cas9 (Integrated DNA Technologies, IDT) with 100 pmol of mouse *Foxp3*-specific guide RNA for 25 min at room temperature. Delivery of RNP into mouse cells was achieved by electroporation (MaxCyte GTx) followed by incubation with AAV5 donor template for 20-24 hours at 37°C. Cells were supplemented with fresh media containing IL-2 and transferred to a new tissue culture dish for another 20-24 hours before being collected, counted, stained with biotinylated anti-LNGFR antibody (Miltenyi Biotec), and enriched using anti-biotin microbeads (Miltenyi Biotec) as previously described.

### In vivo studies using murine lupus models

B6.Sle1.Sle2.Sle3 (Sle123) and male B6.Sle1yaa (Sle1b.YAA) mice were purchased from The Jackson Laboratory and aged to 7–8 months, a timepoint associated with established autoimmunity and circulating autoantibodies. Mice were conditioned with cyclophosphamide (100 mg/kg; Ingenus Pharmaceuticals) 24 hours prior to adoptive cell transfer. Animals then received 3 × 10^6^ murine CAR19 EngTreg by retro-orbital injection. To model the IL-2 cytokine support provided by the IL-2 CISC in the human CAR19 EngTreg product, mice receiving CAR19 EngTreg and conditioning-only controls animals were treated with low-dose IL-2 as previously described by Rose et al (*72*). PB was collected weekly through 2 weeks post-transfer and biweekly thereafter. Blood samples were used for flow cytometric analysis, and serum was isolated using SST tubes (BD) for downstream ELISA-based autoantibody measurements. Animal studies and procedures were approved by the Institutional Animal Care and Use Committee of SCRI and performed in accordance with the NIH Guide for the Care and Use of Laboratory Animals

### Autoantigen Microarrays

Serum autoantibody profiling was performed by the UT Southwestern Microarray & Immune Phenotyping Core Facility using their autoantigen microarray platform. Briefly, serum samples were screened for IgG autoantibody reactivity against a broad panel of self-antigens associated with systemic autoimmune disease, including lupus-associated autoantigens. Arrays were processed and analyzed by the core facility according to established protocols. Resulting fluorescence intensity values were log_2_-normalized prior to comparative analysis of autoantibody reactivity profiles between treatment groups.

### Statistical Analysis

Statistical analyses were performed using GraphPad Prism version 10 unless otherwise indicated. Where error bars are present, data are shown as mean ± SD or mean ± SEM, as indicated in the figure legends. Sample sizes (*n*) represent biological replicates, defined as individual mice or as independent donors where applicable, and are reported in the corresponding figure legends. For comparisons between two groups, unpaired two-tailed Student’s *t* tests were used where indicated, and Welch’s *t* tests were used for comparisons between two groups with unequal variance. For comparisons among more than two groups, one-way ANOVA was used. For experiments involving two independent variables, two-way ANOVA with correction for multiple comparisons was used. Repeated measures two-way ANOVA was used for longitudinal analyses of repeated measurements from the same animals over time. Tukey or Dunnett post-hoc tests for multiple comparisons were used following ANOVA where indicated. Histopathologic data scored by the presence or absence of inflammatory lesions was treated as categorical data and analyzed using Fisher’s exact test, using RStudio. Summed histopathology scores were subsequently analyzed in RStudio using Kruskal-Wallis tests followed by Dunn’s post hoc testing with false discovery rate adjustment for multiple comparisons. All statistical tests were two-sided unless otherwise stated. Multiple-comparison adjustments were applied as indicated in the figure legends. Exact statistical tests used for each experiment are reported in the corresponding figure legends. *P* values less than 0.05 were considered statistically significant.

## List of Supplementary Materials

Materials and Methods

Fig. S1 to S13

Tables S1 to S3

References (*75–76*) (numbers for references only cited in SM)

## Acknowledgments

We thank Jennifer Haddock for assistance with manuscript preparation and for administrative support in her role as assistant to DJR. We also thank Laura Smith for project management and for facilitating coordination and communication with collaborators.

## Funding

GentiBio (DJR, CBo)

Seattle Children’s Research Institute Program for Cell and Gene Therapy (DJR)

Children’s Guild Association Endowed Chair in Pediatric Immunology (DJR)

Hansen Investigator in Pediatric Innovation Endowment (DJR)

## Author contributions

Conceptualization: TD, VDN, CM, CP, TC, ADRE, CBo, TW, PJC, DJR

Methodology: TD, MD, AG, VDN, JM, YC, PJC

Investigation: TD, MD, AG, VDN, MH, BM, ST, JM, MM, PZ, AB, PC, AU, CBe, AV, NPD

Formal analysis: TD, MD, AG, MH, PJC, SA

Visualization: TD, AG, VDN, MH, JM, PZ, AB, PJC

Resources: YC, PK, PC, AU, CBe, AV, NPD

Funding acquisition: CP, CBo, TW, DJR

Project administration: CM, CP, TC, ADRE, CBo, TW, PJC, DJR

Supervision: CBo, TW, PJC, DJR

Writing – original draft: TD, PJC, DJR

Writing – review & editing: TD, MD, VDN, MH, PZ, CBo, PJC, DJR

## Competing interests

VDN, MH, JM, MM, PZ, AB, CM, CP, TC, ADRE, and TW are current or former employees of GentiBio. SWJ is a consultant for Amgen, Bristol Myers Squibb, Merck, and Palleon Pharmaceuticals and previously served as a consultant for IgM Biosciences and Sail Biomedicines. CBo is an inventor on different patents on cancer immunotherapy and genetic engineering. CBo has been a member of the Advisory Board and a consultant for Molmed, Intellia, TxCell, Novartis, GSK, Allogene, Kite/Gilead, Miltenyi, Kiadis, Evir, and Janssen and received research support from Molmed s.p. and Intellia Therapeutics. PJC and DJR are inventors on a patent describing methods for generating antigen-specific engineered regulatory T cells. DJR is a scientific cofounder and former scientific advisory board member of GentiBio and received funding from GentiBio for related studies and from CSL Behring, Be Biopharma, and Emendo Bio for unrelated work.

## Data and materials availability

All data are available in the main text or the supplementary materials. Additional data may be requested and is available upon request.

## References and Notes

1. M. R. Arbuckle, M. T. McClain, M. V. Rubertone, R. H. Scofield, G. J. Dennis, J. A. James, J. B. Harley, Development of autoantibodies before the clinical onset of systemic lupus erythematosus. The New England Journal of Medicine 349, 1526–1533 (2003).

2. G. Biesecker, S. Katz, D. Koffler, Renal localization of the membrane attack complex in systemic lupus erythematosus nephritis. J Exp Med 154, 1779–1794 (1981).

3. A. Rahman, D. A. Isenberg, Systemic lupus erythematosus. N Engl J Med 358, 929–939 (2008).

4. L. Arnaud, F. Chasset, T. Martin, Immunopathogenesis of systemic lupus erythematosus: An update. Autoimmun Rev 23, 103648 (2024).

5. P. Shen, S. Fillatreau, Antibody-independent functions of B cells: a focus on cytokines. Nature Reviews. Immunology 15, 441–451 (2015).

6. D. Ghosh, W. Jiang, D. Mukhopadhyay, E. D. Mellins, New insights into B cells as antigen presenting cells. Current Opinion in Immunology 70, 129–137 (2021).

7. R. A. Elsner, M. J. Shlomchik, Germinal Center and Extrafollicular B Cell Responses in Vaccination, Immunity, and Autoimmunity. Immunity 53, 1136–1150 (2020).

8. S. A. Jenks, K. S. Cashman, E. Zumaquero, U. M. Marigorta, A. V. Patel, X. Wang, D. Tomar, M. C. Woodruff, Z. Simon, R. Bugrovsky, E. L. Blalock, C. D. Scharer, C. M. Tipton, C. Wei, S. S. Lim, M. Petri, T. B. Niewold, J. H. Anolik, G. Gibson, F. E. Lee, J. M. Boss, F. E. Lund, I. Sanz, Distinct Effector B Cells Induced by Unregulated Toll-like Receptor 7 Contribute to Pathogenic Responses in Systemic Lupus Erythematosus. Immunity 49, 725–739 e726 (2018).

9. N. Seki, H. Tsujimoto, S. Tanemura, S. Kojima, F. Miyoshi, J. Kikuchi, S. Saito, M. Akiyama, K. Sugahara, K. Yoshimoto, Y. Kaneko, K. Chiba, T. Takeuchi, Cytotoxic Tph subset with low B-cell helper functions and its involvement in systemic lupus erythematosus. Commun Biol 7, 277 (2024).

10. Y. Atisha-Fregoso, B. Toz, B. Diamond, Meant to B: B cells as a therapeutic target in systemic lupus erythematosus. The Journal of Clinical Investigation 131, e149095 (2021).

11. R. Furie, B. H. Rovin, F. Houssiau, A. Malvar, Y. K. O. Teng, G. Contreras, Z. Amoura, X. Yu, C.-C. Mok, M. B. Santiago, A. Saxena, Y. Green, B. Ji, C. Kleoudis, S. W. Burriss, C. Barnett, D. A. Roth, Two-Year, Randomized, Controlled Trial of Belimumab in Lupus Nephritis. The New England Journal of Medicine 383, 1117–1128 (2020).

12. S. W. Jackson, A. Davidson, BAFF inhibition in SLE-Is tolerance restored? Immunological Reviews 292, 102–119 (2019).

13. J. T. Merrill, C. M. Neuwelt, D. J. Wallace, J. C. Shanahan, K. M. Latinis, J. C. Oates, T. O. Utset, C. Gordon, D. A. Isenberg, H.-J. Hsieh, D. Zhang, P. G. Brunetta, Efficacy and safety of rituximab in moderately-to-severely active systemic lupus erythematosus: the randomized, double-blind, phase II/III systemic lupus erythematosus evaluation of rituximab trial. Arthritis Rheum 62, 222–233 (2010).

14. E. G. Kamburova, H. J. P. M. Koenen, K. J. E. Borgman, I. J. ten Berge, I. Joosten, L. B. Hilbrands, A single dose of rituximab does not deplete B cells in secondary lymphoid organs but alters phenotype and function. American Journal of Transplantation: Official Journal of the American Society of Transplantation and the American Society of Transplant Surgeons 13, 1503–1511 (2013).

15. D. Mougiakakos, G. Kronke, S. Volkl, S. Kretschmann, M. Aigner, S. Kharboutli, S. Boltz, B. Manger, A. Mackensen, G. Schett, CD19-Targeted CAR T Cells in Refractory Systemic Lupus Erythematosus. N Engl J Med 385, 567–569 (2021).

16. A. Mackensen, F. Muller, D. Mougiakakos, S. Boltz, A. Wilhelm, M. Aigner, S. Volkl, D. Simon, A. Kleyer, L. Munoz, S. Kretschmann, S. Kharboutli, R. Gary, H. Reimann, W. Rosler, S. Uderhardt, H. Bang, M. Herrmann, A. B. Ekici, C. Buettner, K. M. Habenicht, T. H. Winkler, G. Kronke, G. Schett, Anti-CD19 CAR T cell therapy for refractory systemic lupus erythematosus. Nat Med 28, 2124–2132 (2022).

17. C. Tur, M. Eckstein, J. Velden, S. Rauber, C. Bergmann, J. Auth, L. Bucci, G. Corte, M. Hagen, A. Wirsching, R. Grieshaber-Bouyer, P. Reis, N. Kittan, J. Wacker, A. Rius Rigau, A. Ramming, M. A. D’Agostino, A. Hartmann, F. Muller, A. Mackensen, A. Bozec, G. Schett, M. G. Raimondo, CD19-CAR T-cell therapy induces deep tissue depletion of B cells. Ann Rheum Dis 84, 106–114 (2025).

18. F. Muller, J. Taubmann, L. Bucci, A. Wilhelm, C. Bergmann, S. Volkl, M. Aigner, T. Rothe, I. Minopoulou, C. Tur, J. Knitza, S. Kharboutli, S. Kretschmann, I. Vasova, S. Spoerl, H. Reimann, L. Munoz, R. G. Gerlach, S. Schafer, R. Grieshaber-Bouyer, A. S. Korganow, D. Farge-Bancel, D. Mougiakakos, A. Bozec, T. Winkler, G. Kronke, A. Mackensen, G. Schett, CD19 CAR T-Cell Therapy in Autoimmune Disease - A Case Series with Follow-up. N Engl J Med 390, 687–700 (2024).

19. A. Nordmann-Gomes, L. Khalili, W. Tang, L. Geraldino-Pardilla, Y. Gartshteyn, R. Clancy, M. Souvignier, S. Suh, M. Sadelain, A. Askanase, CAR T-cell therapy in SLE: A systematic review. Semin Arthritis Rheum 74, 152786 (2025).

20. D. Suan, J. Moore, C. C. Goodnow, Can autoimmune disease be cured by deep CD19+ cell depletion? J Immunol 214, 1075–1092 (2025).

21. D. J. Rawlings, G. Metzler, M. Wray-Dutra, S. W. Jackson, Altered B cell signalling in autoimmunity. Nat Rev Immunol 17, 421–436 (2017).

22. S. W. Jackson, N. S. Kolhatkar, D. J. Rawlings, B cells take the front seat: dysregulated B cell signals orchestrate loss of tolerance and autoantibody production. Current Opinion in Immunology 33, 70–77 (2015).

23. D. J. Rawlings, M. A. Schwartz, S. W. Jackson, A. Meyer-Bahlburg, Integration of B cell responses through Toll-like receptors and antigen receptors. Nature Reviews. Immunology 12, 282–294 (2012).

24. C. E. Faliti, M. Ghimire, M. Garcia-Vega, R. C. Watermeier, A. Callahan, J. Burke, O. Posadas, A. K. Mishra, S. Khurana, V. Greiff, C. D. Scharer, J. M. Lindner, R. G. King, M. Newell, A. Khosroshahi, F. E. Lee, I. Sanz, Defective B cell tolerance in SLE lymph nodes underpins VH (4-34) “clonal damnation” and PD-1(+)TOX(+) autoreactive B cells expansion. medRxiv, (2026).

25. C. W. Freyer, D. L. Porter, Cytokine release syndrome and neurotoxicity following CAR T-cell therapy for hematologic malignancies. J Allergy Clin Immunol 146, 940–948 (2020).

26. J. N. Brudno, J. N. Kochenderfer, Current understanding and management of CAR T cell-associated toxicities. Nat Rev Clin Oncol 21, 501–521 (2024).

27. J. A. Bluestone, M. Anderson, Tolerance in the Age of Immunotherapy. The New England Journal of Medicine 383, 1156–1166 (2020).

28. T. S. Sumida, N. T. Cheru, D. A. Hafler, The regulation and differentiation of regulatory T cells and their dysfunction in autoimmune diseases. Nat Rev Immunol 24, 503–517 (2024).

29. C. M. Wardell, D. A. Boardman, M. K. Levings, Harnessing the biology of regulatory T cells to treat disease. Nat Rev Drug Discov 24, 93–111 (2025).

30. J. A. Bluestone, B. K. Burnett, C. E. Crute, M. D. Fellows, M. Levings, H. Lebrec, L. Pereira Mouriès, J. Rice, P. Rohan, M. G. Roncarolo, D. Rotrosen, Regulatory T cell therapies to treat autoimmune diseases and transplant rejection. Nature Immunology 26, 819–824 (2025).

31. J. H. Buckner, Antigen-specific immunotherapies for autoimmune disease. Nature Reviews. Rheumatology 21, 88–97 (2025).

32. D. A. Boardman, M. K. Levings, Emerging strategies for treating autoimmune disorders with genetically modified Treg cells. J Allergy Clin Immunol 149, 1–11 (2022).

33. L. M. R. Ferreira, Y. D. Muller, J. A. Bluestone, Q. Tang, Next-generation regulatory T cell therapy. Nat Rev Drug Discov 18, 749–769 (2019).

34. Y. Honaker, N. Hubbard, Y. Xiang, L. Fisher, D. Hagin, K. Sommer, Y. Song, S. J. Yang, C. Lopez, T. Tappen, E. M. Dam, I. Khan, M. Hale, J. H. Buckner, A. M. Scharenberg, T. R. Torgerson, D. J. Rawlings, Gene editing to induce FOXP3 expression in human CD4(+) T cells leads to a stable regulatory phenotype and function. Sci Transl Med 12, (2020).

35. S. J. Yang, A. K. Singh, T. Drow, T. Tappen, Y. Honaker, F. Barahmand-Pour-Whitman, P. S. Linsley, K. Cerosaletti, K. Mauk, Y. Xiang, J. Smith, E. Mortensen, P. J. Cook, K. Sommer, I. Khan, D. Liggitt, D. J. Rawlings, J. H. Buckner, Pancreatic islet-specific engineered T(regs) exhibit robust antigen-specific and bystander immune suppression in type 1 diabetes models. Sci Transl Med 14, eabn1716 (2022).

36. G. I. Uenishi, M. Repic, J. Y. Yam, A. Landuyt, P. Saikumar-Lakshmi, T. Guo, P. Zarin, M. Sassone-Corsi, A. Chicoine, H. Kellogg, M. Hunt, T. Drow, R. Tewari, P. J. Cook, S. J. Yang, K. Cerosaletti, D. Schweinoch, B. Guiastrennec, E. James, C. Patel, T. F. Chen, J. H. Buckner, D. J. Rawlings, T. J. Wickham, K. T. Mueller, GNTI-122: an autologous antigen-specific engineered Treg cell therapy for type 1 diabetes. JCI Insight 9, (2024).

37. S. K. Tripathi, A. Grimm, N. P. Dahl, Y. Honaker, P. Knebusch, Y. Chen, P. J. Cook, D. J. Rawlings, HLA-A2 CAR/IL-2-CISC engineered Treg display robust in vitro and in vivo antigen-specific regulatory function. Mol Ther Methods Clin Dev 33, 101561 (2025).

38. M. S. Hunt, S. J. Yang, E. Mortensen, A. Boukhris, J. Buckner, P. J. Cook, D. J. Rawlings, Dual-locus, dual-HDR editing permits efficient generation of antigen-specific regulatory T cells with robust suppressive activity. Mol Ther 31, 2872–2886 (2023).

39. P. J. Cook, S. J. Yang, G. I. Uenishi, A. Grimm, S. E. West, L.-J. Wang, C. Jacobs, A. Repele, T. Drow, A. Boukhris, N. P. Dahl, K. Sommer, A. M. Scharenberg, D. J. Rawlings, A chemically inducible IL-2 receptor signaling complex allows for effective in vitro and in vivo selection of engineered CD4+ T cells. Molecular Therapy: The Journal of the American Society of Gene Therapy 31, 2472–2488 (2023).

40. R. Y. Cheng, K. L. Hung, T. Zhang, C. M. Stoffers, A. R. Ott, E. R. Suchland, N. D. Camp, I. F. Khan, S. Singh, Y. J. Yang, D. J. Rawlings, R. G. James, Ex vivo engineered human plasma cells exhibit robust protein secretion and long-term engraftment in vivo. Nat Commun 13, 6110 (2022).

41. X. Rui, F. Alvarez Calderon, H. Wobma, U. Gerdemann, A. Albanese, L. Cagnin, C. McGuckin, K. A. Michaelis, K. Naqvi, B. R. Blazar, V. Tkachev, L. S. Kean, Human OX40L-CAR-Tregs target activated antigen-presenting cells and control T cell alloreactivity. Science Translational Medicine 16, eadj9331 (2024).

42. K. L. Hung, I. Meitlis, M. Hale, C. Y. Chen, S. Singh, S. W. Jackson, C. H. Miao, I. F. Khan, D. J. Rawlings, R. G. James, Engineering Protein-Secreting Plasma Cells by Homology-Directed Repair in Primary Human B Cells. Mol Ther 26, 456–467 (2018).

43. B. E. McIntosh, M. E. Brown, B. M. Duffin, J. P. Maufort, D. T. Vereide, Slukvin, II, J. A. Thomson, Nonirradiated NOD,B6.SCID Il2rgamma-/-Kit(W41/W41) (NBSGW) mice support multilineage engraftment of human hematopoietic cells. Stem Cell Reports 4, 171–180 (2015).

44. M. Gunawan, Z. Her, M. Liu, S. Y. Tan, X. Y. Chan, W. W. S. Tan, S. Dharmaraaja, Y. Fan, C. B. Ong, E. Loh, K. T. E. Chang, T. C. Tan, J. K. Y. Chan, Q. Chen, A Novel Human Systemic Lupus Erythematosus Model in Humanised Mice. Sci Rep 7, 16642 (2017).

45. M. Doglio, A. Ugolini, C. Bercher-Brayer, B. Camisa, C. Toma, R. Norata, S. Del Rosso, R. Greco, F. Ciceri, F. Sanvito, M. Casucci, A. A. Manfredi, C. Bonini, Regulatory T cells expressing CD19-targeted chimeric antigen receptor restore homeostasis in Systemic Lupus Erythematosus. Nat Commun 15, 2542 (2024).

46. J. H. Stone, A. Khosroshahi, W. Zhang, E. Della Torre, K. Okazaki, Y. Tanaka, J. M. Lohr, N. Schleinitz, L. Dong, H. Umehara, M. Lanzillotta, Z. S. Wallace, M. Ebbo, G. J. Webster, F. Martinez Valle, M. K. Nayar, C. A. Perugino, V. Rebours, X. Dong, Y. Wu, Q. Li, N. Rampal, D. Cimbora, E. L. Culver, M. T. Investigators, Inebilizumab for Treatment of IgG4-Related Disease. N Engl J Med 392, 1168–1177 (2025).

47. M. L. Davila, C. C. Kloss, G. Gunset, M. Sadelain, CD19 CAR-targeted T cells induce long-term remission and B Cell Aplasia in an immunocompetent mouse model of B cell acute lymphoblastic leukemia. PLoS One 8, e61338 (2013).

48. M. Souyris, C. Cenac, P. Azar, D. Daviaud, A. Canivet, S. Grunenwald, C. Pienkowski, J. Chaumeil, J. E. Mejia, J. C. Guery, TLR7 escapes X chromosome inactivation in immune cells. Sci Immunol 3, (2018).

49. G. J. Brown, P. F. Canete, H. Wang, A. Medhavy, J. Bones, J. A. Roco, Y. He, Y. Qin, J. Cappello, J. I. Ellyard, K. Bassett, Q. Shen, G. Burgio, Y. Zhang, C. Turnbull, X. Meng, P. Wu, E. Cho, L. A. Miosge, T. D. Andrews, M. A. Field, D. Tvorogov, A. F. Lopez, J. J. Babon, C. A. Lopez, A. Gonzalez-Murillo, D. C. Garulo, V. Pascual, T. Levy, E. J. Mallack, D. G. Calame, T. Lotze, J. R. Lupski, H. Ding, T. R. Ullah, G. D. Walters, M. E. Koina, M. C. Cook, N. Shen, C. de Lucas Collantes, B. Corry, M. P. Gantier, V. Athanasopoulos, C. G. Vinuesa, TLR7 gain-of-function genetic variation causes human lupus. Nature 605, 349–356 (2022).

50. R. R. Goel, X. Wang, L. J. O’Neil, S. Nakabo, K. Hasneen, S. Gupta, G. Wigerblad, L. P. Blanco, J. B. Kopp, M. I. Morasso, S. V. Kotenko, Z. X. Yu, C. Carmona-Rivera, M. J. Kaplan, Interferon lambda promotes immune dysregulation and tissue inflammation in TLR7-induced lupus. Proc Natl Acad Sci U S A 117, 5409–5419 (2020).

51. L. van der Fits, S. Mourits, J. S. A. Voerman, M. Kant, L. Boon, J. D. Laman, F. Cornelissen, A.-M. Mus, E. Florencia, E. P. Prens, E. Lubberts, Imiquimod-induced psoriasis-like skin inflammation in mice is mediated via the IL-23/IL-17 axis. Journal of Immunology (Baltimore, Md.: 1950) 182, 5836–5845 (2009).

52. L. Morel, B. P. Croker, K. R. Blenman, C. Mohan, G. Huang, G. Gilkeson, E. K. Wakeland, Genetic reconstitution of systemic lupus erythematosus immunopathology with polycongenic murine strains. Proceedings of the National Academy of Sciences of the United States of America 97, 6670–6675 (2000).

53. A. De Groof, P. Hémon, O. Mignen, J.-O. Pers, E. K. Wakeland, Y. Renaudineau, B. R. Lauwerys, Dysregulated Lymphoid Cell Populations in Mouse Models of Systemic Lupus Erythematosus. Clin Rev Allergy Immunol 53, 181–197 (2017).

54. L. Morel, B. P. Croker, K. R. Blenman, C. Mohan, G. Huang, G. Gilkeson, E. K. Wakeland, Genetic reconstitution of systemic lupus erythematosus immunopathology with polycongenic murine strains. Proc Natl Acad Sci U S A 97, 6670–6675 (2000).

55. H. Zhu, H. Luo, M. Yan, X. Zuo, Q. Z. Li, Autoantigen Microarray for High-throughput Autoantibody Profiling in Systemic Lupus Erythematosus. Genomics Proteomics Bioinformatics 13, 210–218 (2015).

56. B. P. Croker, G. Gilkeson, L. Morel, Genetic interactions between susceptibility loci reveal epistatic pathogenic networks in murine lupus. Genes Immun 4, 575–585 (2003).

57. S. Bolivar-Wagers, M. L. Loschi, S. Jin, G. Thangavelu, J. H. Larson, C. S. McDonald-Hyman, E. G. Aguilar, A. Saha, B. H. Koehn, M. Hefazi, M. J. Osborn, M. C. Jensen, J. E. Wagner, C. A. Pennell, B. R. Blazar, Murine CAR19 Tregs suppress acute graft-versus-host disease and maintain graft-versus-tumor responses. JCI insight 7, e160674 (2022).

58. Y. Imura, M. Ando, T. Kondo, M. Ito, A. Yoshimura, CD19-targeted CAR regulatory T cells suppress B cell pathology without GvHD. JCI Insight 5, (2020).

59. Y. Bulliard, R. Freeborn, M. J. Uyeda, D. Humes, R. Bjordahl, D. de Vries, M. G. Roncarolo, From promise to practice: CAR T and Treg cell therapies in autoimmunity and other immune-mediated diseases. Front Immunol 15, 1509956 (2024).

60. M. Tekguc, J. B. Wing, M. Osaki, J. Long, S. Sakaguchi, Treg-expressed CTLA-4 depletes CD80/CD86 by trogocytosis, releasing free PD-L1 on antigen-presenting cells. Proc Natl Acad Sci U S A 118, (2021).

61. S. Glatigny, B. Hollbacher, S. J. Motley, C. Tan, C. Hundhausen, J. H. Buckner, D. Smilek, S. J. Khoury, L. Ding, T. Qin, J. Pardo, G. T. Nepom, L. A. Turka, K. M. Harris, D. J. Campbell, E. Bettelli, Abatacept Targets T Follicular Helper and Regulatory T Cells, Disrupting Molecular Pathways That Regulate Their Proliferation and Maintenance. J Immunol 202, 1373–1382 (2019).

62. S. D. Reighard, S. A. Cranert, K. M. Rangel, A. Ali, I. E. Gyurova, A. T. de la Cruz-Lynch, J. A. Tuazon, M. V. Khodoun, L. C. Kottyan, D. F. Smith, H. I. Brunner, S. N. Waggoner, Therapeutic Targeting of Follicular T Cells with Chimeric Antigen Receptor-Expressing Natural Killer Cells. *Cell Reports*. Medicine 1, 100003 (2020).

63. A. Murai, M. Iwata, S. Miyakawa, D. Warude, M. Sagara, Y. Kikukawa, CXCR5-targeted chimeric antigen receptor T regulatory cells for the selective inhibition of follicular helper T cell and B cell interaction. Cytotherapy 27, 571–580 (2025).

64. S. Jin, M. C. Zaiken, C. McDonald-Hyman, C. R. Hartigan, S. Bolivar-Wagers, J. H. Larson, Y. Peng, S. Hani, M. Riddle, A. Saha, A. Panoskaltsis-Mortari, E. Ko, Y. Zhao, R. Amaro Marquez, P. S. Marri Baskar, C. R. Eide, W. J. Murphy, K. L. Hippen, G. R. Hill, J. Tolar, P. T. Sage, C. A. Pennell, L. S. Kean, B. R. Blazar, CAR19 Tregs treat murine chronic graft-versus-host disease through immune suppression without measurable B cell cytolysis. JCI Insight 11, (2026).

65. C. Campbell, A. Rudensky, Roles of Regulatory T Cells in Tissue Pathophysiology and Metabolism. Cell Metab 31, 18–25 (2020).

66. L. F. Loffredo, T. M. Savage, O. R. Ringham, N. Arpaia, Treg-tissue cell interactions in repair and regeneration. J Exp Med 221, (2024).

67. M. E. Hoover, Milaszewski, M., N. R. Reed, Doherty, A., Gaddis, D.E., Repic, M., Kancharla, A., Wickham, T., Chen, T.F., Zarin, P., Adoptive Transfer of Allogeneic Murine Engineered T Regulatory Cells Ameliorates Disease in Models of Acute Kidney Injury in Mice. Molecular Therapy 32, 41 (2024).

68. P. Zarin, M. Hoover, D. Gaddis, M. Repic, T. F. Chen, GNTI-823: Engineered Allogeneic Tissue Regulatory T Cells Ameliorate AKI via Immune Modulation and Tissue Repair: FR-PO0163. Journal of the American Society of Nephrology 36, 10.1681/ASN.2025c1683aswtsk (2025).

69. A. Grenov, J. Yoon, D. M. Snell, A. Mikolajczak, H. Wang, E. Gürel, A. R. Ronchel, G. Yegen, L. Lee, B. A. Esen, A. Rahman, P. Maciocia, C. G. Vinuesa, Autoantibody origins in lupus and in relapse post CAR-T therapy. bioRxiv, 2025.2010.2020.683393 (2025).

70. Y. Honaker, Dahl, N.P., Singh, S., Singh, A., Lopez, C., Gruber, D., Scharffenberger, S.C., Patel, S., Jones, H., Khan, I.F., Christian, M.L., Yang, S., Sommer, K., Torgerson, T.R., Allenspach, E.J., Ito, C.E.K., Uenishi, G.I., Cost, G.J., Cook, P.J., Rawlings, D.J., Preclinical efficacy and safety assessment of engineered regulatory T cells (EngTreg) for treatment of IPEX and other autoimmune disorders. Molecular Therapy, Submitted (2026).

71. S. Gallagher, I. Yusuf, T. M. McCaughtry, S. Turman, H. Sun, R. Kolbeck, R. Herbst, Y. Wang, MEDI-551 Treatment Effectively Depletes B Cells and Reduces Serum Titers of Autoantibodies in Mice Transgenic for Sle1 and Human CD19. Arthritis Rheumatol 68, 965–976 (2016).

72. A. Rose, C. von Spee-Mayer, L. Kloke, K. Wu, A. Kuhl, P. Enghard, G. R. Burmester, G. Riemekasten, J. Y. Humrich, IL-2 Therapy Diminishes Renal Inflammation and the Activity of Kidney-Infiltrating CD4+ T Cells in Murine Lupus Nephritis. Cells 8, (2019).

73. S. Bolivar-Wagers, J. H. Larson, S. Jin, B. R. Blazar, Cytolytic CD4(+) and CD8(+) Regulatory T-Cells and Implications for Developing Immunotherapies to Combat Graft-Versus-Host Disease. Front Immunol 13, 864748 (2022).

74. M. Hale, B. Lee, Y. Honaker, W. H. Leung, A. E. Grier, H. M. Jacobs, K. Sommer, J. Sahni, S. W. Jackson, A. M. Scharenberg, A. Astrakhan, D. J. Rawlings, Homology-Directed Recombination for Enhanced Engineering of Chimeric Antigen Receptor T Cells. Mol Ther Methods Clin Dev 4, 192–203 (2017).

75. G. I. Uenishi, M. Repic, J. Y. Yam, A. Landuyt, P. Saikumar-Lakshmi, T. Guo, P. Zarin, M. Sassone-Corsi, A. Chicoine, H. Kellogg, M. Hunt, T. Drow, R. Tewari, P. J. Cook, S. J. Yang, K. Cerosaletti, D. Schweinoch, B. Guiastrennec, E. James, C. Patel, T. F. Chen, J. H. Buckner, D. J. Rawlings, T. J. Wickham, K. T. Mueller, GNTI-122: an autologous antigen-specific engineered Treg cell therapy for type 1 diabetes. JCI insight 9, e171844 (2024).

76. O. Chan, M. P. Madaio, M. J. Shlomchik, The roles of B cells in MRL/lpr murine lupus. Ann N Y Acad Sci 815, 75–87 (1997).

