## Supplementary Materials for "CD19 CAR Engineered T Regulatory Cells Promote Durable Immune Tolerance in Experimental Systemic Lupus Erythematosus"

### Plasmid constructs

Candidate human CD19-binding domains, including the FMC63 and Hu19 scFv, were subcloned into multiple chimeric antigen receptor (CAR) architectures and subsequently transferred into lentiviral (LV) and adeno-associated viral (AAV) donor plasmids by Gibson Assembly for generation of human CAR19 EngTreg. Candidate human CAR constructs were screened in vitro prior to use in the described study. For murine studies, a 1D3-based murine CD19 CAR was subcloned into a previously described Foxp3-targeted HDR donor template for generation of mouse CAR19 EngTreg (*35*).

### Viral vector production

Recombinant adeno-associated viruses (AAVs) were produced and titrated using the triple transfection method with a serotype 6 helper plasmid, as previously described (*73*). Vesicular stomatitis virus glycoprotein (VSV-G) pseudotyped lentiviruses were prepared following established protocols (*74*). The titers of recombinant viruses were quantified by quantitative PCR (qPCR) using primers specific for the inverted terminal repeat (ITR) sequences in AAV or the woodchuck hepatitis virus posttranscriptional regulatory element (WPRE) in lentiviruses.

### Human T cell culture, transduction, and genome editing

T cells were cultured in T cell media composed of RPMI 1640 supplemented with 20% fetal bovine serum (FBS), 10 mM HEPES, 2 mM GlutaMAX, and 55 μM β-mercaptoethanol, along with 50 ng/mL human interleukin-2 (IL-2). Activation of T cells was performed using Human T-Expander CD3/CD28 Dynabeads (Thermo Fisher Scientific). Following activation, cells underwent lentiviral transduction for CD19CAR expression and CRISPR/Cas9-mediated gene editing. Parallel cultures were used to generate mock-edited T effector cells (Teff). Polyclonal EngTreg cells were generated by CRISPR editing without LV transduction. CD19CAR Teff cells were generated by LV transduction without CRISPR editing. Post-editing, polyclonal EngTreg and CD19CAR EngTreg were recovered in T cell media supplemented with 50 ng/mL IL-2 for 3 days. Enrichment of IL-2 CISC expressing cells was achieved by culturing in T cell media containing 10 nM rapamycin (Sigma) for an additional 4 days. At day 6 post-editing, cells were reactivated using CD3/CD28 Dynabeads and seeded in 6-well plates fitted with G-Rex gas-permeable culture systems (Wilson Wolf Manufacturing) at a density of 1–5 × $10^6$ cells per well. Cells were expanded in rapamycin-containing media for 7–10 days, with media changes every second day. For dual-HDR editing, the CAR19 EngTreg editing strategy was adopted from methods developed in the Rawlings lab and previously at GentiBio (*38, 75*). $CD4^+$ T cells were isolated via magnetic enrichment from PBMCs (STEMcell) and activated via α-CD3/α-CD28 microbead stimulation (Thermo Fisher Scientific) at a ratio of 3 beads to 1 cell. After activation, cells were gene edited by electroporating 2 RNPs; each consisting of Cas9 protein (Aldevron) and sgRNA (IDT) targeting either the *FOXP3* or *TRAC* locus; transgene templates carrying MND-CISCg-CAR19 (Hu19.IgG4.CD8.CD28.CD3z)-dFRB-FOXP3 for the *FOXP3* loci and MND-CISCb-BGHpA for the *TRAC* loci were delivered via adeno-associated virus serotype 6 (AAV6). In some iterations, the CAR19 was FMC63.CD28.CD28.CD3z. Successfully dual-engineered cells were enriched and expanded with rapamycin and a secondary activation step, and cryopreserved (Cryostor CS10, STEMcell) after purity exceeded 75%.

Mock-edited Teff and CD19CAR Teff were expanded in T cell media supplemented with 50 ng/mL IL-2 but without rapamycin.

### Direct killing assay

Cryopreserved CAR19 EngTreg, Mock-edited control cells, and CAR19 T-effectors were thawed and rested for three days in low IL-2 (5 ng/ml) prior to co-culture with $CD19^+$ Nalm-6 or $CD19^-$ K562 cell lines. Cells were stained after 48 hours with Annexin V and Live/Dead to determine cell killing. Annexin V buffer was obtained from BD Biosciences (BD Pharmingen Annexin V Apoptosis Detection Kit). Cells treated with Staurosporine (Millipore Sigma) served as a cell death control. Cell death was determined by target cells staining positively for Annexin V, Live/Dead, or both markers. Live cells were cells staining negative for both markers.

#### B cell differentiation assay – expanded culture conditions

B cells were isolated from PBMCs using the EasySep Human B Cell Isolation Kit (STEMCELL Technologies) on day 0 and cultured at 37 °C in B cell media consisting of IMDM supplemented with 10% FBS, 10 mM HEPES, 2 mM GlutaMAX, and 5 μM β-mercaptoethanol. Media was further supplemented with 1 μg/mL CpG ODNs (InvivoGen), 1 μg/mL multimeric CD40L (Adipogen) and 40 ng/mL IL-21 (Peprotech) (Phase 1 cytokine mix). On day –2, $CD4^+$ human T cells (CAR19 EngTreg, PFN-1 K/O CAR19 EngTreg, Polyclonal EngTreg, CAR19 Teff) were thawed and activated with α-CD3/α-CD28 microbeads for three days. After stimulation, T cells were rested overnight in T cell media supplemented with 50 ng/mL IL-2. T cell populations were then cocultured on day 2 with activated B cells at 1:1 or 1:5 T:B cell ratios in B cell media supplemented with Phase 1 cytokine mix (phase I intervention). A portion of B cells was reserved and cultured alone for 7 days. On day 3, additional T cells were thawed and stimulated as described above, then cocultured with the reserved B cells on day 7 in B cell media supplemented with 50 ng/mL IL-2 (Peprotech), 50 ng/mL IL-10 (Peprotech), 50 ng/mL IL-6 (Peprotech), and 40 ng/ml IL-21 (Phase 2 cytokine mix). On day 7, co-culture samples from phase I were harvested, pelleted, and resuspended in B cell media supplemented with Phase 2 cytokine mix. On day 10, media from all samples was exchanged with B cell media supplemented with 50ng/ml IL-6, 40ng/ml IL-15, and 15 ng/mL IFN-α 2B (Proteintech) (Phase 3 cytokine mix). Cocultures were maintained until day 13. Input flow cytometry was performed on B and T cell products on days 2 and 7. Coculture outcomes were assessed by flow cytometry on days 3, 7, 10, and 13. Media supernatants were collected on day 13 for total human IgG by ELISA (Invitrogen).

#### Pathological Evaluation – Humanized Mouse Studies

The presence of inflammatory lesions and the presence of human immune cells were assessed by two board-certified pathologists in a double-blinded fashion. For each organ, we evaluated:

- Spleen: the extension of the red and white pulp, their relative proportion and presence of human immune cells and inflammatory lesions
- Lungs: the anatomy, the presence of immune cells and inflammatory lesions in the parenchyma and blood vessels
- Liver: the anatomy, the presence of immune cells, fibrosis and inflammatory lesions
- Kidney: the number of glomeruli and their characteristics, the presence of immune cells and inflammatory lesions

#### Genetic Model - Lupus Nephritis Scoring Methods

Kidneys were scored according to previously published methods (*76*), with modifications. H&E-stained tissues were evaluated via light microscopy by a board-certified veterinary pathologist. If two kidneys were present in section, the most severely affected kidney was scored according to the following criteria:

#### Glomerulus Diameter Measurement

Glomerular diameter was used as an indication of the severity of glomerulonephritis. Larger average diameters generally coincided with increased glomerular disease. The diameters of twenty representative glomeruli, including the capsule (if Bowman's space is not dilated) along the outer cortex were measured, and a mean was determined for each sample.

#### Glomerular Crescent Score

Glomerular crescents were characterized by hypertrophy and hyperplasia of visceral and/or parietal epithelial cells forming a layer within Bowman's space of two or more cells thick. Epithelial cells could exhibit increased basophilia, cytoplasmic vacuolation, and nuclear crowding and could be accompanied by glomerular compression, deformation, synechiae, and/or periglomerular fibrosis. The percentage of glomeruli with crescents was estimated from evaluation of a minimum of five representative 40x fields and a minimum of 50 glomeruli in areas of affected parenchyma, and a score assigned as follows:

| | |
|---|---|
| 0 | Normal, no crescents |
| 0.5 | Very minimal,1% of glomeruli have crescents |
| 1 | Minimal, 2-4% of glomeruli have crescents |
| 2 | Mild, 5-10% of glomeruli have crescents |
| 3 | Moderate, 11-25% of glomeruli have crescents |
| 4 | Marked, 26-50% of glomeruli have crescents |
| 5 | Severe, >50% of glomeruli have crescents |

#### Protein Cast Score

Protein casts were characterized by hyaline, eosinophilic, homogeneous fluid within renal tubules. Protein casts could be accompanied by tubular dilation, degeneration, granular casts, and/or regeneration, which were not captured in this score but generally trended in severity with protein casts. The number of cortical tubules with protein casts was estimated and a score assigned as follows:

| | |
|---|---|
| 0 | Normal tubules |
| 0.5 | Very minimal, affects 1-2% of cortex |
| 1 | Minimal, affects 3-10% of cortex |
| 2 | Mild, multifocal tubules with protein casts affecting 11-25% of cortex |
| 3 | Moderate, multifocal tubules with protein casts affecting 26-50% of cortex |
| 4 | Marked, multifocal tubules with protein casts affecting 51-75% of cortex |
| 5 | Severe, diffuse tubules with protein casts affecting greater than 75% of cortex |

#### Interstitial Infiltrates Score

Interstitial infiltrates were characterized by the infiltration of inflammatory cells, primarily mononuclear cells, into the renal pelvis and cortex or in other areas not directly associated with perivascular spaces. The extent of inflammatory cell infiltrates within the cortex and pelvis was estimated and a score assigned as follows:

| | |
|---|---|
| 0 | Normal, no interstitial inflammation |
| 0.5 | Very minimal, small focal area of inflammatory cell infiltrate generally localized to the pelvis |
| 1 | Minimal, occasional small focal accumulations of inflammatory cells affecting less than 10% of total interstitium and generally localized to the pelvis |
| 2 | Mild, multifocal small to larger infiltrates distributed around the pelvis and cortex, affecting 10-25% of the interstitial area |
| 3 | Moderate, multifocal small to extensive infiltrates in the pelvis and cortex, affecting 26-50% of the interstitial area |

| | |
|---|---|
| 4 | Marked, multifocal to diffuse infiltration affecting 51-75% of the interstitial area of the pelvis and cortex |
| 5 | Severe, diffuse infiltration affecting 76-100% of the interstitial area of the pelvis and cortex |

**Perivascular Infiltrates Score**

Perivascular infiltrates were characterized by the cuffing of vessels with inflammatory cells, primarily mononuclear cells, leading to expansion of the perivascular space; this could be accompanied by degeneration of the associated vascular wall. The extent and severity of inflammatory cell infiltrates in perivascular spaces was estimated and a score assigned as follows:

| | |
|---|---|
| 0 | Normal, no perivascular infiltrates |
| 0.5 | Very minimal, a single focal vessel with minimal perivascular infiltrate |
| 1 | Minimal, small but definite perivascular infiltrates (generally 1-2 affected vessels) |
| 2 | Mild, multifocal foci of perivascular infiltrates (generally 3-4 affected vessels); no necrosis of vessel walls is present |
| 3 | Moderate, multifocal foci of perivascular infiltrates (generally 5-6 affected vessels); may have minimal necrosis of vessel walls |
| 4 | Marked, multifocal foci of perivascular infiltrates (generally 7-8 affected vessels); may have some necrosis of vessel walls |
| 5 | Severe, multifocal foci of perivascular infiltrates (generally >8 affected vessels); necrosis of some vessel walls is present |

**Summed Score**

The scores obtained for crescent, protein cast, interstitial infiltrate, and perivascular infiltrate were summed. The evaluation of the glomerular diameter was not included in the sum score.

**Spleen**

The spleen was evaluated for splenic white pulp characteristics. The total number of white pulp follicles is counted, and the largest diameter follicle is measured in cross section (µm).

**IMQ Model Studies**

*Mice*: Wild-type 7–9-week-old male C57BL/6 mice (The Jackson Laboratory, #000664) were treated topically with the TLR7 agonist imiquimod cream (5%, Taro) or Vaseline to the dorsal and ventral aspects of right ear skin 3 times weekly until the indicated timepoint based on study. Mice were monitored and ear thickness was measured 3x a week using a caliper. When indicated, murine IL-2 (PeproTech, 2ug) was dosed by intraperitoneal injection 2x a week starting on Day -1. cy(Millipore Sigma, 50mg/kg – 200mg/kg) was administered via intraperitoneal injection on Day -1. Local inflammation was scored based on ear tissue thickness, acanthosis, and skin scaling. Mice were sacrificed 48 hours after the final imiquimod treatment and blood was collected by terminal cardiac puncture. Spleens were harvested and processed for flow cytometry.

*Adoptive cell transfer*: 3x$10^6$ CAR19 engineered Treg or CAR19 CD4 T conventional cells were injected intravenously into gender-matched 7-9 week old mice as indicated. 5 days following transfer, frequencies of donor-derived cells in the blood of recipient mice were analyzed by flow cytometry.

*Tissue harvest and analysis*: Blood was collected into EDTA microtainer tubes (BD), washed in PBS, lysed with ACK Lysis Buffer (Gibco), washed in 10% FBS-RPMI, and brought up in PBS for staining. Spleens were mashed through a 40μM sterile filter to obtain single-cell suspensions and washed with 10% FBS-RPMI prior to staining.

**Table S1: Antibodies used for human in vitro and humanized mouse studies**

| Antibody target/marker | Fluorochrome | Supplier | Catalog Number | Clone |
|---|---|---|---|---|
| 2A peptide | PE | Sigma | MABS2005 | 3H4 |
| CAR19 (FMC63) | APC | ACROBiosystems | FM3-AY54P1 | Y45 |
| | FITC | Miltenyi Biotec | 130-127-344 | REA1297 |
| Whitlow/218 Linker | AF488 | Cell Signaling Technology | 55809 | E3U7Q |
| | PE | Cell Signaling Technology | 62405L | E3U7Q |
| CD3 | APC-Cy7 | BioLegend | 300318 | HIT3a |
| | FITC | BD Biosciences | 349201 | SK7 |
| | eFluor506 | Invitrogen | 69-0037-42 | OKT3 |
| | BUV395 | BD Biosciences | 563646 | UCHT1 |
| CD4 | AF700 | BioLegend | 344622 | SK3 |
| | AF700 | BioLegend | 300526 | RPA-T4 |
| | FITC | BD Biosciences | 555346 | RPA-T4 |
| CD8 | PerCP-Cy5.5 | BD Biosciences | 560662 | RPA-T8 |
| | BV786 | BioLegend | 344740 | SK1 |
| CD69 | BV605 | BioLegend | 310938 | FN50 |
| | BV650 | BioLegend | 310934 | FN50 |
| CD137 | PE | BD Biosciences | 555956 | 4B4-1 |
| FOXP3 | PE | BioLegend | 320208 | 259D |
| | AF488 | BioLegend | 320212 | 259D |
| CTLA-4 | APC | BioLegend | 349908 | L3D10 |
| | BV605 | Biolegend | 369610 | BNI3 |
| ICOS | AF700 | BioLegend | 313528 | C398.4A |
| | BUV496 | BD Biosciences | 750321 | DX29 |
| CD127 | PE-CF594 | BD Biosciences | 562397 | HIL-7R-M21 |
| CD25 | PE-Cy7 | BD Biosciences | 335789 | 2A3 |

| | | | | |
|---|---|---|---|---|
| | BV785 | BioLegend | 356140 | M-A251 |
| LNGFR | FITC | Miltenyi biotec | 130-113-420 | ME20.4-1.H4 |
| LAP | PE_Cy7 | BioLegend | 300007 | S20006A |
| GARP | PE | BioLegend | 142903 | F011-5 |
| | BV421 | BioLegend | 352509 | 7B11 |
| IFN-gamma | AF647 | BioLegend | 502516 | 4S.B3 |
| | BV711 | BioLegend | 502539 | 4S.B3 |
| TNF-alpha | APC-Cy7 | BioLegend | 502944 | MAb11 |
| IL-2 | BV605 | BioLegend | 500332 | MQ1-17H12 |
| IL-6 | PE-Cy7 | Biolegend | 501120 | MQ2-13A5 |
| CD19 | APC | BioLegend | 982406 | HIB19 |
| | FITC | BioLegend | 392508 | 4G7 |
| | BUV563 | BD Biosciences | 741361 | HIB19 |
| | BUV615 | BD Biosciences | 741361 | HIB19 |
| CD38 | PerCP-Cy5.5 | BD Biosciences | 551400 | HIT2 |
| CD138 | PE | BioLegend | 356504 | MI15 |
| | PE-Cy5 | Beckman Coulter | A54191 | B-A38 |
| IgM | PE-Cy7 | BioLegend | 314532 | MHM-88 |
| IgG | BV786 | BD Biosciences | 564230 | G18-145 |
| Ig light chain Kappa | BV421 | BioLegend | 392706 | TB28-2 |
| CD45 | APC-Cy7 | BioLegend | 368516 | 2D1 |
| | PerCP | BioLegend | 304026 | HI30 |
| | PE-eFlour610 | Invitrogen | 61-9459-42 | 2D1 |
| CD56 | PE | BioLegend | 362508 | 5.1H11 |
| CD14 | APC-Cy7 | BioLegend | 367108 | 63D3 |
| CD20 | PE-Cy7 | BioLegend | 302312 | 2H7 |
| | BUV805 | BD Biosciences | 612905 | 2H7 |
| CD206 | APC | BioLegend | 321110 | 15-2 |
| CD27 | APC-Cy7 | BioLegend | 356424 | M-T271 |
| HLA-DR | BUV805 | BD Biosciences | 568335 | G46-6 |
| PD1 | BV650 | BioLegend | 329950 | EH12.2H7 |
| CD163 | BV605 | BD Biosciences | 745091 | GHI/61 |
| TNFR2 | PE-TR | Biolegend | 358414 | 3G7A02 |
| 4-1BB | APC | Biolegend | 309810 | 4B4-1 |

| | | | | |
|---|---|---|---|---|
| CD127 | APC/Fire 750 | Biolegend | 986004 | A019D5 |
| Perforin | BV421 | Biolegend | 308122 | dG9 |
| Granzyme B | PE | Biolegend | 372208 | QA16A02 |
| GM-CSF | PE-CF594 | BD Horizon | 562857 | BVD2-21C11 |

**Table S2: Antibodies used in murine EngTreg studies & genetic lupus mouse models**

| Antibody target / marker | Fluorochrome | Supplier | Catalog number | Clone |
|---|---|---|---|---|
| **Viability Dye** | AF-350 NHS Ester | Thermo Fisher | A10168 | -- |
| **CD95** | BV421 | BD Biosciences | 562633 | Jo2 |
| **Blimp-1** | APC 647 | BD Biosciences | 563643 | 5E7 |
| **CXCR5** | PE | BD Biosciences | 551959 | 2G8 |
| **IgD** | BV786 | BD Biosciences | 563618 | 11-26c.2a |
| **CXCR5** | PE-Cy7 | BD Biosciences | 560617 | 2G8 |
| **IL-10** | BV711 | BD Biosciences | 564081 | JES5-16E3 |
| **CD11b** | FITC | BD Biosciences | 553310 | M1/70 |
| **CD3** | BUV395 | BD Biosciences | 569614 | 17A2 |
| **BCL-6** | BUV661 | BD Biosciences | 568062 | K112-91 |
| **CD127** | BUV737 | BD Biosciences | 612841 | SB/199 |
| **CD39** | BV421 | BD Biosciences | 567105 | Y23-1185 |
| **CTLA-4** | APC-R700 | BD Biosciences | 565778 | UC10-4F10-11 |
| **CD69** | BUV496 | BD Biosciences | 741063 | H1.2F3 |
| **KLRG-1** | BUV563 | BD Biosciences | 741343 | 2F1 |
| **TIM-2** | BUV615 | BD Biosciences | 753150 | RMT2-14 |
| **CD45.2** | BUV661 | BD Biosciences | 741516 | 104 |
| **CD44** | BUV805 | BD Biosciences | 741921 | IM7 |
| **PD-1** | BV480 | BD Biosciences | 746784 | J43 |
| **CD73** | BV650 | BD Biosciences | 752735 | TY/11.8 |
| **CD73** | BV785 | BD Biosciences | 752737 | TY/11.8 |
| **Ki67** | BB780 | BD Biosciences | 568761 | B56 |
| **TCF-1** | APC | BD Biosciences | 566693 | S33-966 |
| **Ki67** | BV785 | BD Biosciences | 563756 | B56 |
| **CD43** | BUV737 | BD Biosciences | 612840 | S7 |
| **Ly108** | BUV395 | BD Biosciences | 745730 | 13G3-19D |
| **CD8a** | BUV563 | BD Biosciences | 752637 | 53-6.7 |

| | | | | |
|---|---|---|---|---|
| **CD95** | BUV496 | BD Biosciences | 741103 | Jo2 |
| **CD80** | BUV615 | BD Biosciences | 751328 | 16-10A1 |
| **CD273** | BUV737 | BD Biosciences | 752608 | TY25 |
| **CD23** | BUV805 | BD Biosciences | 741922 | B3B4 |
| **CD24** | BV480 | BD Biosciences | 746709 | M1/69 |
| **CD138** | BV750 | BD Biosciences | 747070 | 281-2 |
| **CD80** | BUV496 | BD Biosciences | 751328 | 16-10A1 |
| **kappa lc** | BUV563 | BD Biosciences | 748800 | 187.1 |
| **BP-1** | BUV615 | BD Biosciences | 752355 | 6C3 |
| **CD93** | BUV661 | BD Biosciences | 741574 | AA4.1 |
| **CD98** | BUV805 | BD Biosciences | 752895 | RL388 |
| **CD1d** | BUV563 | BD Biosciences | 741287 | 1B1 |
| **TACI** | BV421 | BD Biosciences | 742840 | 8F10-3 |
| **TOX / TOX2** | RB780 | BD Biosciences | 570192 | NAN44B |
| **IRF-4** | PE | BD Biosciences | 566646 | Q9-343 |
| **CD11b** | FITC | BD Biosciences | 553310 | M1/70 |
| **CD38** | BB630 | BD Biosciences | 759660 | 90 |
| **CD3** | RB705 | BD Biosciences | 757248 | 17A2 |
| **Slamf-7** | BV421 | BD Biosciences | 747997 | 4G2 |
| **CD21** | APC-Cy7 | BioLegend | 123418 | 7E9 |
| **CD86** | BV605 | BioLegend | 105037 | GL-1 |
| **B220** | BV650 | BioLegend | 103241 | RA3-6B2 |
| **CD4** | BV750 | BioLegend | 100467 | GK1.5 |
| **ICOS** | APC-Cy7 | BioLegend | 313530 | C398.4A |
| **CXCR4** | BV711 | BioLegend | 146517 | L276F12 |
| **Helios** | PerCP | BioLegend | 137230 | 22F6 |
| **T-bet** | PE-CF594 | BioLegend | 644827 | 4B10 |
| **TIGIT** | PE-CF594 | BioLegend | 142110 | 1G9 |
| **LAG-3** | BV650 | BioLegend | 125227 | C9B7W |
| **CXCR3** | BV711 | BioLegend | 155917 | CXCR3-173 |
| **CD38** | PE-Cy7 | BioLegend | 102718 | 90 |
| **CD45.1** | BV605 | BioLegend | 110738 | A20 |
| **CD62L** | BV570 | BioLegend | 104433 | MEL-14 |
| **CD25** | APC-Cy7 | BioLegend | 102026 | PC61 |
| **CCR6** | BV605 | BioLegend | 129819 | 29-2L17 |
| **CD11b** | BV605 | BioLegend | 101257 | M1/70 |

| | | | | |
|---|---|---|---|---|
| **GL-7** | BV421 | BioLegend | 144614 | GL7 |
| **CD19** | PE-Cy5 | BioLegend | 115510 | 6D5 |
| **CD39** | PE/Fire700 | BioLegend | 143820 | Duha59 |
| **CD25** | PE-Cy7 | BioLegend | 101916 | PC61 |
| **CD38** | APC | BioLegend | 165604 | 90 |
| **CD44** | BV570 | BioLegend | 103037 | IM7 |
| **GARP** | PE-Cy7 | BioLegend | 142910 | YGIC86 |
| **BCL-6** | PE/Dazzle 594 | BioLegend | 358510 | 7D1 |
| **TGF-B1 / LAP** | APC | BioLegend | 141406 | TW7-16B4 |
| **Bcl6** | BUV661 | BD Biosciences | 568062 | K112-91 |
| **CD19** | BUV737 | BD Biosciences | 612781 | 1D3 |
| **PD-1** | BV480 | BD Biosciences | 746784 | J43 |
| **CD4** | BV786 | BioLegend | 100453 | GK1.5 |
| **CD3** | FITC | Invitrogen | 11-0032-82 | 17A2 |
| **FOXP3** | APC | Invitrogen | 17-5773-82 | FJK-16s |
| **CD45.2** | AF700 | Invitrogen | 56-0454-82 | 104 |
| **ICOS** | APC-Cy7 | BioLegend | 313529 | C398.4A |
| **CXCR5** | Biotin | BD Biosciences | 551960 | 2G8 |
| **CD45.1** | BV650 | BD Biosciences | 563754 | A20 |
| **LNGFR** | PE-Cy7 | Miltenyi Biotec | 130-125-205 | REA648 |
| **CD8a** | BV510 | BioLegend | 100751 | 53-6.7 |

**Table S3: Guide RNA sequences used for human and mouse gene editing**

| **Genomic Locus** | **Species** | **Target Sequence (5'->3')** |
|---|---|---|
| FOXP3 | Human | TCCAGCTGGGCGAGGCTCCT |
| TRAC | Human | TCTCTCAGCTGGTACACGGC |
| PRF1 | Human | TGTGGCCGGCTCACACTCAC |
| Foxp3 | Mouse | CCAGCUUGGCAAGACUCCUG |
| Rosa26 | Mouse | GGCAGGCTTAAAGGCTAACC |

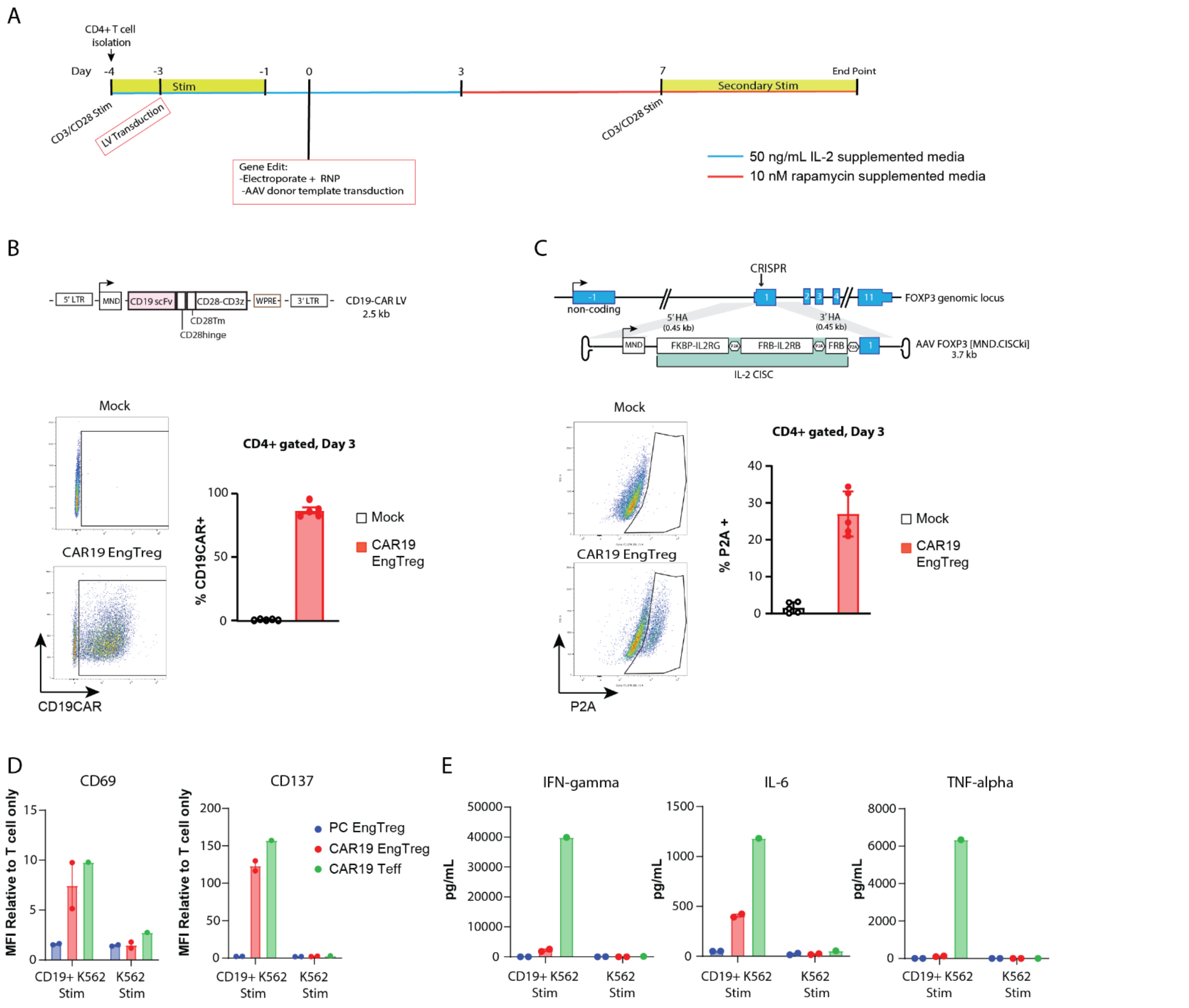


**Fig. S1. Production and characterization of CAR19 EngTreg using an LV+HDR strategy. (A)** Timeline schematic showing CAR19 EngTreg production strategy. (**B**) Schematic of LV vector for CD19 CAR delivery (upper panel) and % transduction on D3 based on flow cytometry for CAR expression (lower panel; tracking FMC63 scFv). (**C**) Schematic of HDR editing strategy at *FOXP3* locus using MND.CISCki AAV donor template (upper panel) and % HDR on D3 post-editing based on P2A epitope staining (lower panel). Bar graphs from 5 experiments representing 3 independent PBMC donors. (**D**) CD69 (left) and CD137 (right) expression on polyclonal EngTreg (n=2 donors), CAR19 EngTreg (n=2 donors), or CAR19 Teff (n=1 donor) in response to 48h co-culture with CD19$^{+}$ or CD19$^{-}$ K562 target cells at 1:1 ratio. Graphed flow cytometry data is normalized to T cell only condition for each cell type. (**E**) From same experiment timepoint, inflammatory cytokine levels in conditioned media as determined by Cytometric Bead Array (CBA) assay.

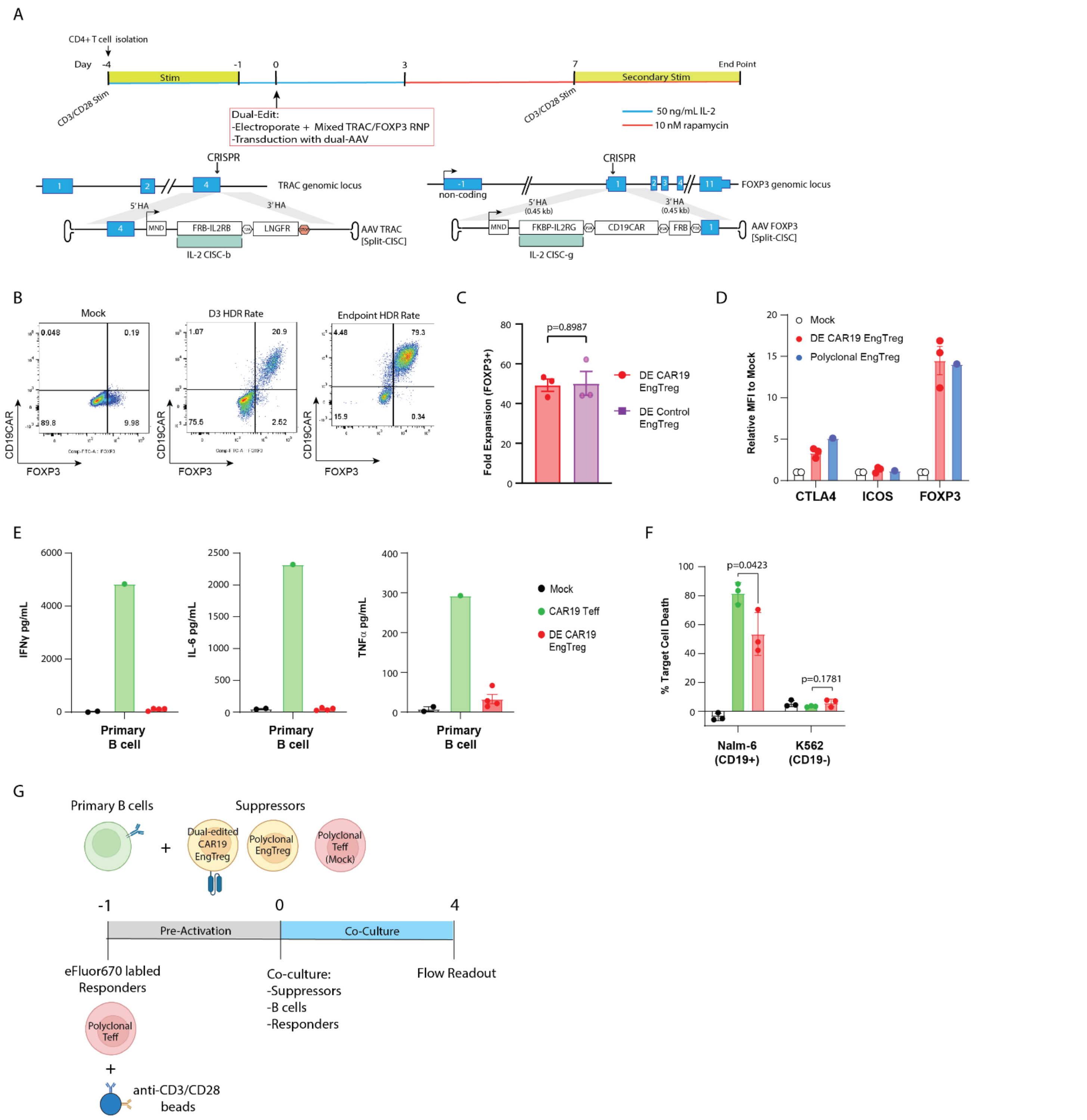


**Fig. S2. Production and characterization of DE CAR19 EngTreg.** (**A**) (*Upper panel*) Timeline schematic of production strategy. (*Lower panels*) Schematic of dual HDR-editing (DE) strategy showing locus-specific HDR donor template and editing outcome for *TRAC* and *FOXP3*, respectively. **B**) Representative flow cytometry of mock-edited vs DE CAR19 EngTreg at D3 and DE CAR19 EngTreg at enrichment endpoint. (**C**) Fold expansion of FOXP3+ cells at enrichment endpoint for DE CAR19 EngTreg or DE Control EngTreg. N=3 independent PBMC donors. (**D**) Immunophenotyping of polyclonal EngTreg (n=1 donors), DE CAR19 EngTreg (n=3 donors), or Mock edited CD4+ T cells (n=2 donors). Graphed data shows relative fold change in MFI of indicated Treg markers relative to Mock. (**E**) Inflammatory cytokine production in DE CAR19 EngTreg, CAR19

Teff, or mock edited T cells in response to B cell targets (CD19$^{mid}$ primary B cells or CD19$^{hi}$ Nalm-6); graphs show cytokine concentration in media supernatant based on CBA assay. (**F**) Flow-based cytotoxicity for stimulated T cell products cultured with a CD19$^{hi}$ Nalm-6 or CD19$^{-}$ K562 1:1 ratio for 48h. (**G**) Schematic for in-vitro antigen specific T cell suppression assay.

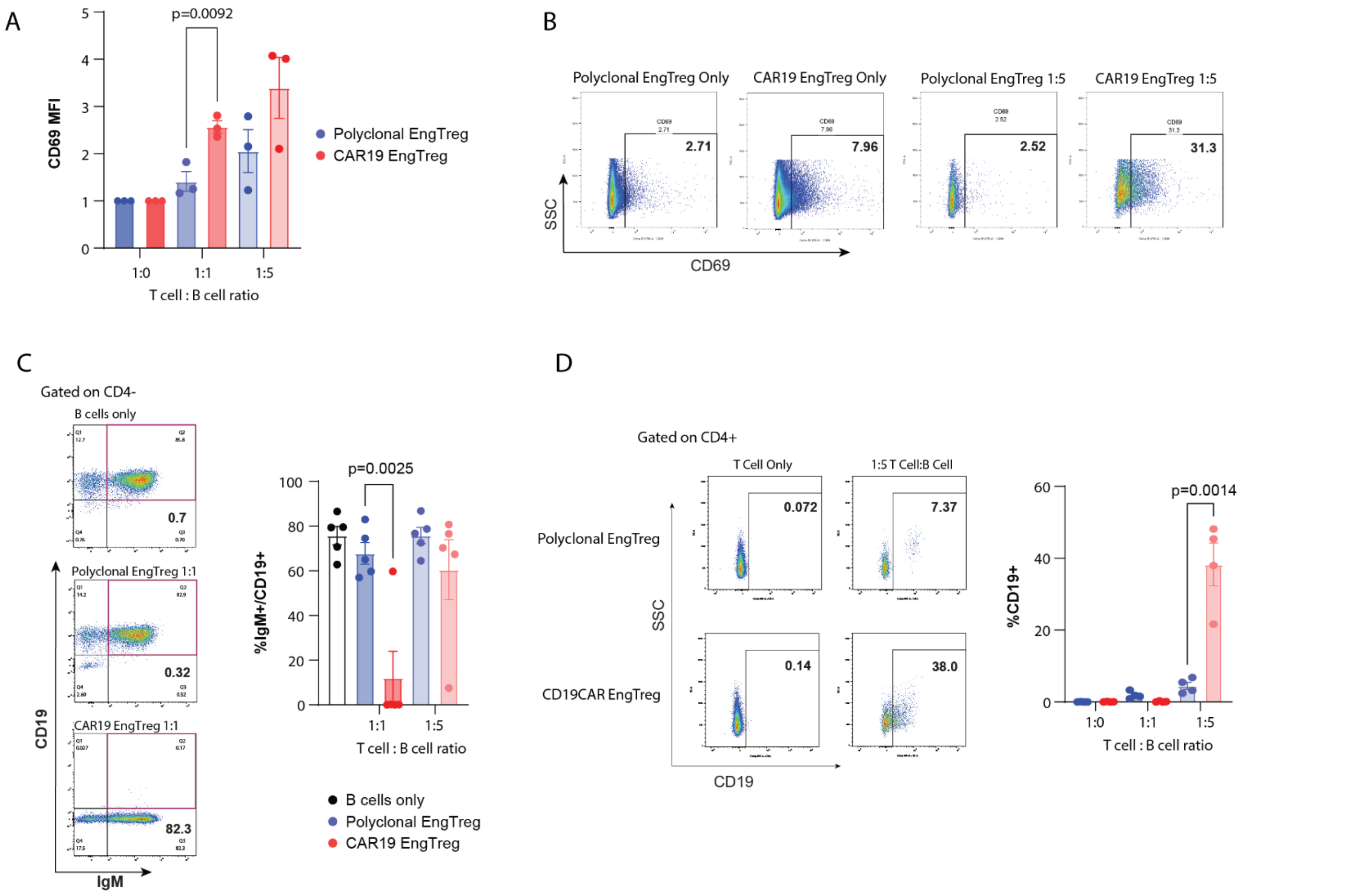


**Fig. S3. Activation and CD19 trogocytosis exhibited by CAR19 EngTreg co-cultured with primary B cells.** (**A**) Quantification of CD69 expression (relative MFI) on polyclonal vs. CAR19 EngTreg on D3, 24 hours after initiation of B cell co-culture at indicated T:B cell ratios. (**B**) Representative flow cytometry showing CD69 expression on polyclonal or CAR19 EngTreg products on D3. (**C**) (*Left panels*) Representative flow cytometry showing IgM vs. CD19 surface expression on co-cultured B cells (gated as CD4- cells) on D3 at 1:1 T:B cell ratio with primary B cells. (*Right panel*) Graph showing proportion of IgM+/CD19+ 'double positive' B cells in cultures containing EngTreg cell products alone or in co-cultures at indicated T:B cell ratio with primary B cells. (**D**) (*Left panels*) Representative flow cytometry for surface CD19 on CD4+ T cell products (demonstrating trogocytosis) cultured alone or in co-culture with primary B cells. (*Right panel*) Graph showing proportion of CD19+ T cells for indicated products. Data are from N=5 donors. Data presented as mean ± SEM. Statistical significance was determined using Welch one-way ANOVA with Dunnett's post-hoc comparisons.

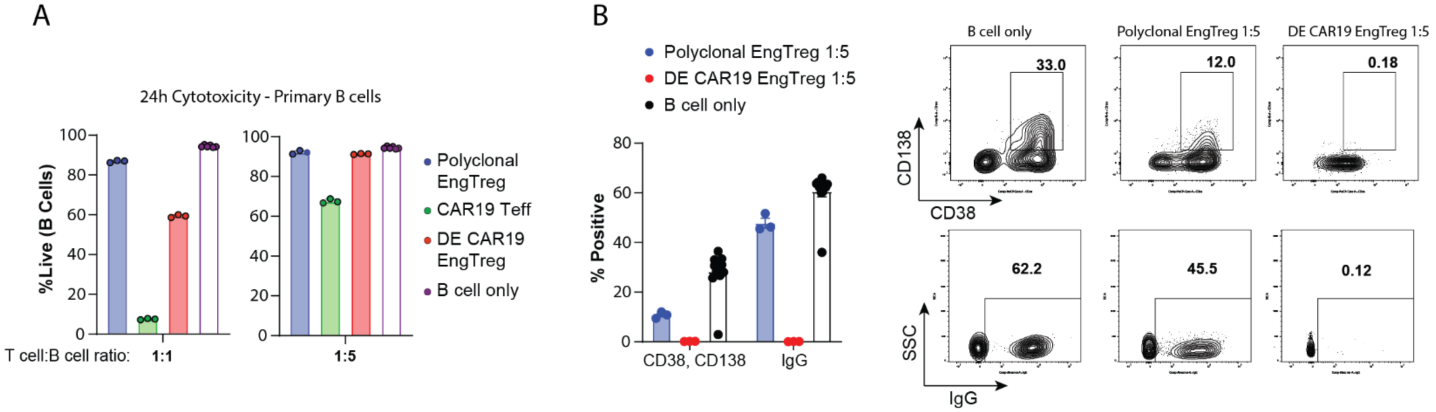


**Fig. S4. Dual-HDR CAR19 EngTreg display antigen-specific suppression toward activated B cells.** (**A**) Viability of activated CD19+ primary B cells based on viability staining 24h after co-culture with indicated T cell products. (**B**) (*Left panel*) PC formation was assessed at Day 13 (based on flow cytometry staining for CD38, CD138 and intracellular IgG) in co-cultures with indicated T cell products. (*Right panels*) Combined data showing % $CD38^{+}CD138^{+}$ PCs and intracellular IgG expression, respectively. Representative flow cytometry data. Data in A-B are from N=1 donor. Data are presented as mean ± SEM.

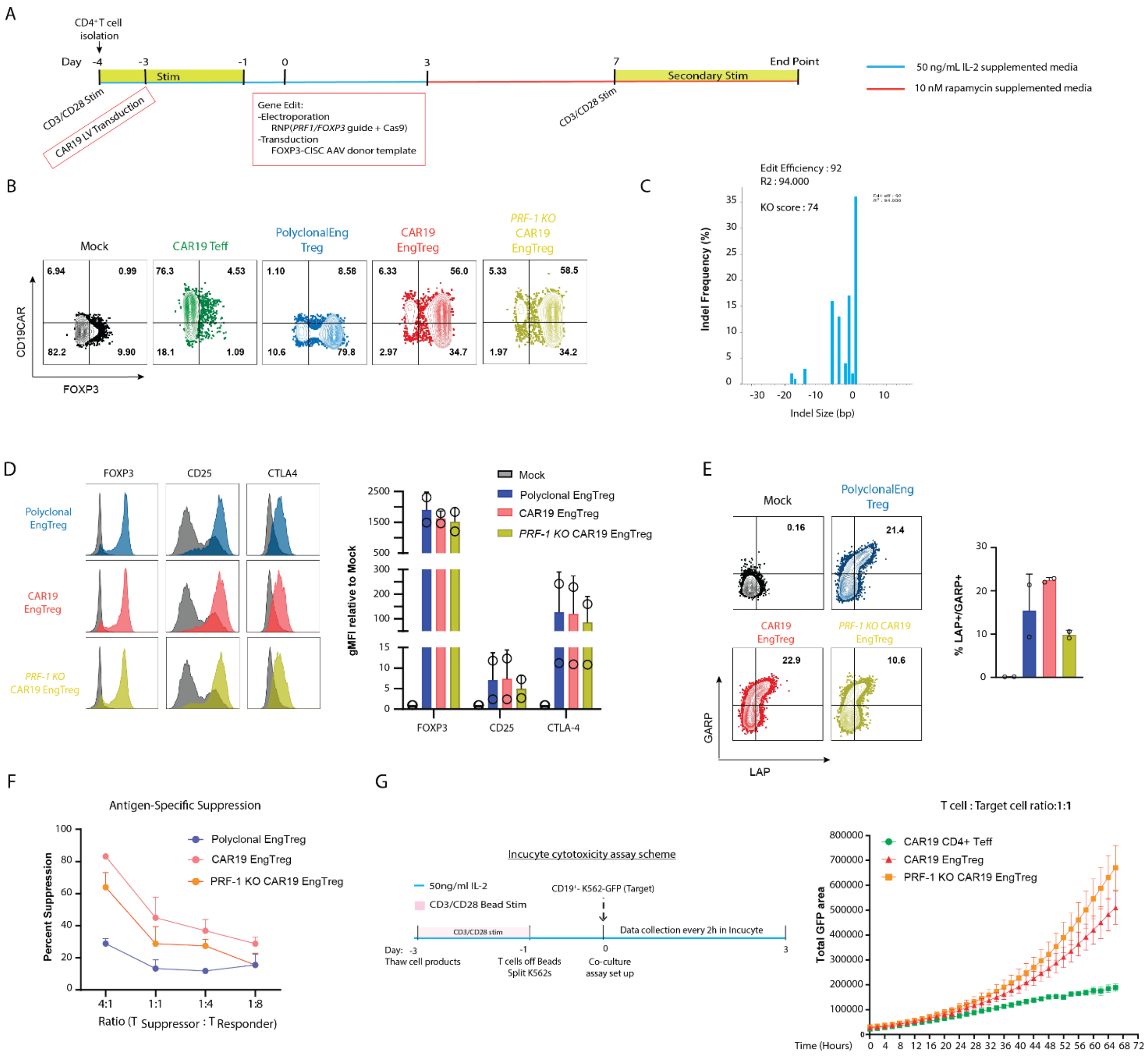


**Fig. S5. Generation and characterization of perforin knock-out CAR19 EngTreg.** (**A**) Schematic and timeline for production and enrichment of PRF-1KO CAR19 EngTreg. (**B**) Representative flow cytometry showing expression of CAR19 and FOXP3 post-enrichment for indicated cell products. (**C**) Inference of CRISPR edits (ICE) analysis of PRF-1 locus at D3 post guide delivery. (**D**) (*Left panels*) Representative flow cytometry analysis and (*right panel*) quantification (based upon MFI) for Treg surface markers for indicated cell products at time of cryopreservation. (**E**) (*Left panels*) Representative flow cytometry analysis and (*right panel*) % TGF-b surface expression (based on LAP and GARP staining) following CD3/CD28 activation for indicated cell products. (**F**) Antigen-specific T cell suppression assay. eFluor670 labeled CAR19+ T responder cells were stimulated with CD3/CD28 and co-cultured with suppressor cells (CAR19 EngTreg, polyclonal EngTreg, or PRF-1 KO CAR19 EngTreg) and CD19+ targets. T responder and T suppressor cells were co-cultured at indicated ratios and eFluor670 dye dilution was measured after 4 days. (**G**) Imaging-based cytotoxicity analysis performed by culturing T cell products with CD19$^{+}$-K562-GFP target cells at 1:1 (left) and 1:2 (right) ratios. Total GFP+ cell area was quantitated continuously for 3 days.

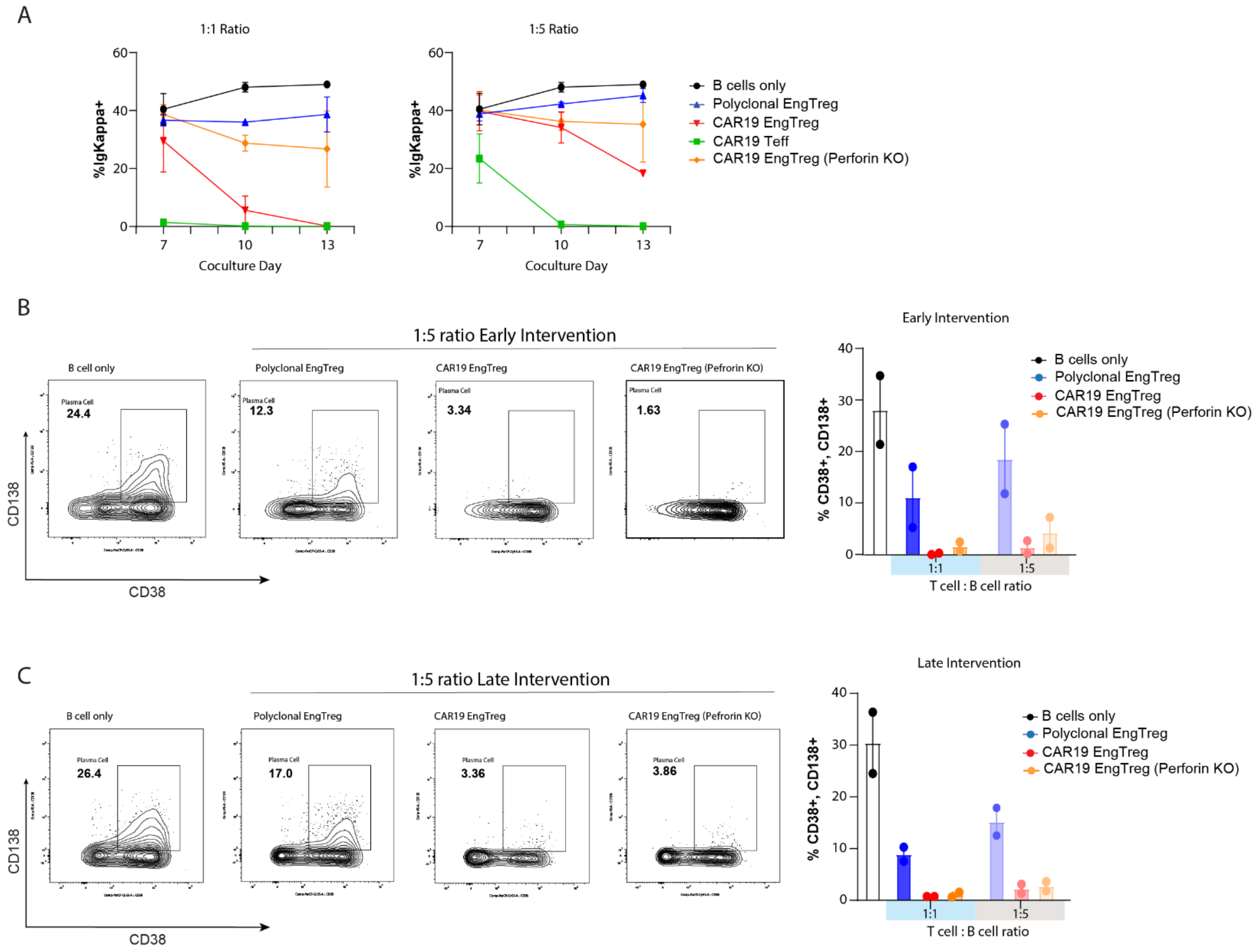


**Fig. S6. PRF-1 KO CAR19 EngTreg suppress PC differentiation in vitro in absence of cytotoxicity.** (**A**) Representative flow cytometry assessing intracellular Ig-Kappa light-chain expression from indicated culture conditions at D7, 10, and 13. (**B**) Flow cytometry analysis of PC formation (based on co-expression of CD38, CD138) on D13 following Early Intervention with indicated T cell products. Representative flow showing data for T:B cell ratio of 1:5. (**C**) PC formation on D13 following Late Intervention with indicated T cell products showing data for T:B cell ratio of 1:5. Data are presented as mean ± SEM and represent two independent experiments from two separate donors.

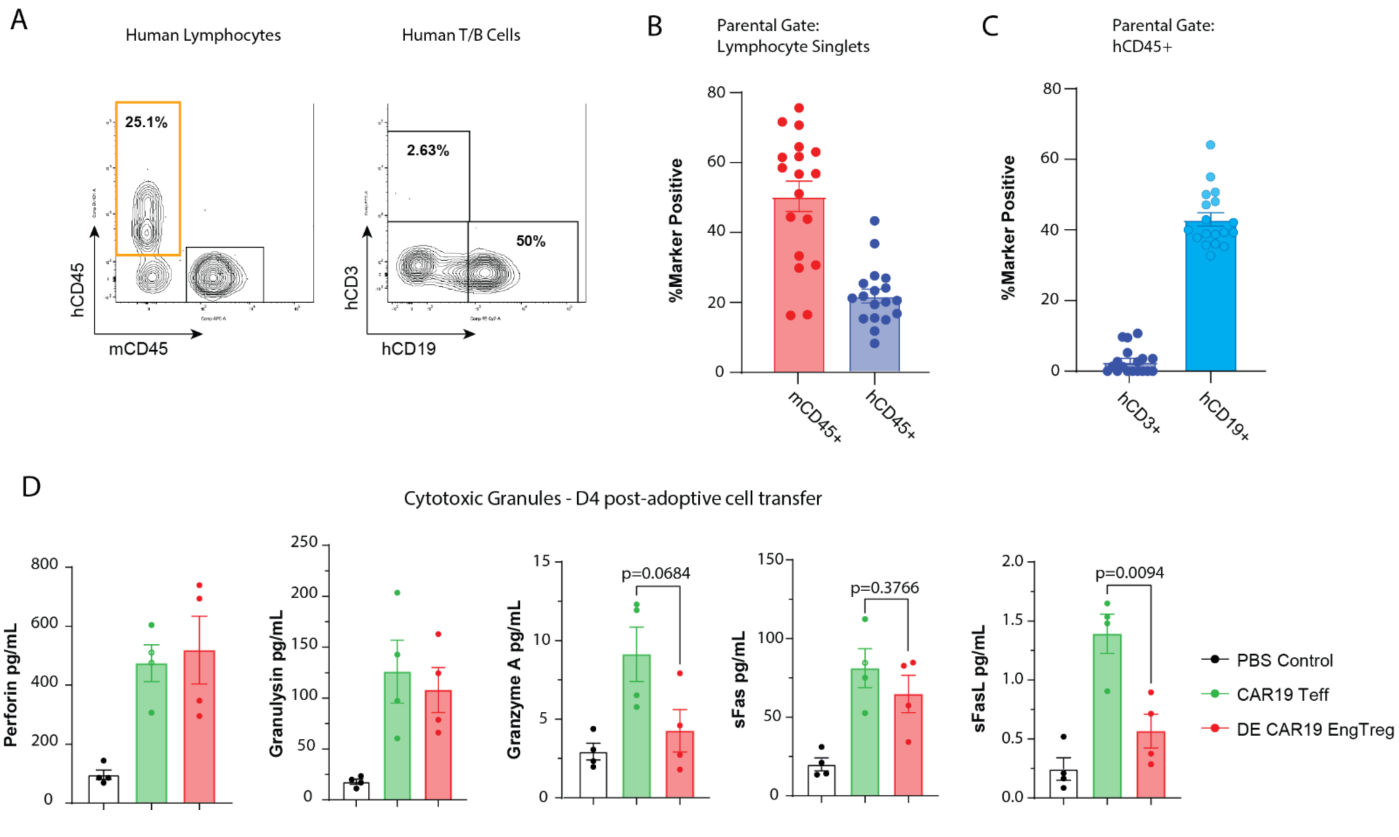


**Fig. S7. Human hematopoietic cell chimerism in NBSGW recipients prior to adoptive transfer of autologous T cell products**. (**A**) Representative flow cytometry showing gating strategy to assess human hematopoietic cell chimerism in PB at 14 weeks post-HSPC transplantation. Left plot shows hCD45$^{+}$ vs mCD45$^{+}$ cells within the lymphocyte singlet gate. Right plot shows proportion of hCD3$^{+}$ T cells and hCD19$^{+}$ B cells within the hCD45 gate (red gate on left plot). (**B**) Graph showing the proportion of human vs. mouse CD45$^{+}$ lymphocytes across all recipient mice. (**C**) Graph showing proportion of human CD3$^{+}$ and human CD19$^{+}$ cells within the hCD45$^{+}$ lymphocyte gate. (**D**) Cytometric Bead Array (CBA) assessment of cytotoxic mediator levels in serum at endpoint. For B-D, data shown as mean +/- SEM, p values calculated by unpaired T-test.

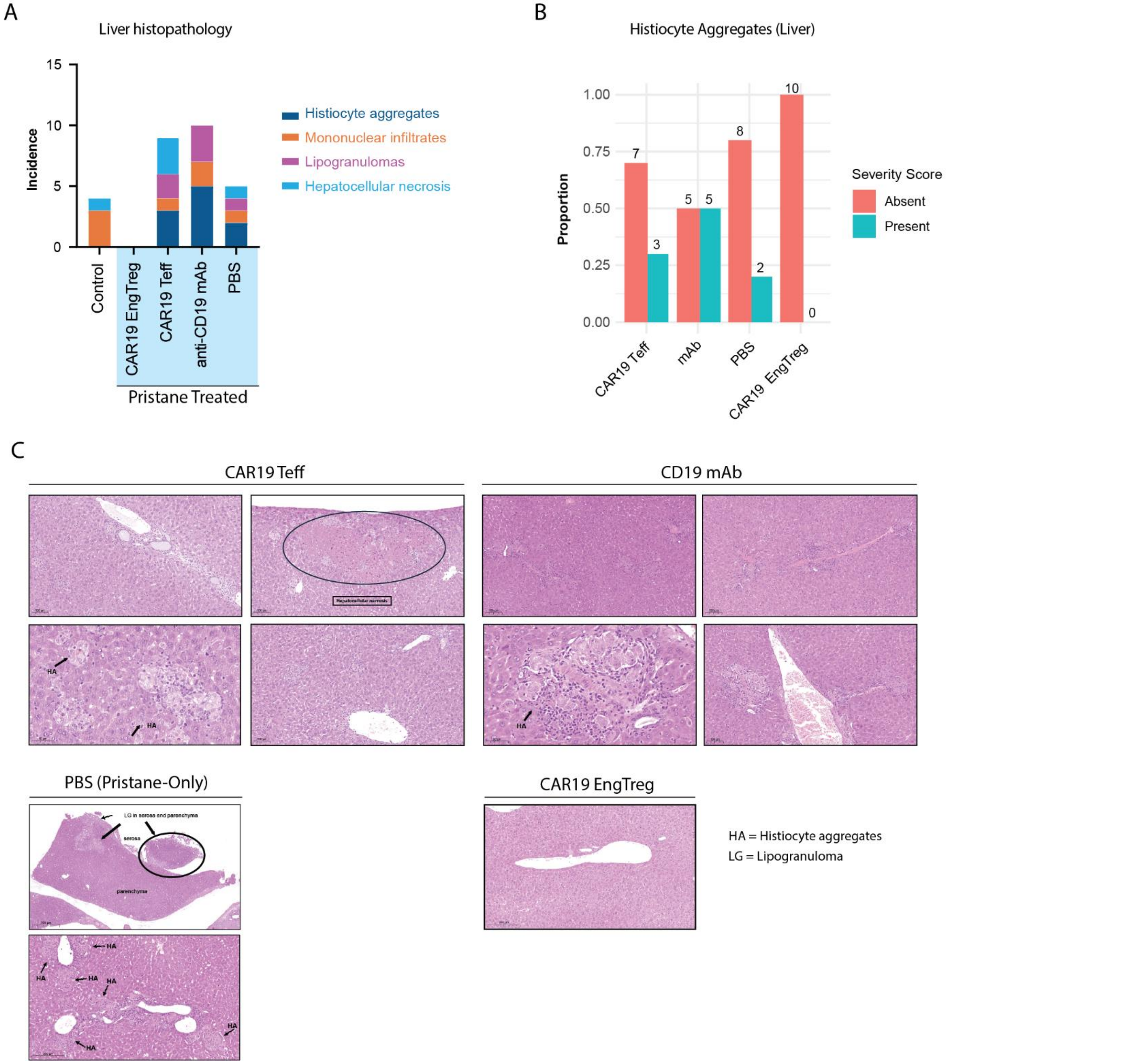


**Fig. S8. Liver histopathology in pristane-induced inflammatory model in humanized mice**. (**A**) Frequency of inflammatory lesions based upon histopathological evaluation in liver. Graphs show the combined proportion of indicated histological findings within each treatment group (CAR19 Teff, CD19 mAb, PBS (pristane-only) and CAR19 EngTreg). (**B**) Quantitation of samples scored as positive (present) or negative (absent) for histiocyte aggregates or mononuclear infiltrates. (**C**) Annotated histology images in livers from indicated treatment groups. HA=Histiocyte aggregates. LG=Lipogranuloma.

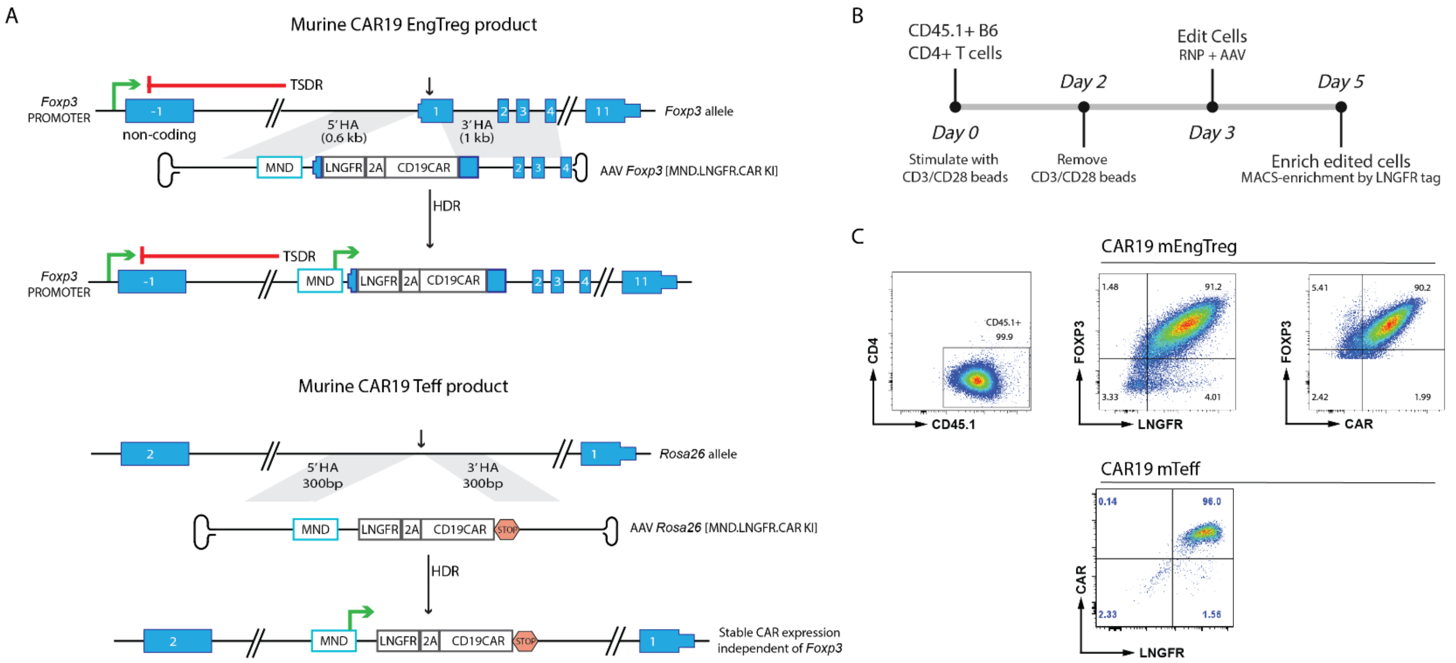


**Fig. S9. Production of murine (m)CAR19 EngTreg and CAR19 Teff for use in adoptive transfer studies**. (**A**) Schematic of HDR editing strategy to generate CAR19 EngTreg and CAR19 Teff from mouse CD4+ T cells. (**B**) Production and enrichment timeline for mouse T cell products. (**C**) Representative flow characterization post-enrichment for CAR19 mEngTreg (*upper panels*) and CAR19 mTeff (*lower panel*).

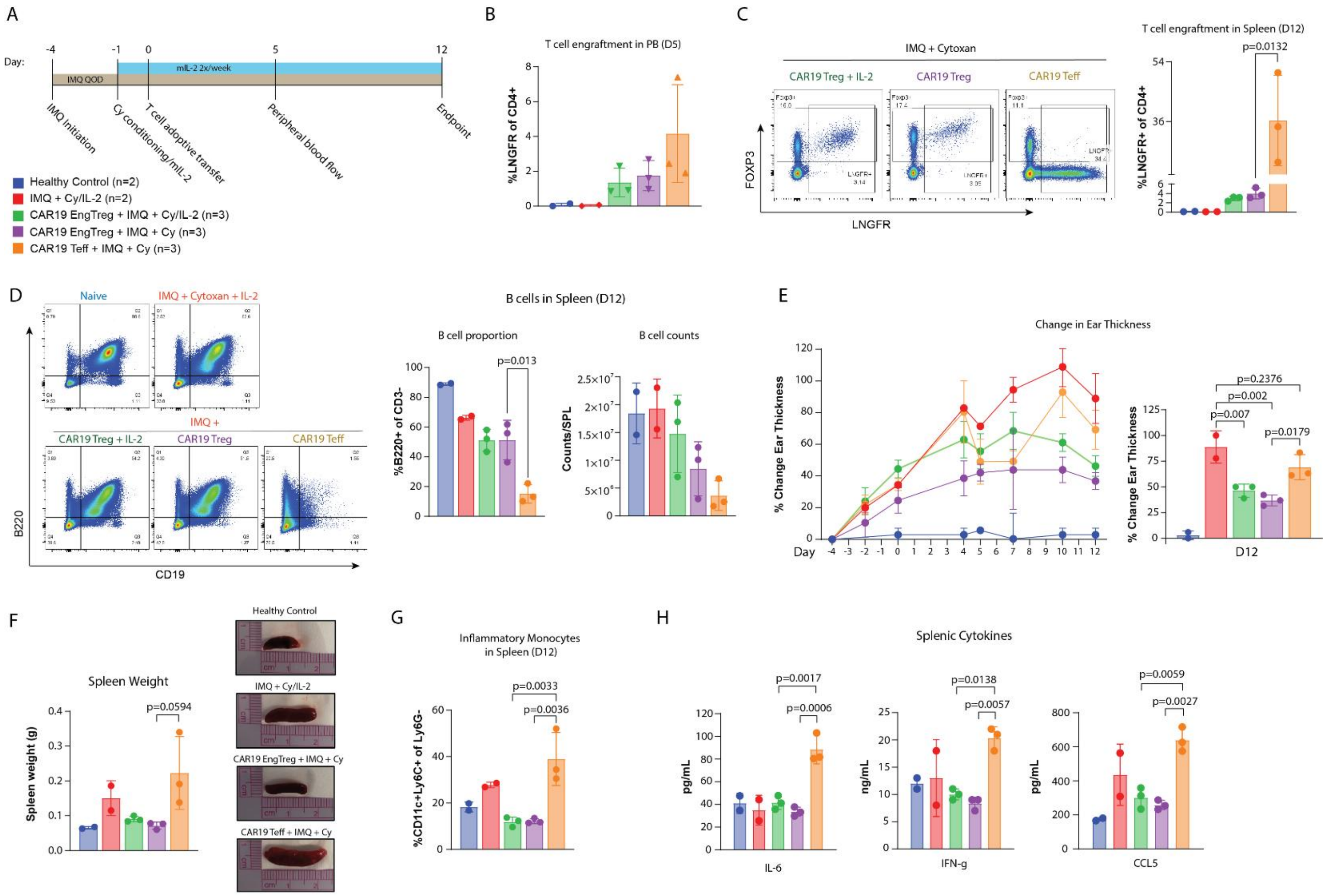


**Fig. S10. mCAR19 EngTreg reduce inflammation in an imiquimod (IMQ)-induced inflammatory model.** (**A**) Schematic and timeline of imiquimod (IMQ) mouse study. (**B**) Engraftment of T cell products in PB at D5 post-adoptive transfer. (**C**) Engraftment of T cell products in spleen at endpoint. (*Left panel*) Representative flow cytometry showing Foxp3/LNGFR expression in CD4+ T cells. (*Right panel*) Proportion of LNGFR+ cells in indicated groups. (**D**) B cells in spleen at endpoint. (*Left panel*) Representative flow cytometry for $B220^{+}/CD19^{+}$ B cells (within of $CD3^{-}$ cell gate). (*Right panel*) Proportion and absolute counts of $B220^{+}$ B cells. (**E**) (*Left panel*) % change in ear thickness over time following IMQ treatment. (*Right panel*) % change at endpoint. (**F**) Spleen size at endpoint. (*Left panel*) Spleen weight and (*right panel*) representative spleen images with ruler for scale. (**G**) Proportion of CD11c+, Ly6C+, Ly6G- inflammatory monocytes in spleen at endpoint. (**H**) Cytometric Bead Array (CBA) measurement of inflammatory cytokines in culture media from stimulated splenocytes isolated at endpoint from each group. For B-H, data shown as mean +/- SEM, p value based on ordinary one-way ANOVA.

A

B

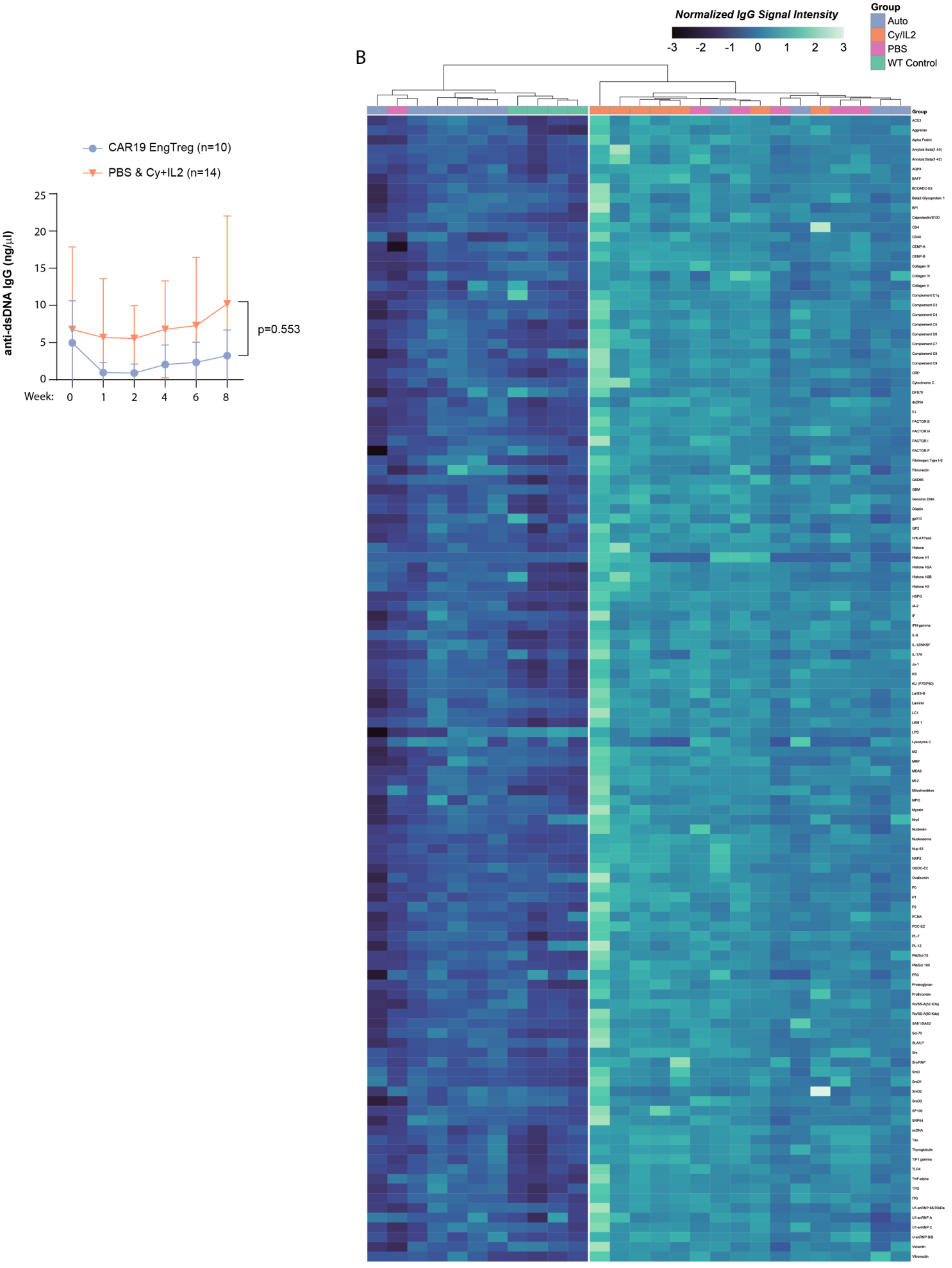

CAR19 EngTreg (n=10)
PBS & Cy+IL2 (n=14)
anti-dsDNA IgG (ng/μl)
25
20
15
10
5
0
Week:
0
1
2
4
6
8
p=0.553
Normalized IgG Signal Intensity
-3
-2
-1
0
1
2
3
Group
Auto
Cy/IL2
PBS
WT Control
Group
ACE2
Aggrecan
Alpha Fodrin
Amyloid Beta(1-40)
Amyloid Beta(1-42)
AQP4
BAFF
BCOADC-E2
Beta2-Glycoprotein 1
BPI
Calprotectin/S100
CD4
CD40
CENP-A
CENP-B
Collagen III
Collagen IV
Collagen V
Complement C1q
Complement C3
Complement C4
Complement C5
Complement C6
Complement C7
Complement C8
Complement C9
CRP
Cytochrome C
DFS70
dsDNA
EJ
FACTOR B
FACTOR H
FACTOR I
FACTOR P
Fibrinogen Type I-S
Fibronectin
GAD65
GBM
Genomic DNA
Gliadin
gp210
GP2
H/K-ATPase
Histone
Histone H1
Histone H2A
Histone H2B
Histone H3
HSPG
IA-2
IF
IFN-gamma
IL-6
IL-12/NKSF
IL-17A
Jo-1
KS
KU (P70/P80)
La/SS-B
Laminin
LC1
LKM 1
LPS
Lysozyme C
M2
MBP
MDA5
Mi-2
Mitochondrion
MPO
Myosin
Nrp1
Nucleolin
Nucleosome
Nup 62
NXP2
OGDC-E2
Ovalbumin
P0
P1
P2
PCNA
PDC-E2
PL-7
PL-12
PM/Scl-75
PM/Scl 100
PR3
Proteoglycan
Prothrombin
Ro/SS-A(52 kDa)
Ro/SS-A(60 Kda)
SAE1/SAE2
Scl-70
SLA/LP
Sm
Sm/RNP
SmD
SmD1
SmD2
SmD3
SP100
SRP54
ssDNA
Tau
Thyroglobulin
TIF1 gamma
TLR4
TNF-alpha
TPO
tTG
U1-snRNP 68/70kDa
U1-snRNP A
U1-snRNP C
U-snRNP B/B
Vimentin
Vitronectin

**Fig. S11. Autoantibody levels following adoptive transfer of CAR19 mEngTreg in the Sle123 lupus model** (**A**) Serum anti-dsDNA IgG was measured by ELISA at baseline prior to pre-conditioning with cyclophosphamide (week 0) and at the indicated timepoints post adoptive cell transfer. Significance determined by repeated measures two-way ANOVA. (**B**) Heatmap of autoantibody microarray data showing hierarchical clustering of autoantibody reactivity across samples. Each column indicates data from a single animal. Color intensity represents relative log2-normalized signal intensity for each autoantibody, and the group identity of each mouse is indicated by the color scheme shown (group key shown in upper right).

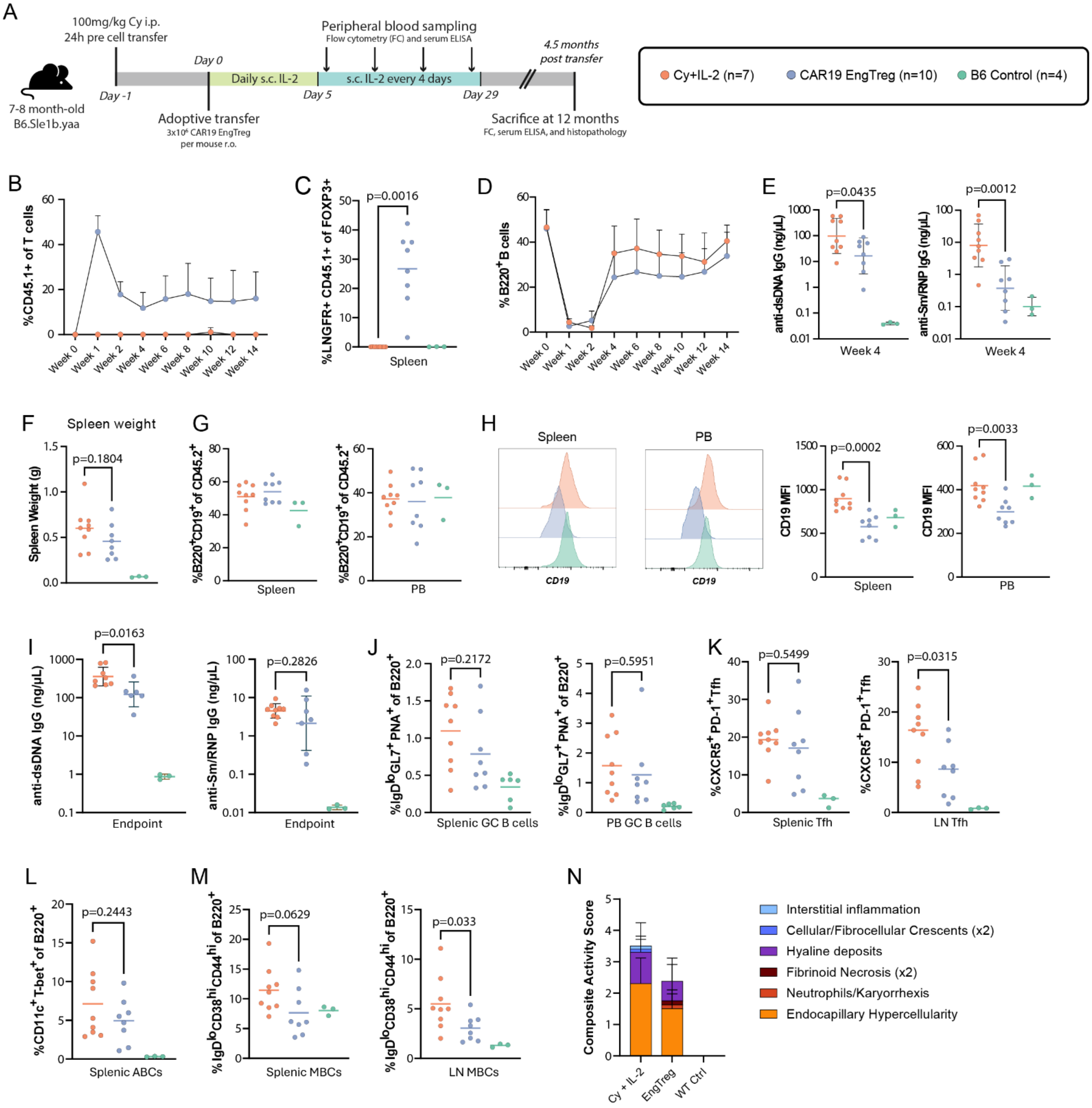


**Fig. S12. mCAR19 EngTreg engraft long-term and modulate pathogenic B cell subsets in the Sle1b.yaa lupus model.** (**A**) Experimental schematic and timeline for Sle1b.yaa mouse model. (**B**) Frequency of CAR19 mEngTreg ($CD45.1^+$ cells within $CD4^+$ T cells) in PB over time post-adoptive transfer. (**C**) Frequency of $CD45.1^+LNGFR^+$ CAR19 mEngTreg expressed as a proportion of total $FOXP3^+$ cells in spleen at endpoint. (**D**) Frequency of $CD19^+$ B cells within $CD45^+$ leukocytes in PB over time. (**E**) Serum dsDNA and Sm/RNP autoantibodies at 4 weeks post-adoptive transfer. (**F**) Spleen weight at endpoint (**G**) Frequency of $CD19^+B220^+$ B cells in spleen (left) and PB (right) at endpoint. (**H**) Representative histograms (left) and combined data (right) showing CD19 MFI on B220+ B cells in spleen and PB at endpoint. (**I**) dsDNA and Sm/RNP autoantibodies at endpoint. (**J**) Frequency of GC B cells (defined as $IgD^{lo}$ $GL7^+$ $PNA^+$) in spleen (left) and LN (right) at endpoint. (**K**) Frequency of $CXCR5^+PD1^+$ T follicular helper (Tfh) cells in spleen (left) and LN at endpoint (right). (**L**) Frequency of $CD11c^+$ $T\text{-}bet^+$ age-associated B cells (ABCs) in spleen at endpoint. (**M**)

Frequency of memory B cells (defined as $IgD^{lo}$ $CD38^{hi}$ $CD44^{hi}$ MBCs) in spleen (left) and LN at endpoint (right). Statistical significance for all comparisons determined by Welch's t-test. Error bars represent mean ± s.d.

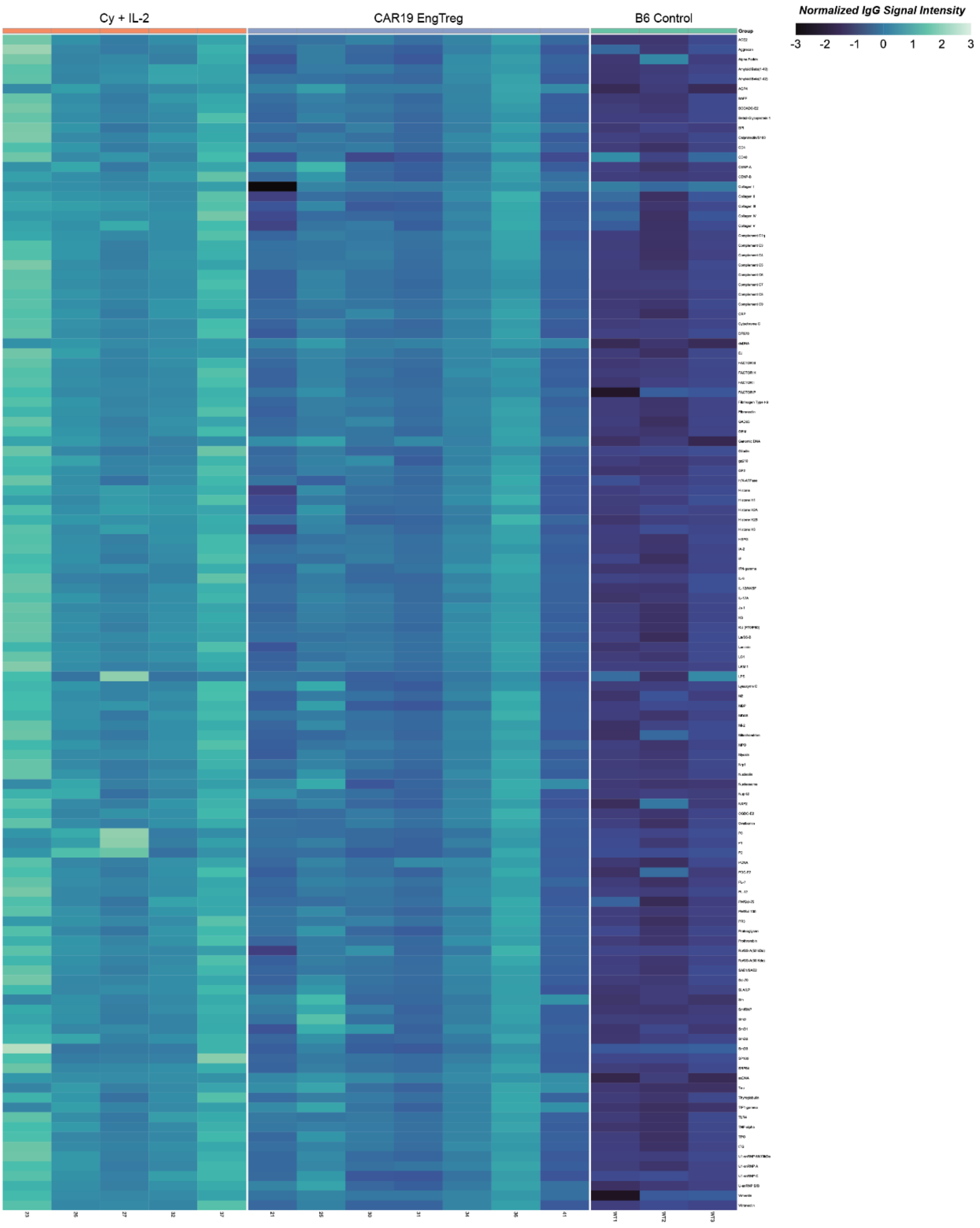
Cy + IL-2
CAR19 EngTreg
B6 Control
Normalized IgG Signal Intensity
-3 -2 -1 0 1 2 3
Group

**Fig. S13. CAR19 mEngTreg reduce broad spectrum autoantibodies in the Sle1b.yaa lupus model.** Serum autoantibody analysis by autoantigen microarray. Heatmap showing log2-normalized IgG signal intensities measured by autoantigen microarray using endpoint serum obtained from indicated treatment groups. Each column shows data from an individual animal and color at top indicates treatment groups, with target antigens indicated on right margin.
